# Tumor cell-specific loss of GPX4 reprograms triacylglycerol metabolism to escape ferroptosis and impair antitumor immunity in NSCLC

**DOI:** 10.1101/2025.11.12.687999

**Authors:** Peng Wang, Shengdan Zhang, Xin Chen, Xu-Dong Yang, Shi Huang, Huiyong Yin, Hao-Yu Duan, Fuling Zhou, Jia Yu, Bo Zhong, Dandan Lin

## Abstract

Glutathione peroxidase 4 (GPX4) is a master regulator of ferroptosis, a process that has been proposed as a potential therapeutic strategy for cancer. Here we have unexpectedly found that inducible knockout of GPX4 in tumor cells significantly promotes non-small cell lung cancer (NSCLC) progression in the autochthonous *Kras*^LSL-G12D/+^*Lkb1*^fl/fl^ (KL) and *Kras*^LSL-G12D/+^*Tp53*^fl/fl^ (KP) mouse models, whereas inducible overexpression of GPX4 in tumor cells exerts the opposite effect. GPX4-deficient tumor cells evade ferroptosis by upregulating the expression of DGAT1/2 to promote the synthesis of triacylglycerol (TAG) and oxidized TAG (oxTAG) and the formation of lipid droplets in cells. In addition, GPX4-deficient tumor cells secrete TAG and oxTAG into the extracellular space to induce dysfunction of antitumor CD8^+^ T cells, thereby coordinating an immunoinhibitory tumor microenvironment (TME). Consistently, treatment with DGAT1/2 inhibitors or inducible overexpression of GPX4 in tumor cells significantly resensitizes tumor cells to ferroptosis and ignites the activation of T cells in the TME to inhibit NSCLC progression. These findings highlight a previously uncharacterized role of tumor cell-specific GPX4 in NSCLC progression and provide potential therapeutic strategies for NSCLC.

## Introduction

Glutathione peroxidase 4 (GPX4) is an enzyme that catalyzes the reduction of cytotoxic phospholipid hydroperoxides (PL-OOH) into PL alcohols (PL-OH) and thereby protects against ferroptosis^1,2^. Deletion of GPX4 leads to the accumulation of PL peroxides and induces cell death in *in vitro* cultured cells^3–9^. Accordingly, *Gpx4*^-/-^ mouse embryos die at E7.5 and Cre-ER;*Gpx4*^fl/fl^ mice exhibit lethality after 2 weeks of tamoxifen (Tam) treatment^3,10–12^, whereas the human *GPX4* transgene rescues the lethal phenotype of *Gpx4*^-/-^ mice and inhibits PL peroxide-induced cell death^13^, demonstrating an indispensable role of GPX4 in the survival and development of mice under homeostatic conditions. In RAS^mut^ cancer cells, genetic deletion or inhibition of GPX4 induces PL peroxidation and ferroptosis and inhibits tumor cell growth in mice^14,15^, suggesting a protumor role of GPX4 in xenograft or syngeneic graft models. However, conditional deletion of GPX4 in the pancreas (*Pdx1*-Cre;*Gpx4*^fl/fl^) of mice promotes tumorigenesis in the KRas^G12D^ model^16^, arguing a protective role of GPX4 in the spontaneous RAS^mut^ cancer model. It should be noted that such a strategy of GPX4 depletion might cause cell death or tissue injury at the early stage of cancer development or even before the malignant transformation of tumor cells, which might have an impact on the functions of GPX4 in tumor progression *in vivo*.

The tumor microenvironment (TME), which includes immune and non-immune cells, metabolites, and cytokines, critically regulates the progression and drug response of cancers^17–19^. Lung cancer is the leading cause of cancer-related mortality^20,21^, and approximately 85% of the diagnosed lung cancers are non-small cell lung cancer (NSCLC)^17^. Various mutations, including those in the tumor-driving genes *KRAS*, *EGFR,* and *ALK* and the tumor suppressor genes *TP53* and *LKB1*, have been identified in NSCLC^22–25^. Studies with genetically engineered mouse models (GEMMs) harboring the mutations (e.g., *Kras*^LSL−G12D/+^*Tp53*^fl/fl^, KP; *Kras*^LSL−G12D/+^*Lkb1*^fl/fl^, KL) have been conducted to elucidate the mechanisms of and screen effective therapies for NSCLC^26–28^. For example, we have found that the chemokine CCL7 secreted by alveolar macrophages in the TME recruits type 1 conventional dendritic cells to promote an antitumor T-cell response and inhibit NSCLC progression^29,30^, whereas neutrophils in the TME produce IL-36γ to alleviate the oxidative stress and promote NSCLC progression by increasing glutathione metabolism in the KL and KP models^31^. In addition, antioxidants such as N-acetylcysteine and vitamin E accelerate KRas^G12D^-driven NSCLC progression by tuning down the global oxidative stress in the TME^32^. Considering the essential role of GPX4 in the reduction of lipid peroxides that contribute to the oxidative stress of cancer cells, whether and how the regulation of lipid peroxidation by GPX4 in tumor cells modulates NSCLC progression in the autochthonous mouse models remain to be investigated.

Here, we have adopted a dual recombinase-mediated gene mutation system (Cre-loxP and DreER^T2^-rox) and a Cre recombinase plus doxycycline (Dox)-rTTA system to specifically and inducibly delete and overexpress GPX4 in tumor cells in the autochthonous KL and KP NSCLC mouse models, respectively. Interestingly, inducible knockout of GPX4 in the tumor cells of the established autochthonous KL or KP tumors significantly promotes the progression of NSCLC and accelerates the death of mice, and inducible overexpression of GPX4 in tumor cells has the opposite effect. Mechanistically, inducible knockout of GPX4 in tumor cells leads to the accumulation of TAG and oxidized TAG (oxTAG) in tumor cells and the formation of lipid droplets in a DGAT1/2-dependent manner, thereby protecting tumor cells from ferroptosis. In addition, inducible knockout of GPX4 in tumor cells promotes the secretion of TAG and oxTAG, which induces dysfunction and exhaustion of antitumor CD8^+^ T cells in the TME. Selective overexpression of GPX4 in tumor cells or pharmacological inhibition of DGAT1/2 impairs the synthesis of TAG and oxTAG, promotes the generation of oxPE/PC in tumor cells, and enhances antitumor immunity to inhibit NSCLC progression in autochthonous mouse models. These findings have demonstrated an unexpected role of tumor cell-specific GPX4 in reprogramming TAG metabolism and antitumor immunity in the TME and the progression of autochthonous NSCLC.

## Results

### Inducible knockout of GPX4 in tumor cells promotes NSCLC progression in autochthonous KP and KL mouse models

To investigate the role of GPX4 in tumor cells in autochthonous KP and KL non-small cell lung cancer (NSCLC) models, we took advantage of the Cre-loxP and the DreER^T2^-rox systems to inducibly delete GPX4 in tumor cells of the established KP and KL tumors.

Specifically, the targeting vector consisting of rox-*Gpx4*-rox-DreER^T2^-loxP2272-STOP-loxP2272-CAG promoter and the flanking homologous sequences was knocked into the *Gpx4* gene locus to generate *Gpx4*-rox-CAG-LSL-DreER^T2^ mice (termed *Gpx4*^m/m^ hereafter) (Figure S1A and S1B). Cre-mediated removal of the STOP cassette allowed the expression of the DreER^T2^ protein that would induce the knockout of GPX4 only after 4-hydroxytamoxifen (4-OHT) treatment (Figure S1C). We next intranasally infected the *Kras*^LSL−G12D/+^*Lkb1*^fl/fl^*Gpx4*^m/m^ (KLG4^m/m^) mice with Ad-Cre, and 5 weeks later (when KRas^G12D^ induced malignant transformation and when the tumor lesions were formed)^27,30,31,33^, the mice were intraperitoneally injected with corn oil or tamoxifen (Tam) (which was metabolized into 4OHT in mice) every other day for 2 weeks. The mice were then euthanized for various analyses (Figure S2A, the autochthonous NSCLC models in this study were induced in this way otherwise indicated). PCR analysis of genomic DNA from different tissues suggested that exons 5-7 of the *Gpx4* gene were specifically deleted in tumors but not in lung non-cancerous tumor-adjacent tissue (NAT), heart, liver, spleen, kidney, or brain or in the organs of corn oil-treated KLG4^m/m^ counterparts (Figure S2B). We further FACS-sorted different types of cells in the KL and the KLG4^m/m^ tumors (Figure S2C), and found that the rearrangement of the *Gpx4* gene selectively occurred in tumor cells (EpCAM^+^CD31^-^) (Figure S2D). Additionally, the protein levels of GPX4 were substantially decreased in EpCAM^+^CD31^-^ tumor cells but not in endothelial cells (CD31^+^CD49d^-/int^), stromal cells (CD31^-^CD49d^+^), or immune cells (CD45^+^) in KLG4^m/m^ tumors compared to the KL counterparts (Figure S2E), suggesting efficient, specific and inducible deletion of GPX4 in tumor cells in the KLG4^m/m^ autochthonous NSCLC mouse model.

We next examined the effect of inducible knockout of GPX4 in tumor cells on the progression of autochthonous murine NSCLC. Interestingly, we found that the survival of KLG4^m/m^ mice was significantly shorter than that of KL mice after Ad-Cre infection and Tam treatment (Figure 1A). In addition, the tumor burdens and the individual tumor sizes in the lungs of KLG4^m/m^ mice were significantly greater than those in the lungs of KL mice, as suggested by micro-computed tomography (Micro-CT) imaging and hematoxylin and eosin (HE) staining analyses (Figure 1B and 1C). With similar treatments, the *Kras*^LSL−G12D/+^*Tp53*^fl/fl^*Gpx4*^m/m^ (KPG4^m/m^) mice exhibited shorter survival time, larger tumor sizes, and heavier tumor burdens than the KP mice (Figure 1D-1F). In addition, the KLG4^m/m^ mice had larger tumor sizes and heavier tumor burdens in the lungs than did the KL mice after intranasal injection of Ad-SPC-Cre (which has been shown to specifically and efficiently express the Cre recombinase in lung alveolar type II epithelial cells^34–36^) followed by Tam treatment (Figure S3A and S3B). These data demonstrate that inducible knockout of GPX4 in tumor cells promotes autochthonous NSCLC progression in the KL and KP mouse models.

**Figure 1.**
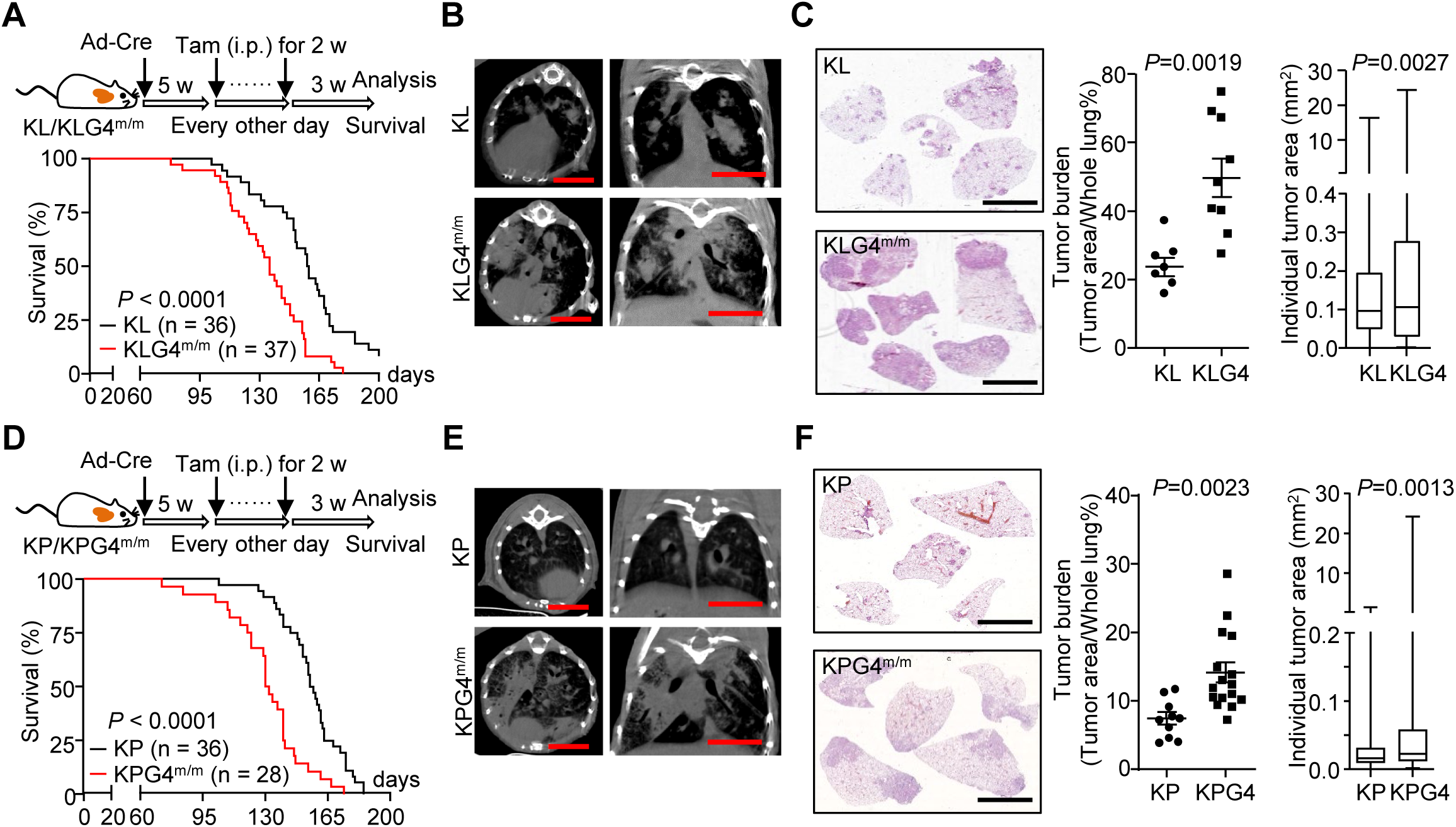
Inducible knockout of GPX4 in tumor cells promotes NSCLC progression in the autochthonous KP and KL mouse models. (A) A scheme of tumor induction (upper) and the survival (lower) of *Kras*^LSL-G12D/+^*Lkb1*^fl/fl^ (KL, n = 36) and *Kras*^LSL-G12D/+^*Lkb1*^fl/fl^*Gpx4*^m/m^ (KLG4^m/m^, n = 37) mice. The KL and the KLG4^m/m^ mice were intranasally injected with Ad-Cre (2×10^6^ PFU per mouse) for 5 weeks followed by intraperitoneal injection of tamoxifen every other day for 2 weeks. The mice were allowed to rest for another 3 weeks followed by various analyses or monitored for survival. (B, C) Representative images of micro-CT (B) and HE staining (C, left) and statistics of tumor burdens (C, middle) and individual tumor sizes (C, right) of tumor-burdened lungs from the KL (n = 7) and KLG4^m/m^ (n = 9) mice treated as in (A). (D) A scheme of tumor induction (upper) and the survival (lower) of *Kras*^LSL-G12D/+^*Tp53*^fl/fl^ (KP, n = 36) and *Kras*^LSL-G12D/+^*Tp53*^fl/fl^*Gpx4*^m/m^ (KPG4^m/m^, n = 28) mice that were treated as described in (A). (E, F) Representative images of micro-CT (E) and HE staining (F, left), and statistics of tumor burdens (F, middle) and individual tumor sizes (F, right) of tumor-burdened lungs from KP (n = 10) and KPG4^m/m^ (n = 16) mice treated as in (D). Statistical analyses were performed with Log-Rank analysis (A, D) or two-tailed student’s *t*-test (C, F). Graphs show mean ± SEM (C, F). Scale bars represent 5 mm (B, C, E, F). Data are combined results of three independent experiments (A, D) or representative results of two independent experiments (B, C, E, F). \**P* < 0.05, \*\**P* < 0.01, \*\*\**P* < 0.001, \*\*\*\**P* < 0.0001.

### Inducible knockout of GPX4 inhibits tumor growth in syngeneic graft mouse models

Previous studies have shown that knockdown or inhibition of GPX4 in RAS^mut^ tumor cell lines induces cancer cell ferroptosis and inhibits tumor growth^14,15^. We next examined the role of GPX4 in tumor cell growth in syngeneic graft models. The CD45^-^CD31^-^EpCAM^+^ tumor cells from KL and KLG4^m/m^ tumors were subcutaneously inoculated into the flanks of wild-type C57BL/6 mice followed by various analyses (Figure S3C). The results suggested that GPX4 was efficiently deleted in the KLG4^m/m^ tumors and that the growth of KLG4^m/m^ tumor cells was severely compromised compared to the KL tumor cells (Figure S3C and S3D), suggesting that GPX4 is indispensable for tumor cell growth in the subcutaneous syngeneic graft model.

To examine the role of GPX4 in maintaining tumor growth, we isolated the CD45^-^CD31^-^EpCAM^+^ tumor cells from KL and KLG4^m/m^ mice that were intranasally infected with Ad-Cre for 10 weeks and subcutaneously inoculated the cells into the flanks of wild-type C57BL/6 mice followed by Tam treatment (Figure S3E). The results suggested that the growth of KLG4^m/m^ tumors was similar to that of KL tumors before Tam treatment, whereas the growth of KLG4^m/m^ tumors but not KL tumors was significantly inhibited after Tam treatment, which was accompanied by a decrease of GPX4 in KLG4^m/m^ tumors (Figure S3E-S3G). These data demonstrate that GPX4 is required for tumor cell growth and maintenance in subcutaneous syngeneic graft models.

### Liprostatin-1 restores syngeneic KLG4^m/m^ tumor growth

Knockout or inhibition of GPX4 promotes lipid peroxidation and ferroptosis in *in vitro* cell cultures, which is rescued by lipophilic radical-trapping agents such as liprostatin-1 (Lip-1) or α-tocopherol^3,12,15^. Consistent with this notion, the tumor cells from Tam-treated syngeneic KLG4^m/m^ tumors exhibited higher lipid peroxidation and more severe ferroptotic death than did those from Tam-treated syngeneic KL tumors, as indicated by C11-BODIPY and SYTOX staining (Figure S3H). Treatment with Lip-1 substantially decreased lipid peroxidation and ferroptotic cell death of KLG4^m/m^ tumor cells and restored the growth of KLG4^m/m^ tumors that should have shrunk after Tam treatment (Figure S3E-S3H). These data suggest that inducible knockout of GPX4 results in ferroptosis of KL tumor cells in syngeneic subcutaneous models.

We next examined lipid peroxidation in the tumor cells of the autochthonous KL and KLG4^m/m^ tumors and found that the C11-BODIPY staining in CD45^-^CD31^-^EpCAM^+^ tumor cells from autochthonous KLG4^m/m^ tumors was significantly higher than in those from KL tumors (Figure S4A). Interestingly, however, we did not observe increased death or mitochondrial damage in KLG4^m/m^ CD45^-^CD31^-^EpCAM^+^ tumor cells compared to the KL counterparts as revealed by the SYTOX staining and transmission electron microscopy (TEM) analyses (Figure S4B and S4C). This phenomenon was further confirmed by immunofluorescence staining of 4-hydroxynonenal (4-HNE, a downstream metabolite of PL peroxidation, which is commonly used as a surrogate marker of ferroptosis), and terminal deoxynucleotidyl transferase (TdT)-mediated dUTP nick end labeling (TUNEL) within the EpCAM^+^ cells in KL and KLG4^m/m^ tumors (Figure S4D). Lipidomic analysis revealed that the levels of classical ferroptosis-associated oxygenated phosphatidylcholine (PC) (PC 18:0/22:4[2O] and PC 18:0/20:4[2O]) and phosphatidylethanolamine (PE) (PE 18:0/20:4[2O] and PE 18:0a_HETE), polyunsaturated fatty acid (PUFA)-PC and PUFA-PE and free fatty acid were comparable between the KL and the KLG4^m/m^ tumor cells^12,37–39^ (Figure S4E-S4G and Table S1A), indicating that inducible GPX4 deficiency in tumor cells does not lead to the accumulation of ferroptotic PL hydroperoxides in the autochthonous NSCLC models.

### The autochthonous *Gpx4*-null tumor cells adapt to triacylglycerol synthesis and lipid droplet formation

Further analysis of our lipidomic data suggested that triacylglycerol (TAG) (especially PUFA-TAG) was increased and diacylglycerol (DAG) was decreased in KLG4^m/m^ tumor cells compared to KL tumor cells (Figure 2A-2C and Table S1B). In addition, the results from targeted lipidomics assays suggested that oxidized TAG (oxTAG) was significantly increased in KLG4^m/m^ tumor cells compared to KL tumor cells (Figure 2D and Table S1A). Consistent with the notion that cells convert excessive lipids (mainly TAG) into lipid droplets^40,41^, we observed increased Oil Red O staining in KLG4^m/m^ tumor cells compared to KL tumor cells (Figure 2E). In addition, the results from the ultrastructural section and TEM analysis revealed more and larger lipid droplets in KLG4^m/m^ tumor cells than in KL tumor cells (Figure 2F). Notably, the lipid droplets contained more oxidized lipids in KLG4^m/m^ tumor cells than in KL tumor cells, as indicated by the C11-BODIOY staining (Figure 2G), which is consistent with the observations that more oxTAG was identified in KLG4^m/m^ tumor cells than in KL tumor cells (Figure 2D). Similarly, we also observed a higher abundance of lipid droplets containing oxidized lipid species in KPG4^m/m^ tumor cells than in KP tumor cells (Figure 2H). Collectively, these data implicate that inducible deletion of GPX4 in tumor cells in the autochthonous NSCLC models leads to the synthesis and storage of TAG and oxTAG to evade ferroptosis. In support of this notion, it has been shown that lipid droplets protect the Drosophila glial cell niche and neural stem cells from harmful PUFA oxidation-induced ferroptosis^42^.

**Figure 2.**
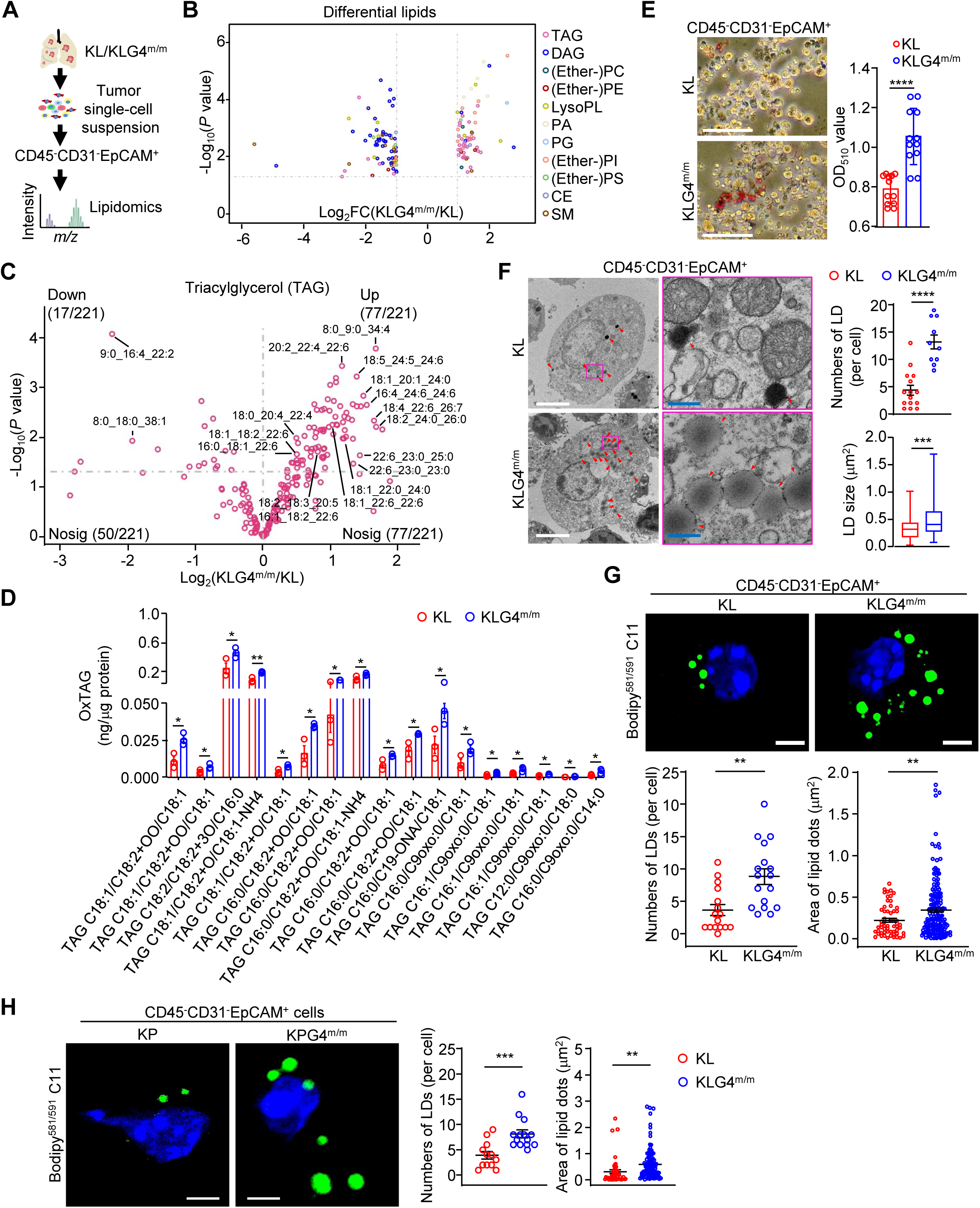
Inducible knockout of GPX4 in tumor cells increases the TAG accumulation and the lipid droplet formation in the autochthonous NSCLC models. (A) A workflow scheme of lipidomics analyses of KL and KLG4^m/m^ CD45^-^CD31^-^EpCAM^+^ tumor cells. (B) Volcano plot of differential lipids (*P* value < 0.05 and |Log_2_[fold change(FC)]| >= 1) in KLG4^m/m^ versus KL CD45^-^CD31^-^EpCAM^+^ tumor cells. (C) Volcano plot showing the levels of triacylglycerols (TAGs) in CD45^-^CD31^-^EpCAM^+^ tumor cells derived from KLG4^m/m^ mice (n = 3) versus KL-derived tumor cells (n = 3). (D) Quantitative assessment of oxidized triacylglycerols (oxTAGs) in CD45^-^CD31^-^EpCAM^+^ tumor cells from lung tumors of KL (n = 3) and KLG4^m/m^ (n = 3) mice. (E) Representative images (left) and statistical analysis (right) of Oil Red O staining of CD45^-^CD31^-^EpCAM^+^ tumor cells from KL (n = 12) and KLG4^m/m^ (n = 12) lung tumors. (F) Representative images from transmission electron microscopy (left, red arrowheads indicate lipid droplets) along with statistics of the numbers and sizes (right) of lipid droplets (LDs) in CD45^-^CD31^-^EpCAM^+^ tumor cells from KL (n = 15) and KLG4^m/m^ (n = 10) mice. The magenta boxed areas are shown at higher magnification on the right. (G) Representative images (upper images) and statistical analysis (lower graphs) of oxidized lipid-containing LDs in KL (n = 16) and KLG4^m/m^ (n = 17) CD45^-^CD31^-^EpCAM^+^ tumor cells. (H) Representative images (upper images) and statistical analysis (lower graphs) of oxidized lipid-containing LDs in KP (n = 12) and KPG4^m/m^ (n = 14) CD45^-^CD31^-^EpCAM^+^ tumor cells. Graphs show mean ± SEM (D-H). Statistical analyses were performed with multiple *t*-test (D) and two-tailed student’s *t*-test (E-H). Scale bars represent 100 μm (E), 5 μm (white, F, G and H) and 500 nm (blue, F). N = 3 mice per group (B-D). Data are representative of three independent experiments (E-H). * *P* < 0.05, ** *P* < 0.01, *** *P* < 0.001, **** *P* <0.0001.

To investigate the mechanism underlying the increase in TAG synthesis and storage in *Gpx4*-null tumor cells, we performed transcriptome sequencing (mRNA-seq) of KL and KLG4^m/m^ tumor cells (Figure 3A), and found that the expression of genes involved in TAG synthesis such as *Gpd1l*, *Gpam*, *Agpat4*, *Plpp1*, *Dgat2* and *Srebf2*^41,43,44^ was significantly upregulated in KLG4^m/m^ tumor cells compared to KL tumor cells (Figure 3B-3C and Table S2A), which was further confirmed by RT-qPCR assays (Figure S5A and S5B), indicating that GPX4 reprograms the expression of genes involved in TAG metabolism. Consistent with this notion, analysis of scRNA-seq data suggested that tumor cells expressing high levels of *GPX4* mRNA (*GPX4*^high^) exhibited lower levels of *DGAT2*, *GPD1L*, and *APOE* mRNA than those expressing low levels of *GPX4* mRNA (*GPX4*^low^) in the human NSCLC tissues or the KL NSCLC mouse model^31,45^ (Figure S5C and S5D). Interestingly, however, the expression of *Dgat2* and *Apoe* became comparable between KLG4^m/m^ and KL tumor cells at 12 h after culture and was lower in KLG4^m/m^ tumor cells than in KL tumor cells at 24 h after culture (Figure S5E). In addition, the expression of *Dgat2* and *Apoe* was not upregulated in subcutaneous syngeneic KLG4^m/m^ tumors compared to KL tumors (Figure S5F and S5G), indicating that the microenvironment in the autochthonous NSCLC models licenses GPX4-mediated regulation of genes involved in TAG synthesis. These data suggest that the autochthonous *Gpx4*-null tumor cells upregulate the expression of *Dgat2* and adapt to triacylglycerol synthesis and lipid droplet formation.

**Figure 3.**
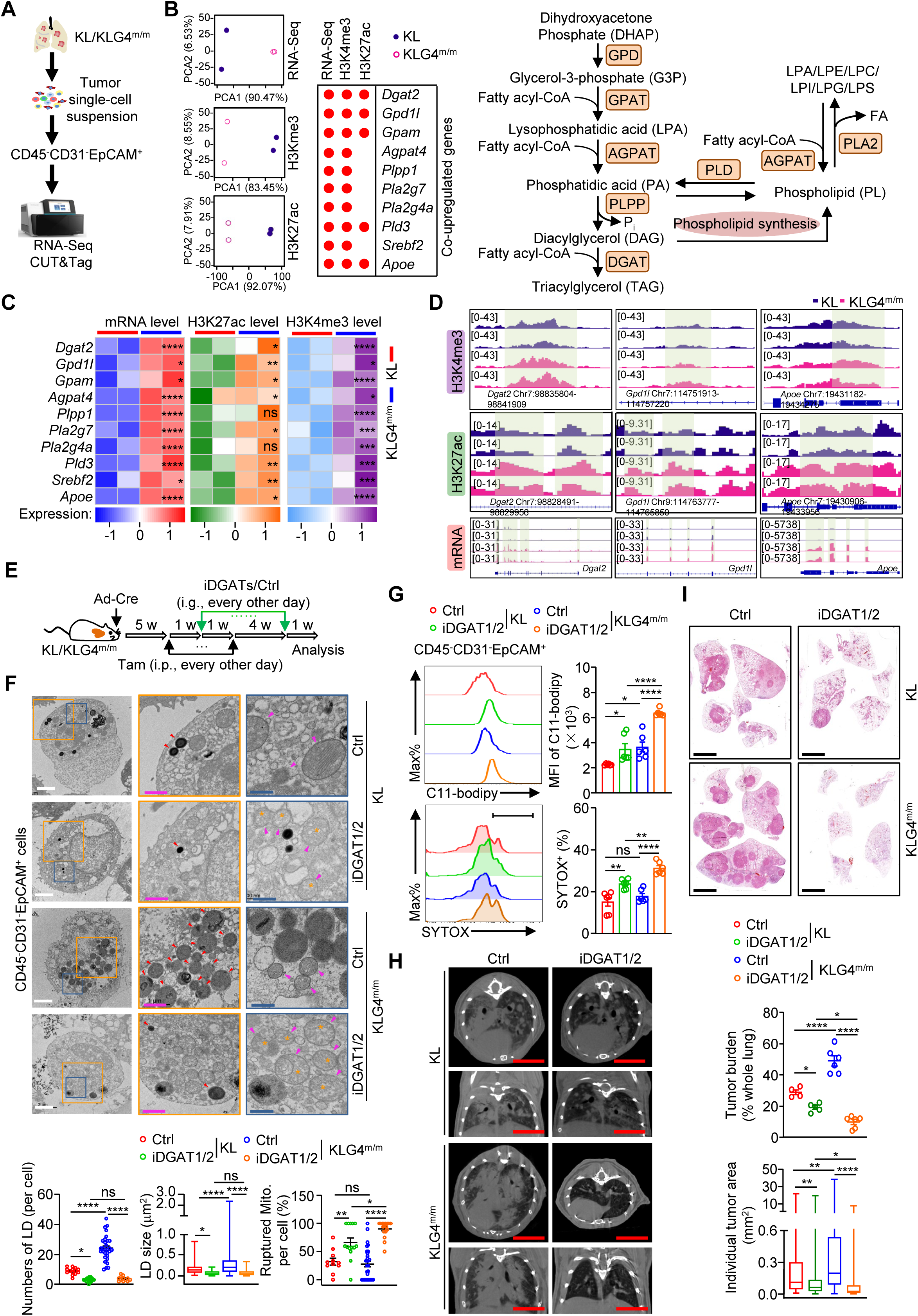
Inhibition of TAG synthesis re-sensitizes *Gpx4*-deficient tumor cells to ferroptosis. (A) A workflow scheme of RNA-seq and CUT&Tag (H3K4me3 and H3K27ac) of KL and KLG4^m/m^ CD45^-^CD31^-^EpCAM^+^ tumor cells. (B) The principal component analysis (PCA) plot showing the differentially expressed genes (DEGs) in RNA-Seq and the differential histone modifications in CUT&Tag assays of KL and KLG4^m/m^ CD45^-^CD31^-^EpCAM^+^ tumor cells (left). Dotplot showing the upregulated DEGs and the histone modifications in CUT&Tag assays of KL and KLG4^m/m^ CD45^-^CD31^-^EpCAM^+^ tumor cells (middle). Scheme of TAG and phospholipids synthesis pathway (right). GPD, glycerol-3-phosphate dehydrogenase; GPAT, glycerol-3-phosphate acyltransferase; AGPAT, 1-acylglycerol-3-phosphate O-acyltransferase; PLPP, phospholipid phosphatase; DGAT, diacylglycerol acyltransferase; PLD, Phospholipase D; PLA2, Phospholipase A2; FA, fatty acid. (C) Heatmap showing the expression levels, H3K4me3 and H3K27ac modifications of the 10 TAG synthesis-related DEGs. (D) Representative Integrative Genomics Viewer (IGV) displaying CUT&Tag sequencing and RNA-sequencing signal profiles across *Dgat2*, *Gpd1l* and *Apoe* loci in CD45^-^CD31^-^EpCAM^+^tumor cells derived from lung tumors of KL and KLG4^m/m^ mice. (E) Schematic illustration depicts tumor induction and iDGAT1/2 (composed of T863 and PF06424439) treatment in KL and KLG4^m/m^ mice that were intranasally injected with Ad-Cre (2×10^6^ PFU per mouse) for 5 weeks followed by intraperitoneal injection of tamoxifen every other day for 2 weeks. One week after Tam treatment, the mice were injected with iDGAT1/2 (composed of T863 and PF06424439, 20 mg and 40 mg per kg body weight, respectively) every other day by gavage for 5 weeks. The mice were rested for one week followed by various analyses. (F) Representative images from transmission electron microscopy showing the morphology of lipid droplets (red arrowheads, in middle panel images) and mitochondria (bottom panel images, mitochondria were indicated by magenta arrowheads and ruptured mitochondria were highlighted with orange asterisks) along with statistics of the numbers and the sizes of lipid droplets (LDs) and the percentage of ruptured mitochondria in CD45^-^CD31^-^EpCAM^+^ tumor cells from lung tumors of KL and KLG4^m/m^ mice treated as in (E). The orange and blue boxed areas are shown at higher magnifications in the middle (orange box) and the bottom (blue box), respectively. (G) The levels of C11-bodipy staining (upper panels) and SYTOX staining (lower panels) of CD45^-^CD31^-^EpCAM^+^ tumor cells from lung tumors of KL and KLG4^m/m^ mice treated as in (E). N=6 for each group. (H, I) Representative images of HE staining (H, left), statistics of tumor burdens and individual tumor sizes (H, right) and representative images of micro-CT (I) of tumor-burdened lungs from the KL (n = 4 for control and n = 5 for iDGAT1/2, respectively) and KLG4^m/m^ (n = 6 for control and n = 6 for iDGAT1/2) mice treated as in (E). N=2 mice per group (B-D). Graphs show mean ± SEM (F-H). Statistical analyses were performed with two-way ANOVA (F-H). Scale bars represent 2 μm (white, F), 1 μm (magenta, F), 500 nm (blue, F), and 5 mm (H and I). Data are representative of three (F-I) independent experiments. * *P* < 0.05, ** *P* < 0.01, *** *P* < 0.001, **** *P* <0.0001. ns, not significant.

### Knockout of GPX4 promotes H3K4me3 and K3K27ac modifications on *Dgat2* gene locus in autochthonous tumors

We next performed high-throughput Cleavage Under Targets and Tagmentation (CUT&Tag) sequencing assays (H3Kme3 or H3K27ac) to examine whether the transcriptional upregulation of genes involved in TAG metabolism was associated with epigenetic modifications on the loci of genes. The results suggested that the H3Kme3 and H3K27ac modifications on the locus of the *Dgat2* and *Gpd1l* genes in KLG4^m/m^ tumor cells were significantly increased compared to those in KL tumor cells (Figure 3B-3D and Table S3A and S3B), which was confirmed by ChIP-qPCR assays (Figure S6A and S6B). In contrast, however, we found that the epigenetic markers (H3Kme3 or H3K27ac) on the loci of *Dgat2* and *Gpd1l* were comparable between the syngeneic KL and KLG4^m/m^ tumor cells or even lower in KLG4^m/m^ tumor cells than in KL tumor cells (Figure S6C), consistent with the observations that the levels of *Dgat2* and *Gpd1l* were not upregulated in syngeneic KLG4^m/m^ tumor cells than in KL tumor cells (Figure S5F and S5G). These data together suggest that inducible knockout of GPX4 in autochthonous tumor cells results in upregulation of TAG synthesis-related genes at both the epigenetic and the transcriptional levels.

### Inhibition of TAG synthesis sensitizes the autochthonous *Gpx4*-null tumor cells to ferroptosis

Diacylglycerol acyltransferase 1 and 2 (DGAT1/2) are rate-limiting enzymes in TAG synthesis that catalyze the conversion of DAG into TAG^44,46^. T863 and PF06424439 are inhibitors of DGAT1 and DGAT2 respectively (collectively referred to as iDGAT1/2 hereafter) that substantially inhibit TAG synthesis^43,47^. We further found that the injection of iDGAT1/2 by gavage significantly decreased the number of lipid droplets and promoted mitochondrial rupture in KL and KLG4^m/m^ tumor cells (Figure 3E and 3F and Figure S7A). Consistently, results from lipidomics and targeted lipidomics analyses suggested that TAG (especially PUFA-TAG) and oxTAG were substantially decreased in KL and KLG4^m/m^ tumor cells after iDGAT1/2 treatment (Figure S7B-E and Table S1C). In addition, treatment with iDGAT1/2 increased levels of oxPE and oxPC and C11-BODIPY and SYTOX staining in KL and KLG4^m/m^ tumor cells (Figure 3G, Figure S7E, and Table S1D), indicating that iDGAT1/2 promotes ferroptosis of autochthonous KL and KLG4^m/m^ tumor cells. Consistently, iDGAT1/2 attenuated tumor progression in the autochthonous KL and KLG4^m/m^ models (Figure 3H and 3I). Notably, iDGAT1/2 promoted mitochondrial rupture, lipid oxidation (oxPE and oxPC), and cell death more potently in KLG4^m/m^ tumor cells than in KL tumor cells and inhibited tumor progression more extensively in KLG4^m/m^ mice than in KL mice (Figure 3F-3I and Figure S7A and S7E). Taken together, these data indicate that the inhibition of TAG synthesis by iDGAT1/2 sensitizes KLG4^m/m^ tumor cells to ferroptosis and inhibits NSCLC progression in the autochthonous KL model.

### The autochthonous *Gpx4*-null tumor cells exhibit increased efflux of TAG and oxTAG

It has been recognized that liver cells synthesize very low-density apolipoprotein (APO) as the main carrier of TAG to promote the efflux of TAG^48,49^. Interestingly, we found that *Apoe* was more highly expressed in KLG4^m/m^ and KPG4^m/m^ tumor cells than in KL and KP tumor cells, respectively (Figure 3B-3D and Figure S5A-D), and that a portion of APOE was colocalized with lipid droplets in KLG4^m/m^ and KL tumor cells (Figure 4A). In addition, we observed elevated levels of TAG in the tumor interstitial fluid (TIF) of KLG4^m/m^ or KPG4^m/m^ tumors compared with those in KL or KP tumors, respectively (Figure 4B and Figure S8A), which were compromised by iDGAT1/2 treatment (Figure 4B). The results from ultrastructural section and TEM analyses suggested the efflux of lipid droplets in KLG4^m/m^ and KL tumor cells (Figure S8B), indicating that inducible knockout of GPX4 in tumor cells results in increased expression of APOE and efflux of TAG.

**Figure 4.**
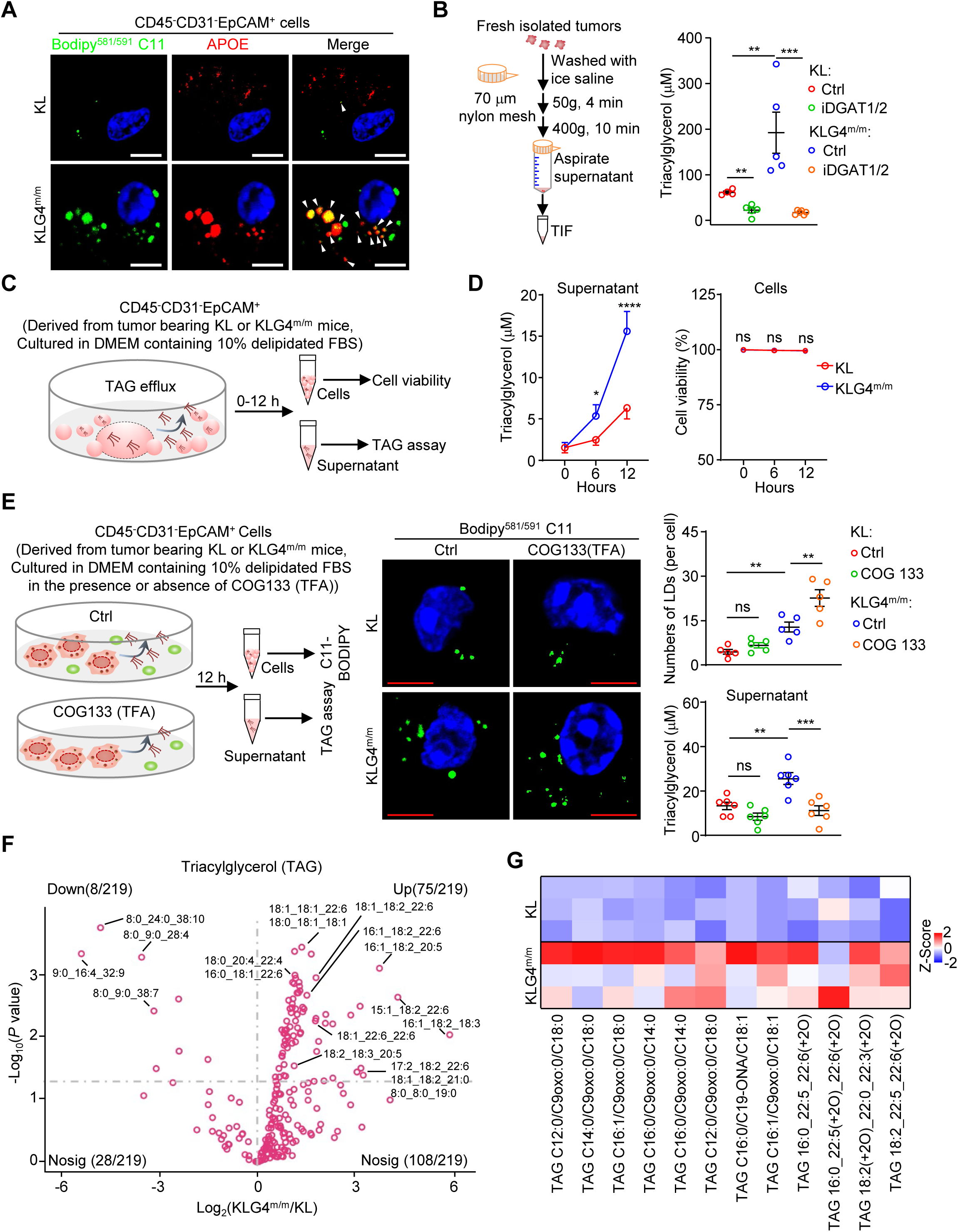
The TAG efflux is elevated in *Gpx4*-null tumor cells of the autochthonous NSCLC models. (A) Representative images of C11-bodipy and anti-APOE staining of CD45^-^CD31^-^EpCAM^+^ tumor cells from lung tumors of KL and KLG4^m/m^ mice. The white arrowheads indicated the simultaneous detection of Bodipy581/591 C11 (green) and APOE (red). (B) A scheme of tumor interstitial fluid (TIF) collection (left) and quantification of the TAG (right) in TIF from lung tumors of KL and KLG4^m/m^ mice that were treated as in Figure 3E. KL, n = 4 and 5 for control and iDGAT1/2, respectively. KLG4^m/m^, n = 5 and 6 for control and iDGAT1/2, respectively. (C, D) A scheme diagram depicting the detection of TAG efflux (C) and the quantification of TAG efflux and cell viability (D) in CD45^-^CD31^-^EpCAM^+^ tumor cells from KL and KLG4^m/m^ mice (n = 4 for each group). (E) A scheme diagram (left) depicting CD45^-^CD31^-^EpCAM^+^ tumor cells from KL and KLG4^m/m^ mice treated with APOE inhibitor (COG133) for 12 h followed by Bodipy^581/591^ C11 staining and measurement of TAG in the culture supernatants. Representative images (middle images) and statistical analysis (top-right graph) of oxidized lipid-containing LDs in KL (n = 5 for Ctrl and COG133) and KLG4^m/m^ (n = 5 for Ctrl and COG133) CD45^-^CD31^-^EpCAM^+^ tumor cells, and quantification of the TAG (bottom-right graph) in culture supernatants from KL and KLG4^m/m^ CD45^-^CD31^-^EpCAM^+^ tumor cells with or without COG133 treatment (n = 6 per group). (F) Volcano plot showing the levels of triacylglycerols (TAG) in the supernatants of CD45^-^CD31^-^EpCAM^+^ tumor cells derived from KLG4^m/m^ mice (n = 5) compared to tumor cells derived from KL mice (n = 5). The tumor cells were cultured in a delipidated medium for 12 h and the supernatants were harvested for lipidomics analysis. (G) Heatmap showing the representative oxidized TAGs (oxTAGs) in the supernatants of CD45^-^EpCAM^+^CD31^-^ tumor cells sorted from lung tumors of KL (n = 3) and KLG4^m/m^ (n = 3) mice. Each column represents z-score normalized intensities of the detected oxTAG species. Graphs show mean ± SEM (B, D, and E). * *P* < 0.05, ** *P* < 0.01, *** *P* < 0.001, **** *P* <0.0001. ns, not significant. Statistical analyses were performed with one-way ANOVA (B and G) and two-way ANOVA (D). Scale bars represent 5 μm (A and E). Data are representative of three independent experiments (A, B, D, and E).

To further substantiate this notion, we isolated the KLG4^m/m^ or KPG4^m/m^ and the KL or KP tumor cells and cultured them in delipidated medium for 12 hours followed by quantification and characterization of TAG in the supernatants (Figure 4C and Figure S8A). The results showed higher levels of TAG were detected in the supernatants of KLG4^m/m^ or KPG4^m/m^ tumor cells than in those of KL or KP tumor cells (Figure 4D and Figure S8A).

Notably, the KLG4^m/m^ tumor cells did not die at 12 h after culture (Figure 4D), which is consistent with the previous observations that deletion of GPX4 in cells results in ferroptosis at a later time (48-72 h) after culture^12^. Treatment with APOE inhibitor COG133 (TFA) significantly downregulated the TAG levels in the supernatants of KLG4^m/m^ tumor cells and increased the lipid droplets in KLG4^m/m^ tumor cells (Figure 4E). In addition, non-targeted and targeted lipidomic assays suggested that TAG and oxTAG levels were significantly increased in the supernatants of KLG4^m/m^ tumor cells compared to KL tumor cells (Figure 4F and 4G and Table S1E and S1F). In contrast, we did not observe a dramatic increase in other ferroptosis-inducing lipid species, such as PC and PE, or free fatty acids (FFA)^50–52^ in the supernatants of KLG4^m/m^ tumor cell cultures compared to the KL counterparts (Figure S8C-S8E and Table S1E and S1F). Collectively, these data suggest that inducible knockout of GPX4 in the autochthonous KL NSCLC model upregulates the expression of APOE to promote the efflux of TAG and oxTAG.

### KLG4^m/m^ tumor cell-secreted lipids promote dysfunction and exhaustion of CD8^+^ T cells

Previous studies have shown that oxidized lipids impair CD8^+^ T cell effector functions in the TME^53,54^. Consistent with the observation that KLG4^m/m^ tumor cells secreted higher levels of TAG and oxTAG than KL tumor cells did (Figure 4C-G), the supernatants from KLG4^m/m^ tumor cell cultures more potently suppressed the production of TNFα, IFNγ, and granzyme B (GZMB) in P14 cells than those from KL tumor cell cultures in an *in vitro* acute activation model (Figure S9A and S9B). Importantly, this suppression was abolished upon delipidation of the supernatants (Figure S9A and S9B). In the chronic stimulation model involving repeated low-dose antigen exposure^55^, culture supernatants from KLG4^m/m^ tumor cells promoted more robust upregulation of Tim3, PD-1, and TOX in P14 cells than those from KL tumor cells (Figure S9C and S9D). This effect was reversed by the delipidation of the supernatants (Figure S9C and S9D), indicating that GPX4-deficient tumor cells secrete lipid mediators that lead to dysfunction and exhaustion of CD8^+^ T cells.

### CD8^+^ T cells in the TME of KLG4^m/m^ tumors exhibit dysfunction and exhaustion

Analysis of the transcriptomic data of CD8^+^ T cells in the TME of KLG4^m/m^ tumors revealed downregulated expression of genes involved in T cell activation pathways and upregulated expression of genes involved in T cell exhaustion compared to those in KL tumors (Figure 5A-C, Figure S10A, and Table S2B and S4), which was confirmed by RT-qPCR assays (Figure S10B). Consistently, KLG4^m/m^ tumors showed significantly lower frequencies and absolute numbers of IFNγ^+^ and granzyme B^+^ (GZMB^+^) CD8^+^ T cells than KL tumors did (Figure 5D). Furthermore, we observed increased proportions of exhausted TOX^+^TCF1^-^ CD8^+^ T cells and upregulated surface expression of the exhaustion markers Tim-3 and PD-1 on CD8^+^ T cells^56^ in KLG4^m/m^ tumors compared to KL tumors (Figure 5D).

**Figure 5.**
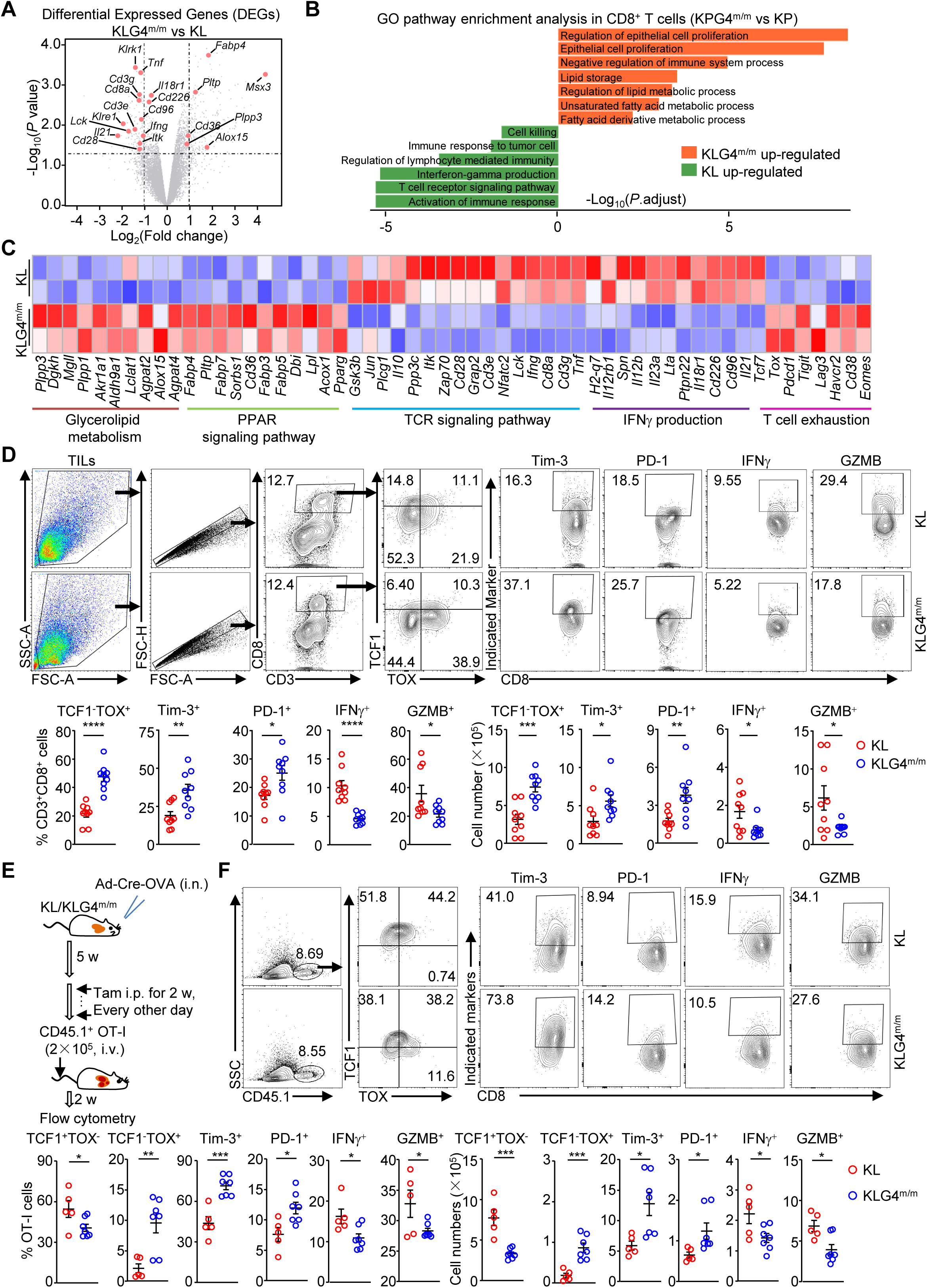
Depletion of *Gpx4* in tumor cells of the autochthonous NSCLC models results in dysfunction of CD8^+^ T cells in the TME. (A) Volcano plot of differentially expressed genes (DEGs, *P* value <0.05 and |log_2_(Fold change)|>=1) in tumor-infiltrated CD8^+^ T cells (CD8^+^ TILs) from KLG4^m/m^ (n = 2) and KL (n = 2) mice that were intranasally injected with Ad-Cre (2 × 10^6^ PFU per mouse) for 5 weeks followed by intraperitoneal injection of tamoxifen (Tam, 80 mg/kg, resolved in corn oil) every other day for 2 weeks and rest for 3 weeks. (B) Gene ontology (GO) analysis of the upregulated and downregulated genes (*P* value < 0.05 and |fold change| >=2) in CD8^+^ TILs from KLG4^m/m^ (n=2) versus KL (n=2) mice treated as in (A). (C) Z-score heatmap of representative genes from transcriptomic data of tumor-infiltrated CD8^+^ T cells from lung tumors of KLG4^m/m^ (n=2) and KL (n=2) mice treated as in (A). (D) Representative flow cytometry images (upper charts) and quantification analysis (lower graphs) of tumor-infiltrated lymphocytes (TILs) from lung tumors of KL (n = 9) and KLG4^m/m^ (n = 9) mice treated as in (A). (E) A scheme of tumor induction plus tamoxifen treatment of KL and KLG4^m/m^ mice followed by adaptive transfer of the naive CD45.1^+^ OT-I cells. KL and KLG4^m/m^ mice were intranasally injected with Ad-Cre (2 × 106 PFU per mouse) for 5 weeks followed by intraperitoneal injection of tamoxifen (Tam, 80 mg/kg, resolved in corn oil) every other day for 2 weeks. Subsequently, CD45.1^+^ OT-I cells (2 × 10^5^ per mouse) were adoptively transferred into the mice for 2 weeks followed by flow cytometry analysis. (F) Representative flow cytometry images (upper charts) and quantification analysis (lower graphs) of the expression of TCF1, TOX, Tim-3, PD-1, IFNγ, and GZMB in OT-I cells from lung tumors of KL (n = 5) and KLG4^m/m^ (n = 7) mice treated as in (E). Graphs show mean ± SEM (graphs of D and E). Statistical analyses were performed using two-tailed student’s *t*-test (D and E). Data are representatives of two independent experiments (D-F). * *P* < 0.05, \*\**P* < 0.01, *** *P* < 0.001, \*\*\*\**P* <0.0001.

Similarly, decreased IFNγ^+^CD8^+^ and GZMB^+^CD8^+^ T cells and increased TOX^+^TCF1^-^ CD8^+^ T cells were found in KPG4^m/m^ tumors compared to KP tumors (Figure S10C). In contrast, the IFNγ^+^CD8^+^ and GZMB^+^CD8^+^ T cells were comparable between the KLG4^m/m^ and the KL bronchial draining lymph nodes (Fig S10D), indicating that inducible knockout of GPX4 in tumor cells results in CD8^+^ T cell dysfunction and exhaustion in the TME of KRas^G12D^ autochthonous NSCLC mice.

To examine whether CD8^+^ T cells were responsible for the tumor control in KL and KLG4^m/m^ mice, we induced tumors in KL and KLG4^m/m^ mice for five weeks followed by two-week injection (i.p.) of tamoxifen and five-week injection (i.p.) of αCD8 or the control IgG (Figure S11A). Results from flow cytometry analysis demonstrated efficient depletion of CD8^+^ T cells in bronchial draining lymph nodes (dLNs) and spleens (Figure S11B). Notably, depletion of CD8^+^ T cells substantially increased the burden and sizes in both KL and KLG4^m/m^ mice (Figure S11C and S11D), indicating that CD8^+^ T cells are required to control tumor progression in the KL NSCLC model.

To examine whether tumor-specific CD8^+^ T cells in the TME were dysfunctional in the TME of KLG4^m/m^ tumors, we intranasally injected KL and KLG4^m/m^ mice with Ad-Cre-P2A-OVA virus for 5 weeks, followed by intraperitoneal injection of tamoxifen every other day for two successive weeks and by adoptive transfer of the CD45.1^+^ naive OTI cells for 2 weeks^33^ (Figure 5E). The results from flow cytometry analyses revealed that the percentages and the absolute numbers of GZMB^+^ and IFNγ^+^ OT-I cells were significantly reduced in the TME of KLG4^m/m^ mice compared to those in KL mice (Figure 5F). Moreover, compared to the counterparts from KL tumors, the OT-I cells from KLG4^m/m^ tumors showed increased TOX^+^TCF1^-^ subpopulations and upregulated expression of Tim3 and PD-1 (Figure 5F).

Collectively, these data demonstrate that inducible knockout of GPX4 in autochthonous KL tumor cells leads to dysfunction and exhaustion of antitumor CD8^+^ T cells to foster an immunosuppressive TME and promote NSCLC progression.

### Inhibition of DGAT1/2 restores CD8^+^ T cell function in the TME of KLG4^m/m^ tumors

We next examined whether treatment with iDGAT1/2 restored CD8^+^ T cell functions in the TME of KLG4^m/m^ mice (Figure 6A). The results from flow cytometry analyses suggested that treatment with iDGAT1/2 significantly increased the intracellular levels of IFNγ and GZMB in and attenuated the expression of PD-1 and Tim-3 on CD8^+^ T cells from KLG4^m/m^ tumors (Figure 6B and 6C). In addition, the percentages and the absolute numbers of exhausted TCF1^-^TOX^+^ CD8^+^ T cells in the TME of KLG4^m/m^ mice were significantly reduced after treatment with iDGAT1/2 (Figure 6B and 6C). Notably, iDGAT1/2 promoted the expression of IFNγ and GZMB in CD8^+^ T cells and reduced the exhaustion of CD8^+^ T cells from KLG4^m/m^ tumors to a level similar to that in those from KL tumors (Figure 6B and 6C), suggesting an essential role of (ox)TAG in the dysfunction and exhaustion of CD8^+^ T cells in the TME. Taken together, these findings indicate that KLG4^m/m^ tumor cells synthesize and secrete TAG (and oxTAG) to promote the dysfunction and exhaustion of CD8^+^ T cells in the TME.

**Figure 6.**
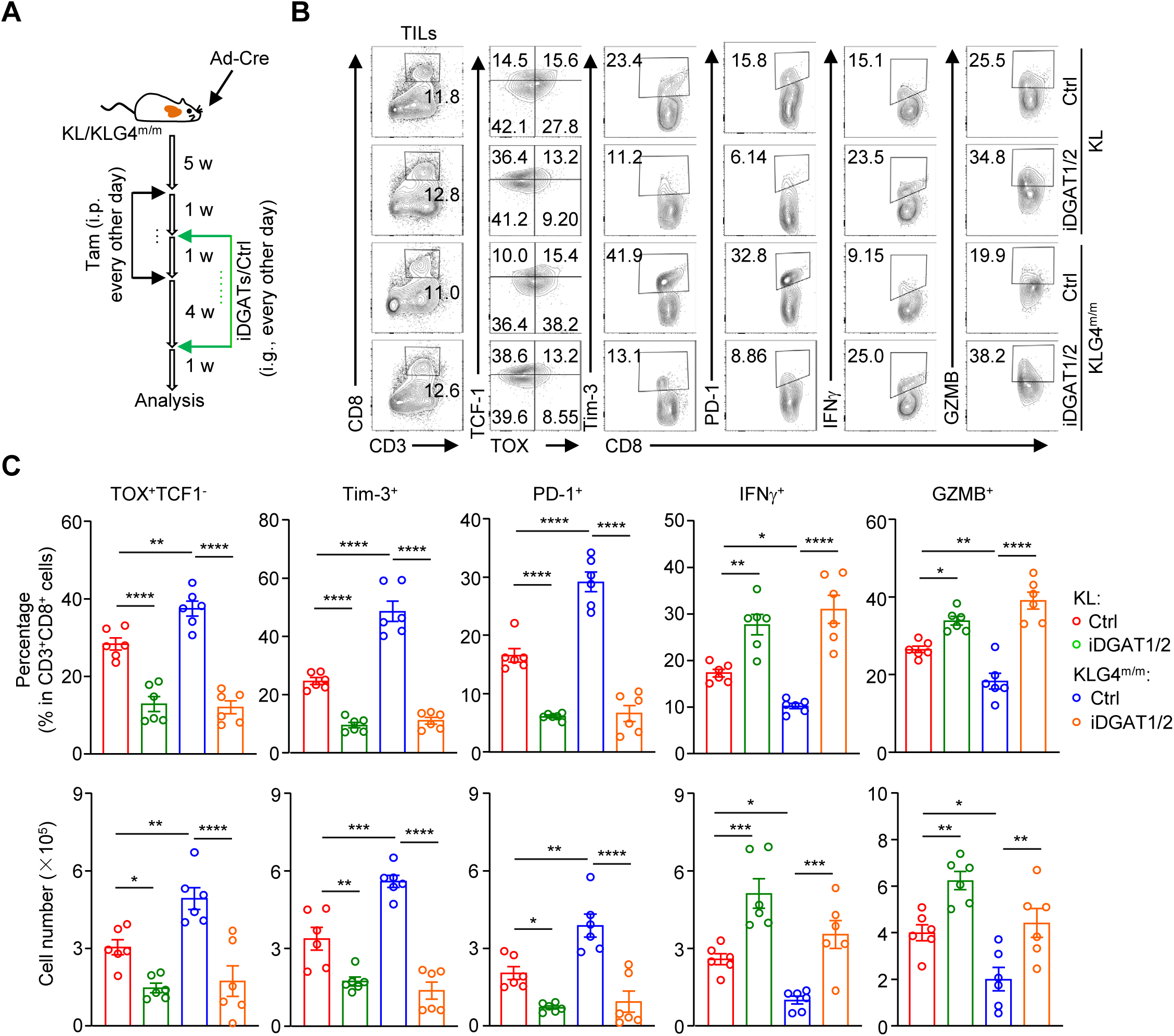
**Inhibition of DGAT1/2 restores CD8^+^ T cell function in the TME of KLG4^m/m^ tumors.** (A) Schematic illustration depicts tumor induction and iDGAT1/2 (composed of T863 and PF06424439) treatment in KL and KLG4^m/m^ mice that were intranasally injected with Ad-Cre (2×10^6^ PFU per mouse) for 5 weeks followed by intraperitoneal injection of tamoxifen every other day for 2 weeks. One week after Tam treatment, the mice were injected with iDGAT1/2 (composed of T863 and PF06424439, 20 mg and 40 mg per kg body weight, respectively) every other day by gavage for 5 weeks. The mice were rested for one week followed by various analyses. (B-C) Representative flow cytometry images (B) and quantification analysis (C) of tumor-infiltrated lymphocytes (TILs) from lung tumors of KL (n = 6 for iDGAT1/2 or control dissolvent) and KLG4^m/m^ (n = 6 for iDGAT1/2 or control dissolvent) mice treated as in (A) under indicated treatment. Graphs show mean ± SEM (C). * *P* < 0.05, ** *P* < 0.01, *** *P* < 0.001, **** *P* <0.0001. ns: not significant (two-way ANOVA for C). Data are representative results of two independent experiments (B, C).

### Inducible expression of GPX4 in tumor cells coordinates an immune-active TME and inhibits NSCLC progression

Having demonstrated that inducible knockout of GPX4 in tumor cells ignites an immunoinhibitory TME to promote NSCLC progression, we next investigated whether inducible expression of GPX4 in tumor cells would enhance antitumor immunity in the TME and inhibit the progression of NSCLC. To test this idea, we generated a line of mice in which GPX4 was inducibly overexpressed in tumor cells (termed G4^OE^ mice hereafter) based on Cre-loxP and tetracycline-controlled (tet-on) gene expression systems (Figure S12A). PCR analysis of the tail genomic DNA indicated that the targeting vector was successfully knocked into the *H11* site (Figure S12B). GPX4 was inducibly overexpressed only after Cre recombinase-mediated removal of the STOP cassette and in the presence of doxycycline (Dox) (Figure S12C and S12D). Next, we obtained the *Kras*^LSL-G12D/+^*Lkb1*^fl/fl^*Gpx4*^OE^ (KLG4^OE^) mice and induced tumorigenesis in these mice by intranasal injection of Ad-Cre followed by feeding with Dox-supplemented food (Figure 7A), and found that GPX4 was selectively upregulated in CD45^-^EpCAM^+^CD31^-^ tumor cells but not in endothelial cells (CD45^-^CD31^+^CD49d^-/int^), stromal cells (CD45^-^CD31^int^CD49d^+^) or CD45^-^CD31^-^EpCAM^-^ cells in tumors, or other organs (including the liver, spleen, kidney, and brain) from KLG4^OE^ mice fed Dox-supplemented food compared to those fed normal food (Figure S12E and S12F), suggesting efficient, specific and inducible overexpression of GPX4 in tumor cells in the KLG4^OE^ autochthonous NSCLC mouse model.

**Figure 7.**
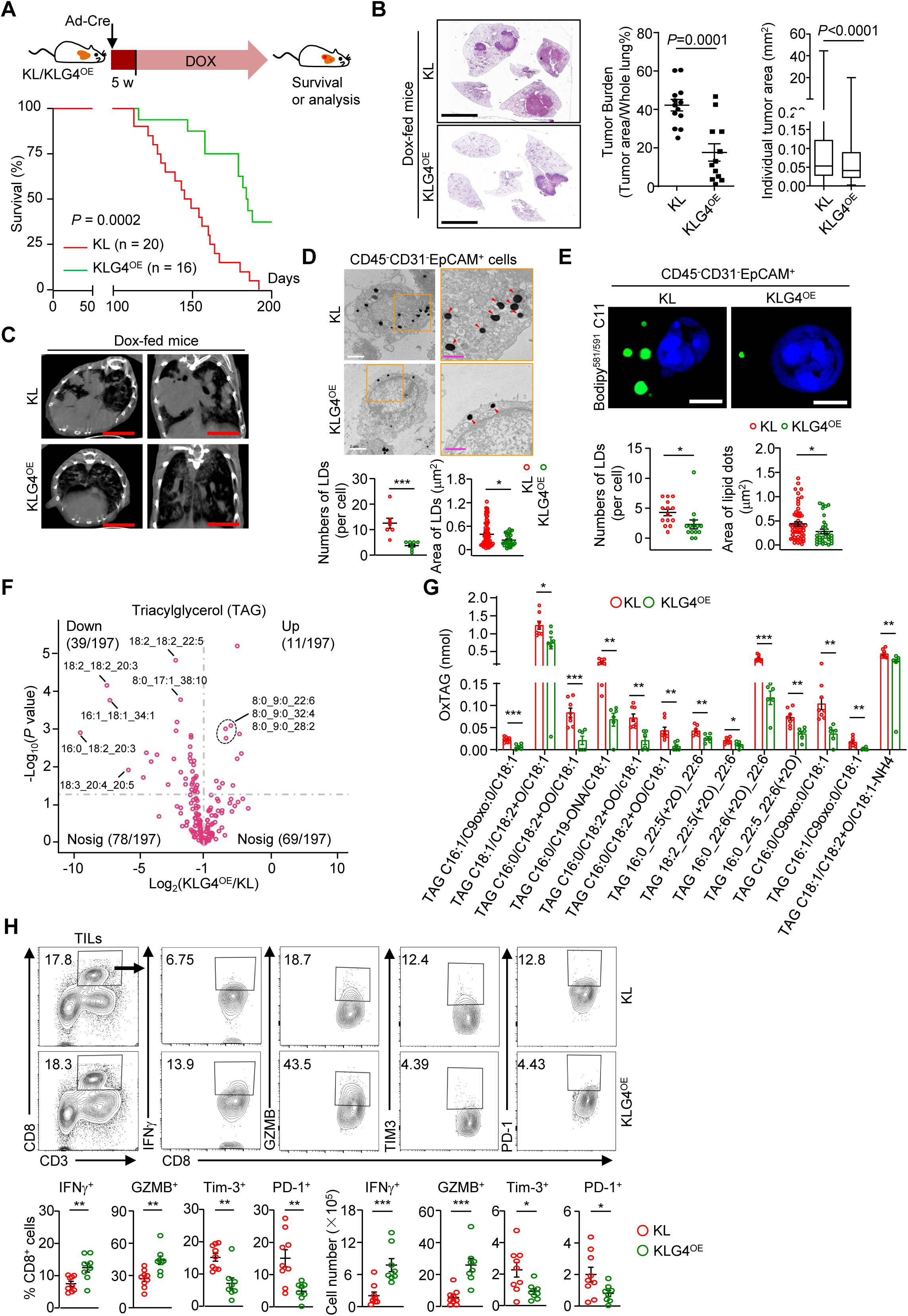
Inducible overexpression of *Gpx4* in tumor cells alleviates NSCLC progression. (A) A scheme of tumor induction (upper) and the survival (lower) of *Kras*^LSL-G12D/+^*Lkb1*^fl/fl^ (KL, n = 20) and *Kras*^LSL-G12D/+^*Lkb1*^fl/fl^*Gpx4*^OE^ (KLG4^OE^, n = 16) mice that were intranasally injected with Ad-Cre (2×10^6^ PFU/mice) for 5 weeks followed by feeding with Dox- supplemented chow diet until the endpoint of study. (B, C) Representative images of HE staining (b, left) and statistics of tumor burdens (b, middle) and individual tumor sizes (b, right) and micro-CT (c) of tumor-burdened lungs from the KL (n = 7) and KLG4^OE^ (n = 9) mice treated as in (A). (D) Representative images from transmission electron microscopy (upper) and quantification of lipid droplets (LDs) (lower graphs) of CD45^-^CD31^-^EpCAM^+^ tumor cells from lung tumors of KL (n = 7) and KLG4^OE^ (n = 8) mice treated as in (A). The orange boxed areas were shown at higher magnification in the bottom images in which arrowheads indicated LDs. (E) Representative images of confocal microscopy (upper) and quantification of the numbers and sizes (lower graphs) of oxidized lipids-containing LDs in CD45^-^CD31^-^EpCAM^+^ tumor cells from KL and KLG4^OE^ mice (n = 14 per group) treated as in (A). (F) Volcano plot displaying the triacylglycerols (TAGs) in the culture supernatant of CD45^-^CD31^-^EpCAM^+^ tumor cells from KLG4^OE^ mice (n = 6) compared to those from KL mice (n = 8). (G) Quantitative assessment showing the oxidized triacylglycerols (oxTAG) in the supernatant of CD45^-^CD31^-^EpCAM^+^ tumor cells culture from lung tumors of KL (n = 8) and KLG4^OE/+^ (n = 6) mice treated as in (A). (H) Representative images and quantitative analysis of flow cytometry assays of TILs from lung tumors of KL (n = 9) and KLG4^OE^ (n = 8) mice treated in (A). Graphs show mean ± SEM (B, D, E, G, and H). Statistical analysis was performed using Log-rank analysis (A), two-tailed student’s *t*-test (B, D, E, G, and H) or multiple *t*-tests (G). Scale bars represent 5 mm (B and C), 2 μm (white, D), 1 μm (magenta, D), or 5 μm (E). Data are combined results of three independent experiments (A) or representative results of two independent experiments (B-E, H). * *P* < 0.05, ** *P*<0.01, *** *P* < 0.001, **** *P* <0.0001.

We next assessed the effects of inducible GPX4 overexpression in tumor cells on NSCLC progression with the KLG4^OE^ mice. As shown in Figure 7A, the KLG4^OE^ mice survived significantly longer than the KL mice did. The results of histological analysis and micro-CT imaging suggested that the total tumor burden and the individual tumor size were significantly reduced in the lungs of KLG4^OE^ mice compared to those in the lungs of KL mice (Figure 7B and 7C), suggesting that inducible overexpression of GPX4 in tumor cells inhibits the progression of NSCLC in the autochthonous KL mouse model. Interestingly, genes involved in TAG synthesis and efflux such as *Dgat1/2*, *Gpd1l*, *Gpam* and *Apoe* were significantly downregulated in the KLG4^OE^ tumor cells compared to the KL tumor cells (Figure S12G), and the lipid droplets in KLG4^OE^ tumor cells were substantially less and smaller than in KL tumor cells (Figure 7D), which aligns with the observation that higher GPX4 mRNA levels correlated with lower *DGAT2*, *GPD1L* and *APOE* in human NSCLC tissue (Figure S5C). Notably, the lipid droplets contained fewer oxidized lipids in KLG4^OE^ tumor cells than in KL tumor cells as indicated by the C11-BODIPY staining (Figure 7E).

Consistently, fewer TAG and oxTAG were produced by the KLG4^OE^ tumor cells than the KL tumor cells (Figure 7F, 7G and Table S1G and Table S1H). These data suggest that inducible overexpression of GPX4 in tumor cells reduces the synthesis and efflux of TAG and oxTAG.

We further found that the expression of intracellular IFNγ and GZMB in CD8^+^ T cells was potentiated, whereas the surface exhaustion markers Tim-3 and PD-1 were reduced in CD8^+^ T cells from the TME of KLG4^OE^ tumors compared to those from the control counterparts (Figure 7H), which aligns with the TCGA and scRNA-seq data showing that high expression of *GPX4* is correlated with increased survival of pancreatic ductal adenocarcinoma patients^16^ and decreased T cell exhaustion score in human NSCLC tissue^45,57–59^ (Figure S13). Collectively, these data suggest that inducible overexpression of GPX4 in tumor cells promotes the antitumor functions of CD8^+^ T cells to inhibit NSCLC progression.

## Discussion

It has been demonstrated that knockout of GPX4 in cancer cell lines results in ferroptosis and that inducible deletion of GPX4 in mice causes renal failure and death within two weeks^12^. However, our results showed that inducible deletion of GPX4 in tumor cells in the KL and KP autochthonous NSCLC mouse models had little effect on the ferroptosis of tumor cells, indicating that the tumor cells in these models adapt to GPX4 deficiency-induced ferroptotic stress. In support of this notion, we found that the peroxidation of PE or PC whose products are classical ferroptosis inducers^12,37–39^ was comparable between the KLG4^m/m^ and the KL tumor cells. In contrast, genes involved in TAG synthesis, such as *Gpd1l*, *Gpam*, *Agpat4*, *Plpp1*, *Dgat2* and *Srebf2*, were upregulated, and the levels of TAG and oxTAG and the lipid droplets were increased in KLG4^m/m^ tumor cells compared to KL tumor cells. In addition, inhibition of DGAT1/2 significantly downregulated the levels the PUFA-TAG and oxTAG and upregulated the levels of oxPE and oxPC in KL and KLG4^m/m^ tumor cells, indicating that the GPX4-DGAT1/2 axis reprograms ferroptotic oxPE/PC metabolism to oxTAG synthesis and thereby inhibits ferroptosis of KL and KLG4^m/m^ tumor cells. Together with the observations that TAG biosynthesis confers resistance to ferroptosis^42,43,60^, these findings suggest that tumor-cell-specific GPX4 deficiency reprograms TAG metabolism to evade ferroptosis in autochthonous NSCLC mouse models (Figure S14). When this study was in the review and revision process, a publication showed that sgRNA-mediated knockout of *Gpx4* inhibited NSCLC progression in the *Kras*^LSL-G12D/+^*Tp53*^fl/fl^*Rosa26*^LSL-Cas9/LSL-Cas9^ mouse model^61^. However, it should be noted that *Gpx4* was deleted at the time of KRas^G12D^ expression. While acute deletion of GPX4 results in ferroptosis within 48-72 h^12^, KRas^G12D^-induced malignant transformation requires 2-3 weeks^26,62^. Therefore, such an early deletion of *Gpx4* would result in death of normal and incompletely transformed epithelial cells, which expectedly inhibit NSCLC progression and might be the possible reason for the descrepancies between that study and ours.

In the syngeneic graft mouse models, however, knockout or inducible knockout of GPX4 in tumor cells resulted in compromised growth and potentiated ferroptosis that was restored by Lip-1, which is consistent with previous reports^15,38,39^. It should be noted that the TME between the autochthonous models and the syngeneic models is different, which might be responsible for the opposite phenotypes of the tumor cell-specific GPX4-knockout syngeneic and autochthonous NSCLC models. For the syngeneic model, a large number of tumor cells are inoculated subcutaneously, and few immune cells are identified within the tumors. In contrast, in the autochthonous NSCLC models and human NSCLC tissues, the tumor cells originate from a few mutated malignant cells, undergo constant immune editing by the immune cells in the TME, and account for a small portion of the whole tumor tissue^31,63^. This autochthonous versus syngeneic differences are not unique to GPX4, as it has been observed that knockout of FSP1 did not affect tumor cell growth in in vitro cultured tumor cells or in the syngeneic or xenograft models^61^. Although the exact TME factors responsible for the distinct behaviors between GPX4-deficient autochthonous and syngeneic lung tumors are unclear, our study highlights the caution in interpreting the syngeneic models that are extensively used to evaluate the pro- or anti-tumor activity of a specific gene or compound.

We observed that knockout of GPX4 in autochthonous KL tumor cells, but not in subcutaneous syngeneic KL tumor cells or normal lung epithelial cells, led to the upregulation of *Dgat2* and *Gpd1l* mRNA levels, and such expression difference of *Dgat2* and *Gpd1l* mRNA between the autochthonous KL and KLG4^m/m^ tumor cells was gradually diminished in *in vitro* cultures, indicating that factors in the TME license the upregulation of genes involved in TAG metabolism. It should be noted that the lung tumor identity may be altered during *in vitro* culture or subcutaneous transplantation, characterized by the loss of lineage-specific markers such as TTF-1 (also known as NKX2-1) after the removal from the pulmonary microenvironment^64,65^. Because TTF-1 interacts with transcriptional factors Foxa1 and Foxa2 that bind to lung-specific loci to maintain pulmonary identity, the loss of TTF-1 might result in an altered binding of Foxa1/2 to genes involved in TAG synthesis^65,66^. In this context, it has been demonstrated that Foxa1 is a potent inhibitor of hepatic TAG synthesis by repressing the expression of genes such as *GPAM* and *DGAT2*^67^. In addition, we observed that H3K4me3 and H3K27ac modifications on the loci of *Dgat2* and *Gpd1l* were significantly increased in autochthonous KLG4^m/m^ tumor cells but not in syngeneic KLG4^m/m^ tumor cells compared to the respective KL counterparts, indicating that the epigenetic modification factors are involved in such differential regulatory processes. In support of this notion, we found that the autochthonous KLG4^m/m^ tumor cells produced higher amounts of TAG and oxTAG and exported them into the extracellular space, leading to CD8^+^ T-cell functional impairment, which was substantially inhibited by iDGAT1/2. A thorough understanding of how GPX4 coordinates the identity-maintaining machinery and epigenetic factors in autochthonous versus syngeneic to reprogram TAG metabolism requires further investigation. Nonetheless, these data indicate that TAG and oxTAG from tumor cells shape an immunosuppressive TME to promote NSCLC progression.

Our data demonstrate that inducible overexpression of GPX4 in tumor cells ignites an immune-active TME characterized by potentiated activation of CD8^+^ T cells, leading to compromised NSCLC progression of the autochthonous models. Together with the observations that lipid peroxidation suppresses CD8^+^ T cell effector functions^53,54,68^, boosting the expression or the function of GPX4 in the TME would be a benefit for the NSLC patients. In this context, it has been reported that high expression of GPX4 is correlated with increased survival in pancreatic ductal adenocarcinoma (PDAC) patients^16^, and the levels of *GPX4* mRNA were negatively correlated with the levels of *DGAT2*, *GPD1L*, and *APOE* mRNA in human NSCLC tumor cells and the T cell exhaustion score in human NSCLC tumor tissues. In addition, we provided evidence that inhibition of TAG synthesis via iDGAT1/2^43,47^ significantly decreased the synthesis and the efflux of TAG (oxTAG) and the accumulation of lipid droplets in tumor cells. Accordingly, iDGAT1/2 treatment sensitizes tumor cells to ferroptosis and attenuates tumor progression in the autochthonous NSCLC models. Notably, two phase 2 clinical trials have shown high safety and efficacy of the DGAT2 inhibitor (ervogastat, PF-06865571) in non-alcoholic fatty liver disease metabolism^69^. It is worth noting the future clinical use of ervogastat and PF-06865571in NSCLC. Taken together, our findings reveal the preclinical outcomes of specific ablation of GPX4 in tumor cells and highlight potential therapeutic interventions for NSCLC patients.

## Methods

### Mice

The *Kras*^LSL-G12D/+^ (#008179), the *Tp53*^fl/fl^ (#008462), and the *Lkb1*^fl/fl^ (#014143) mice were purchased from the Jackson Laboratory and crossed to obtain the *Kras*^LSL-G12D/+^*Tp53*^fl/fl^ (KP) and the *Kras*^LSL-G12D/+^*Lkb1*^fl/fl^ (KL) strains as previously described^29–31^. The *Kras*^LSL-G12D/+^*Tp53*^fl/fl^*Gpx4*^m/m^ (KPG4^m/m^) and the *Kras*^LSL-G12D/+^*Lkb1*^fl/fl^*Gpx4*^m/m^ (KLG4^m/m^) mice were generated by GemPharmatech following these steps. The eggs from wild-type C57BL/6 mice were fertilized *in vitro* with sperm from KP or KL mice, both on the C57BL/6 background.

The targeting vector containing the rox-*Gpx4* (exon 5 to exon 7)-rox-DreER^T2^-loxP2272-STOP-loxP2272-CAG promoter and the flanking homologous sequences of the *Gpx4* gene locus, along with the gRNA-Cas9 complex, was subsequently injected into the zygotes to obtain the *Kras*^LSL-G12D/+^*Tp53*^fl/+^*Gpx4*^m/+^or the *Kras*^LSL-G12D/+^*Lkb1*^fl/+^*Gpx4*^m/+^ mice. Since the *Gpx4* gene locus is approximately 60 kb upstream of the *Lkb1* gene locus, which contains two loxp66 sites (5’-ATAACTTCGTATAGCATACATTATACGAAGTTAT-3’), two distinct Cre recombinase recognition sites, loxP2272 sites (5’-ATAACTTCGTATAAAGTATCCTATACGAAGTTAT-3’) flanking the STOP cassette within the *Gpx4* locus, were introduced into the targeting vector to ensure the correct removal of the STOP cassette and the *Lkb1* exon 3 to 6 in the presence of Cre recombinase. The F0 *Kras*^LSL-G12D/+^*Tp53*^fl/+^*Gpx4*^m/+^ or *Kras*^LSL-G12D/+^*Lkb1*^fl/+^*Gpx4*^m/+^ mice were then crossed with KP or KL mice for at least five generations to obtain the *Kras*^LSL-G12D/+^*Tp53*^fl/fl^*Gpx4*^m/+^ (KPG4^m/+^) or the *Kras*^LSL-G12D/+^*Lkb1*^fl/fl^*Gpx4*^m/+^ (KLG4^m/+^) mice, respectively. The KPG4^m/+^ or KLG4^m/+^ mice were crossed to obtain the *Kras*^LSL-G12D/+^*Tp53*^fl/fl^*Gpx4*^m/m^ (KPG4^m/m^) or *Kras*^LSL-G12D/+^*Lkb1*^fl/fl^*Gpx4*^m/m^ (KLG4^m/m^) mice for maintenance and experiments, respectively. In addition, the F0 *Tp53*^fl/+^*Gpx4*^m/+^ mice were crossed with wild-type C57BL/6 mice for at least three generations to obtain the *Gpx4*^m/+^ mice that were crossed to generate the *Gpx4*^m/m^ mice for maintenance and experiments.

The *Gpx4*^OE^ mice were generated by GemPharmatech through CRISPR/Cas9-mediated gene editing. In brief, the targeting vector consisting of pTRE3G promoter-loxp-STOP-loxp-m*Gpx4* (NM_001367995.1)-polyA-rTTA-CAG promoter and the flanking homologous sequences of the *H11* site and the gRNA-Cas9 RNPs were injected into the in vitro obtained zygotes of wild-type C57BL/6 mice. The F0 *Gpx4*^OE^ mice were crossed with the KL mice to obtain the *Kras*^LSL-G12D/+^*Lkb1*^fl/+^*Gpx4*^OE^ that were crossed with the KL mice for at least six generations to obtain the *Kras*^LSL-G12D/+^*Lkb1*^fl/fl^*Gpx4*^OE^ (KLG4^OE^) mice for maintenance and experiments. Alternatively, the F0 *Gpx4*^OE^ mice were crossed with wild-type C57BL/6 mice for three generations to obtain the *Gpx4*^OE^ mice that were used for experiments.

The CD45.1^+^ OT-I mice carrying a transgenic TCR specific for H-2K^b^ and OVA_257–264_ were purchased from Cyagen Biosciences (Suzhou, China). The wild-type C57BL/6 mice were purchased from GemPharmatech Co., Ltd (Nanjing, China). The sequences of gRNAs were 5’-GGCTGTCTTCCGGCCTTGGA-3’ and 5’-TGCATGCTTGAAGCCCTCCA-3’ for *Gpx4*^m/+^ mice, and 5’-CTGAGCCAACAGTGGTAGTA-3’ for *Gpx4*^OE^ mice, respectively.

No statistical methods were used to predetermine the sample size. For all experiments presented in this study, age- and sex-matched mice were used and the sample sizes were large enough to determine statistically significant effects. The control and experimental groups of mice were cohoused and randomly allocated to different treatments. All mice were housed in the specific pathogen-free animal facility at Medical Research Institute, Wuhan University with a housing temperature of 22±1℃ and relative humidity of 50-60 % with a 12-hour dark/12-hour light cycle and fed with standard food and water otherwise indicated. All animal experiments were performed in accordance with protocols approved by the Institutional Animal Care and Use Committee of Wuhan University (Approval No.21020A).

### Genotyping

Genomic DNA was prepared from the tails of 4-week-old mice. The tissues were lysed in the lysis buffer (0.5 M Tris-HCl, pH 8.5, 5 mM EDTA, 0.2 % SDS, and 0.8 μg/μL proteinase K) for overnight at 65℃. After incubation, the samples were centrifuged at 10,000 g for 10 min at room temperature to obtain a supernatant containing the genomic DNA. The supernatant was transferred to a new 1.5 mL Eppendorf tube containing 1 mL 100% ethanol and mixed thoroughly. Subsequently, the mixtures were centrifugated at 10,000 g for 10 min at room temperature. The supernatants were discarded, and the resulting precipitants containing DNA were dissolved in deionized water for subsequent PCR analysis. The PCR cycling conditions were as follows: 94°C for 2 min; 94°C for 30 seconds; 58°C (61°C for *Kras*) for 30 seconds; 72°C for 30 seconds; repeat steps 2 through 4 for a total of cycles; 72°C for 5 min; 16°C for 10 min, unless indicated otherwise. The PCR products were analyzed by agarose gel electrophoresis as previously described ^29–31^. The primers used for genotyping are summarized in Table S5.

### Cell culture and *in vitro* treatment

Primary mouse lung fibroblasts (MLFs) were isolated from ∼8-10-week-old mice. Lungs were minced and digested in calcium and magnesium-free HBSS buffer supplemented with 10 mg/mL type I collagenase (Worthington) and 20 μg/mL DNase I (Sigma-Aldrich) for 2.5 h at a cell culture incubator with intervals of pipette. The cell suspensions were centrifuged and filtered through a 70 μm nylon mesh followed by culture in DMEM containing 15% FBS (v/v), 1% streptomycin–penicillin. Two days later, the adherent fibroblasts were rinsed with pre-warmed PBS and cultured in a 6-well plate for subsequent experiments.

For MLFs derived from *Gpx4*^+/+^ and *Gpx4*^m/m^ mice, cells were infected with Ad-Cre for 48 h followed by treatment with 4-hydroxytamoxifen (4OHT, 1μM, Sigma, Cat# H6278) for 48 h. The cells were harvested for immunoblot analysis. For MLFs derived from *Gpx4*^+/+^ and *Gpx4*^OE^ mice, cells were infected with Ad-Cre, one day later, Doxycycline (20 μg/mL) was added to the cell medium for another 3 days. The cells were harvested for RT-qPCR and immunoblot analysis.

### Preparation and infection of Ad-Cre-P2A-OVA viruses

The pDC316-mCMV-CRE-P2A-OVA vector (DesignGene, Shanghai) was co-transfected with the package plasmid pBHGlox(delta)E1,3Cre into HEK293A cells for 10 h^33^. After transfection, the medium was replaced with fresh DMEM (supplemrnted with 10% FBS and 1% streptomycin-penicillin) and cells were cultured for an additional five days. Following incubation, cells were harvested via centrifugation at 3,000 g for 5 min, resuspended in DMEM, and subjected to three freeze-thaw cycles at -80℃ and 37℃. The viruses in the supernatants were collected by centrifugation at 15,000 g for 10 min, followed by three rounds of infection and collection from HEK293A cells. The resulting supernatants, containing Ad-Cre-OVA viruses, was filtered, collected, and stored at -80℃. To determine viral titers, aliquots or serial dilutions of the Ad-Cre-OVA viruses were used to infect 3T3^LSL-RFP^ cells (kindly provided by Dr. Hong-Bin Ji, Chinese Academy of Science, Shanghai) for 48 h, followed by flow cytometry analysis to determine the titers. For tumor induction, age- and sex-matched KL/KLG4^m/m^ mice were intranasally administered with Ad-Cre-P2A-OVA (5× 10^6^ PFU per mouse), and 5 weeks later, the mice were intraperitoneally injected with tamoxifen every other day for 2 weeks. The mice were either rested for 3 weeks followed by histological analyses or rested for 2 weeks followed by adoptive transfer of CD45.1^+^ OT-I T cells (2×10^5^ per mouse) and analyzed after an additional 2 weeks.

### Induction of the autochthonous NSCLC tumors

The experiments were performed as previously described^29–31^. 8-10-week-old, sex-matched KL/KLG^m/m^, KP/KPG4^m/m^ or KL/KLG4^OE^ mice were anesthetized by intraperitoneal injection of 0.7% sodium pentobarbital (w/v, 10 μL/g body weight), followed by intranasal injection of Ad-Cre viruses (HANBIO, Shanghai) (2×10^6^ PFU in 60 μl PBS per mouse) or Ad-SPC-Cre (5×10^6^ PFU in 60 μL PBS per mouse) (kindly provided by Dr. Hongbin Ji, Chinese Academy of Science, Shanghai)^34^ for 5 weeks. Subsequently, the KL/KLG4^m/m^ and KP/KPG4^m/m^ mice were intraperitoneal injected with Tamoxifen (Tam, 10 mg/mL dissolved in corn oil and 80 mg/kg body weight) every other day for 2 weeks and rested until death for survival analysis, or rested for another 3 weeks for subsequent analysis. The KL/KLG4^OE^ mice were fed with normal food until the fifth week after Ad-Cre injection. The mice were fed with Dox-supplemented food (+Dox) until death for survival analysis or for 8 weeks for subsequent analysis.

### Treatment with iDGAT1/2 in autochthonous KL NSCLC mouse models

The KL/KLG4^m/m^ mice were intranasally injected with Ad-Cre and administered tamoxifen intraperitoneally for two weeks as described above. One week after initiating the tamoxifen treatment, the mice were randomly assigned to different treatment groups, and followed by iDGAT1/2 treatments. The iDGAT1/2, consisting of T863 (a DGAT1 inhibitor^43,47^, 20mg per kg of body weight, HY-32219, MedChemExpress) and PF-06424439 (a DGAT2 inhibitor^47^, 40mg per kg of body weight, HY-108341, MedChemExpress), were solubilized in a solvent containing 5% DMSO, 30%PEG300, 5% Tween-80 and 60% H_2_O. The iDGAT1/2 and the control were administered by gavage every other day and continued for 5 weeks. One week later, the mice were anesthetized for micro-CT scanning, and the lungs from tumor-bearing mice were collected for subsequent histological analysis or flow cytometry.

### CD8^+^ T cell depletion in tumor-bearing KL/KLG4^m/m^ mice

The KL and KLG4^m/m^ mice were intranasally injected with Ad-Cre for 5 weeks followed by intraperitoneal administration of tamoxifen for 2 weeks. After completion of tamoxifen treatment, the mice were randomly assigned to different treatment groups for intraperitoneal injection with αCD8α (200 μg per injection, clone: 2.43, Selleck, cat#A2102) or IgG2b (200 μg per injection, clone: LTF-2, Selleck, cat#A2116) twice a week for 5 consecutive weeks.

One week later, the mice were anesthetized for micro-CT scanning, the spleens and the bronchial draining lymph nodes were collected for flow cytometry analysis, and the lungs from tumor-bearing mice were collected for subsequent histological analysis.

### Hematoxylin and Eosin (HE) staining

HE staining was performed as previously^29–31^. Briefly, lung tissues were fixed with 2.5 mL 4% paraformaldehyde (PFA) for 4 hours followed by dehydratation in 75%, 95%, 100% ethanol successively (1 hour for each gradient). The lungs were embedded in paraffin and sectioned (5 μm) for subsequent staining with hematoxylin and eosin (Beyotime Biotech). Images were acquired using an Aperio VERSA 8 (Leica) multifunctional scanner. Tumor burden and individual tumor size were determined through ImageScope (Leica) as described previously^29–31^.

### Multi-color immunohistochemistry (mIHC)

The experiments were performed with an Opal 5-color Manual IHC Kit (Absin, Cat #: abs50013) following the manufacturer’s instructions. The slides were deparaffinized in xylene and successively rehydrated in 100%, 95%, and 75% ethanol. The antigen retrieval was performed by heating slides in a microwave for 30 minutes in sodium citrate buffer (pH 6.0). The sections were cooled down naturally to room temperature and quenched in 3% hydrogen peroxide to block endogenous peroxidase activity followed by blocking in 10% horse serum in 1×PBS at room temperature for 15 min. The sections were treated with Click-iT^TM^ TUNEL Colorimetric IHC Detection Kit (Invitrogen, Cat# C10625) according to the manufacturer’s instructions and stained with the following antibodies: rabbit anti-mouse EpCAM (Abclonal, Cat #: A19301, 1:400) at room temperature for 1 h, and rabbit derived 4-HNE (Abcam, Cat #: ab46545, 1:500) at 4℃ overnight. A secondary horseradish peroxidase-conjugated antibody (Absin, cross-react with mouse/rabbit) was added and incubated at room temperature for 15 min. Signal amplification was performed using TSA working solution diluted at 1:100 in 1×amplification diluent (Absin) and incubated at room temperature for 15 min. After each cycle of staining, heat-induced epitope retrieval was performed to remove all the antibodies including primary antibodies and secondary antibodies. The samples were counterstained for nuclei with DAPI for 10 min after all the antigens above have been labelled and mounted in mounting medium. The multispectral images were scanned by ZEISS AXIOSCAN7 at 20× magnification and were analyzed with SliderViewer (Version 2.5).

### Micro-CT scanning

Tumor-burdened mice were anesthetized under 1.0%–1.5% isoflurane via a respiratory mask of the inhalation anesthesia machine and scanned by a NEMO^@^ Micro-CT system (NMC200, PINGSENG TECHNOLOGY, China) to assess the lung tumor burdens. A pneumatic pillow was positioned on the thoraxes of tumor-burdened mice and connected to a pressure transducer to monitor respiratory motion and inform prospective gating. Reconstruction was performed using Avatat3 software with a ‘CT-gating’ strategy and an ‘interaction’ algorithm. Other parameters including the resolution of CT is 1k×1k, the number of iterations is 6, and the gated phase is 8, unless indicated otherwise, reminding parameters were set with default parameters according to the manufacturer’s instructions. The micro-CT images were analyzed by Avatar3 and saved in tiff format.

### Quantitative Real-Time PCR

These experiments were performed as previously described^31,70^. Total RNA was extracted from cells using TRIzol reagent (Takara), and the first-strand cDNA was reverse-transcribed with All-in-One cDNA Synthesis SuperMix (Aidlab, Cat: 342123AH). Gene expression was examined with a Bio-Rad CFX Connect system by a fast two-step amplification program with 2×SYBR Green Fast qPCR Master Mix (Aidlab, Cat: 342123AX). The expression levels of genes were normalized to that of the gene encoding β-actin. Gene-specific primers used in this study are summarized in Table S5.

### Isolation of tumor interstitial fluid (TIF)

We used a previously reported method to isolate tumor tissue interstitial fluid via a centrifugation method^71,72^. Briefly, tumor-bearing lungs were harvested and washed in pre-cold saline for three times, and drying the tumors via blotting paper after the saline rinse. Then the tumors were placed on 70 μm nylon mesh in a 50 mL Falcon tube and spun at < 50g for 5 min to remove surface liquid. Next, samples were centrifuged at 400g for an additional 10 min at 4 ℃ and carefully aspirate the supernatant for interstitial fluid collection.

### Triacylglycerol (TAG) measurement

Triacylglycerol levels in supernatants from cultured cells or tumor interstitial fluid (TIF) from lung tumors were quantified using the Triglyceride-Glo^TM^ assay kit (Promega, Cat #J3160) according to the manufacturer’s instructions. Briefly, primary CD45^-^CD31^-^EpCAM^+^ tumor cells of KL and KLG4^m/m^ were sorted and washed with cold PBS. These cells were then cultured in a 24-well plate at a density of 5×10^6^ cells per well, using 300 μL DMEM supplemented 10% delipidated FBS (v/v, kindly provided by Dr. Yan Wang, Wuhan University). At the indicated time points (0, 6, and 12 h) after initiating the culture, the supernatants were collected. A 25 μL aliquot of the supernatant was transferred to a 96-well plate containing 25 μL Glycerol Lysis Solution with lipase. This mixture was gently shaken and incubated for 30 min at 37 ℃. Following this, 50 μL glycerol detection mix was added to each well and incubated at room temperature for 1 h. Luminescence was measured using a plate-reading luminometer, and glycerol concentrations were calculated. For TAG measurement in TIF, the TIF was collected as described above and subsequently diluted and quantified according to the manufacturer’s instructions..

### Immunoblot assays

The immunoblot assays were performed as described previously^31,70^. In brief, cells were lyzed and the normal tissue or tumor tissues were homogenized with NP-40 lysis buffer (150 mM NaCl, 1 mM EDTA, 1% nonidet P-40) supplemented with proteinase and phosphatase inhibitors (Biotool). The cell lysates or tissue homogenates were cleared by centrifuge at 15000 g for 10 min at 4°C. The supernatants were quantified and loaded to 12% sodium dodecyl sulfate-polyacrylamide (SDS-PAGE) gel for electrophoresis followed by transfer onto nitrocellulose membranes. Blocking was performed in 5% skim milk (w/v) in PBS for 40 min at room temperature, and the membranes were incubated with primary antibodies for overnight at 4℃ and followed by TBST wash every 10 minutes for three times. Subsequently, the membranes were incubated with horseradish peroxidase (HRP)-conjugated secondary antibodies for 1 h followed by incubation with the enhanced chemiluminescence kit (Bio-rad, Cat# 1705061) for analysis. The primary and secondary antibodies used in this study were listed in Table S6.

### Preparation of single-cell suspensions from lung tumors

The freshly tumor-burdened lungs from KL/KLG4^m/m^ mice with or without indicated treatments, as well as KL/KLG4^OE^ mice were perfused through alveolar lavage and cardiac lavage with cold PBS. The tumor-burdened lungs or lung tumors were isolated and cut into small pieces (1∼2 mm in diameter) followed by digesting into single-cell suspensions by Tumor Dissociation Kit (Miltenyi Biotech, Cat# 130-096-730) according to the manufacturer’s instructions. Briefly, total small pieces were transferred into a gentleMACS C Tube with the enzyme mix containing 2.35 mL of DMEM, 100 μL of Enzyme D, 50 μL of Enzyme R, and 12.5 μL of Enzyme A. The C Tube was tightly closed and attached to the sleeve of the gentleMACS^TM^ Octo Dissociator (Miltenyi Biotech) with the tumor isolation program. After termination of the program, the C tube was detached from the Dissociator and incubated at 37°C shaker with continuous rotation at 220 rpm for 40 min. Subsequently, repeat the tumor isolation program twice and perform a short spin up to 1,500 rpm to collect the sample at the bottom of the tube. The dissociated cells were filtered through a 70 μm cell strainer and centrifuged at 3,000 g for 5 min to remove the supernatant, and then the pellets were resuspended in red blood cell lysis buffer. Cells were recovered by adding an equal volume of 1×PBS and centrifugation at 3,000 g to remove the supernatant. Finally, cells were resuspended in PBS containing 1% FBS (v/v) and used for further analysis.

### Preparation of tumor-infiltrated lymphocytes (TILs)

The obtained single-cell suspensions of tumor-burdened lungs were centrifuged at 1,500 g for 5 min at room temperature, and the precipitants were re-suspended with 37% Percoll (Cat#17-0891-09, GE Healthcare) in 1×PBS (v/v). The suspension was centrifuged at 3,000 g for 30 min at 4℃ and the supernatant was discarded. The precipitants containing TILs were re-suspended in 1×PBS containing 1% FBS and used for further analysis.

### Cell sorting

The single-cell suspensions of tumor-burdened lungs or spleen were incubated with CD16/32 antibody for 20 min at 4℃ before surface staining with fluorescence-conjugated antibodies for 30 min at 4℃. Information of antibodies used in fluorescence-activated cell sorting (FACS) was listed in Table S6. The stromal cells (CD45.2^-^CD31^-^CD49d^+^), endothelial cells (CD45.2^-^CD31^+^) or tumor cells (CD45.2^-^EpCAM^+^CD31^-^) were separated with a BD FACSAria II cell sorter (BD Biosciences) (> 90% purity). When necessary, the cells were isolated with CD45 magnetic Nanobeads sorting (Biolegend, Cat #:480028), Dynabeads™ FlowComp™ Mouse CD8 Kit (Invitrogen, Cat# 11462D), Dynabeads™ FlowComp™ Mouse CD4 Kit (Invitrogen, Cat# 11461D) or MojoSort™ Mouse CD326 (EpCAM) Selection Kit (Biolegend, Cat# 480142) prior to cellular surface staining according to the manufacturer’s instructions.

Single-cell suspensions from spleens of OT-I mice were prepared by mechanical disruption in ice-cold 1×PBS. Cells in suspension were centrifuged for 5 min at 1,500 g. After red blood cell lysis, cells were filtrated through a 70 μm nylon mesh. Splenic CD8^+^ T cells or intratumoral CD8^+^ T cells of lung tumors were isolated using a Dynabeads™ FlowComp™ Mouse CD8 Kit (Invitrogen, Cat# 11462D) according to the manufacturer’s instructions.

### Syngeneic graft mouse model and treatment

In the syngeneic graft mouse models, CD45^-^CD31^-^EpCAM^+^ tumor cells were sorted from lung tumors of KL/KLG4^m/m^ mice that were injected with Ad-Cre followed by tamoxifen treatment. The tumor cells were cultured in DMEM supplemented with 10% FBS (v/v, Gibco) in the presence of Lip-1 (25 μM, Selleck, Cat #S7699), and the medium was changed every other day until cells outgrew and stable cell lines were formed. The cells were harvested and injected in the left back flank of C57BL/6 mice (5×10^6^/mouse, s.c.), the tumor length (L) and width (W) were measured and the tumor size was calculated as the formula: 0.5×L×W^2^.

Once the tumor volume reaches approximately to 1,500 mm^3^, the mice were euthanized and the tumors were harvested for various analyses.

To inducible knockout of GPX4 in syngeneic graft mouse models, primary CD45^-^CD31^-^EpCAM^+^ tumor cells were sorted from lung tumors of KL/KLG4^m/m^ mice (without tamoxifen treatment) after 8 weeks of Ad-Cre injection and subcutaneous transplanted into the left flanks of wild-type C57BL/6 mice. When the tumors became palpable, the mice were assigned randomly into different treatment groups. Tamoxifen was intraperitoneally injected daily at a dosage of 80 mg per kg body weight (dissolved in corn oil) and continued for 5 days and followed by intraperitoneally injected with Lip-1 (10 mg/kg) or control (as described below) once a day for 15 successive days. The tumor length (L) and width (W) were measured and the tumor size was calculated as the formula: 0.5×L×W^2^. Once the tumor volume approximates to 1500 mm^3^, the tumors were harvested and prepared for various analyses.

### Cell staining and flow cytometry analysis

The antibodies and reagents used for flow cytometry staining are summarized in Table S6. Flow cytometry protocol has been previously described^29,30^. In short, the single-cell suspensions of tumor-burdened lungs or the obtained TILs were re-suspended in 1×PBS containing 1% FBS (v/v) and blocked with anti-mouse CD16/32 antibodies for 20 min prior to staining with the antibody mixture. Surface staining was performed in PBS containing 1% FBS (v/v) at 4℃ for 30 min. For intracellular cytokine staining, cells were fixed and permeabilized with a fixation and permeabilization solution kit (Cat#424401 and 421002, respectively, Biolegend) according to the manufacturer’s instructions followed by staining with the specific antibodies against intracellular markers. For the detection of cytokine production, TILs were stimulated for 4 hours at the 37℃ incubator (5% CO_2_) in the presence of PMA (50 ng/mL, Cat#P8139, Sigma), Ionomycin (500 ng/mL, Cat#I0634, Sigma), and Golgi-stop (1:1000, Cat#554724, BD Biosciences), followed by intracellular staining.

Staining of TCF1 and TOX was performed with the True-Nuclear Transcription Factor Staining Buffer Set (Cat#424401, Biolegend) according to the manufacturer’s instructions. Subsequently, cells were fixed in 4% paraformaldehyde for 15 min at 4 ℃, and then centrifuged at 3,000 g for 5 min at 4℃ and the supernatant was discarded. The bottom cells were re-suspended in 1×PBS containing 1% FBS and used for flow cytometry analysis. Flow cytometry data were acquired on a FACSCelesta or LSRFortessaX20 flow cytometer (BD Biosciences) and analyzed by using FlowJo (v10.8.1) software (Tree Star).

### Lipid peroxidation and cell viability measurement

Experiments were performed according to the manufacturer’s protocol. Briefly, single-cell suspensions of lung tumors were cultured in blank RPMI-1640 growth medium containing 30 nM SYTOX Green Dead Cell Stain (Invitrogen) or 5 μM C11-BODIPY (lipid peroxidation sensor, Invitrogen), and incubated for 30 min at 37°C in a cell culture incubator before surface markers staining. Cells were centrifuged, washed, and resuspended in 200 μL fresh PBS followed by flow cytometry analysis within 2 hours of staining.

### Transmission electron microscopy (TEM) of cells

CD45^-^CD31^-^EpCAM^+^ tumor cells with or without iDGAT1/2 treatment were sorted as described above. After washing for three times with ice-cold PBS, the cells were lifted and pelleted at progressively increasing g forces (1,000 g for 5 min, 3,000 g for 5 min, 6,000 g for 5 min). The cell pellets were fixed in 2.5% electron microscopy grade glutaraldehyde in PBS at 4℃ for 2 hours, postfixed in 2% aqueous osmium tetraoxide, dehydrated in gradual ethanol (30-100%), propylene oxide (two cycles at 4℃ for 5 minutes) followed by infiltrated sequentially in 1:1 (v:v) propylene oxide/epoxy resin for 4 h, 1:2 (v:v) propylene oxide/epoxy resin (overnight), 100% fresh epoxy resin for 4 h. Finally, the cell pellets were embedded in 100% fresh epoxy resin and cured for 48 h at 65℃. Ultrathin sections of 50 nm were collected onto 200 mesh copper grids, and stained with Sato lead Sodium citrate for 1 min and observed using a JEM-1400 plus electron microscope operated at 100kV. For analysis, mitochondria and lipid droplets were identified by a combination of manual and automatic segmentation. The area of each mitochondria and lipid droplet identified was calculated using ImageJ v.1.52a (Bethesda) based on pixel sizes during TEM image acquisition.

### Oil red O staining

CD45^-^CD31^-^EpCAM^+^ tumor cells were isolated from lung tumors of KL and KLG4^m/m^ mice as previously described. The cells were washed twice with ice-cold PBS and then fixed in 4% formalin at room temperature for 15 minutes. After discarding the 4% formalin, the cells were washed with 60% isopropyl alcohol and dried at room temperature. Next, the cells were stained with 5% Oil Red O reagent (Beyotime, Cat#C0157M) for 10 min.

Imaging was performed using a LEICA microscope equipped with a 10×objective, and images were captured with a HAYEAR 8MP USB3.0 CMOS Video Electronic Microscope Camera (Shenzhen Hayear Electronics Co. Ltd.) using the S-EYE imaging software (Shenzhen Hayear Electronics Co. Ltd.). Three independent fields were acquired for each experimental condition, and representative images from one field of view are presented. After imaging, the Oil Red O solution was discarded, and the cells were dried at 37℃ for 1 hour.

The staining was quantified by extracting the Oil Red O stain with 100% isopropyl alcohol, and the absorbance was measured using a spectrophotometer at 510 nm.

### Immunofluorescence and confocal microscopy analysis

CD45^-^CD31^-^EpCAM^+^ tumor cells from lung tumors of KL/KLG4^m/m^ or KL/KLG4^OE/+^ were sorted as described above and washed with PBS three times. After that, the cells were incubated with 2 μM C11-bodipy solution in the dark for 30 min at 37 ℃ cell incubator and washed with PBS three times. Followed by fixing in 4% paraformaldehyde (PFA) for 15 min and washed with PBS three times. For APOE staining, the C11-bodipy labelling and 4% PFA fixed cells were permeabilized with 0.5% saponin in PBS for 5 min on ice and washed with PBS three times. Then, the cells were blocked in 1% BSA containing PBS (v/v) for 1 h and stained in blocking buffer with primary antibody overnight at 4℃. The cells were further stained with 594-conjugated secondary antibody for 1 h at 4℃. Finially, the cells were plated on slides and stained with In Situ Microplate Nuclear Stain and AntiFade (Sigma, Cat# DUO82064-1KIT), and the coverslips were mounted on slides. Images were acquired on an Olympus FV1000 fluorescence microscope.

### Delipid of the culture supernatants of tumor cells

We used a density-gradient ultracentrifugation method to remove the majority of lipoprotein classes from the culture supernatants of CD45^-^CD31^-^EpCAM^+^ tumor cells, derived from KL and KLG4^m/m^ tumors^73^. In brief, the density of the culture supernatants was increased to 1.3 g/mL through the addition of solid potassium bromide. After constructing the gradient, we centrifuged them at 27500 rpm for 24 hours at 4℃, without using a brake at the end of the run. Following this, we carefully aspirated the upper phase, which contained the lipoprotein, and transferred the lower aqueous phase to a sterile dialysis tubing with a molecular weight cutoff of 1000. Next, we immersed the dialysis tubing in 0.9% NaCl for 12 hours at 4℃, a process that was repeated four times. Finally, the lipid-free supernatants were filtered through a 0.22 μm filter and stored at -20℃, ready for use in *in vitro* cell stimulation experiments.

### *In vitro* chronic and acute stimulation of P14 CD8^+^ T cells with tumor cells-derived supernatants

CD8^+^ T cells were isolated from spleens of P14 transgenic mice (carrying a transgenic T cell antigen receptor that recognizes H-2D^b^GP_33-41_ epitope of LCMV, which were kindly provided by Prof. Liang Cheng, Wuhan University) and T-cell-depleted APCs were isolated from spleens of wild-type C57BL/6 mice and pulsed with 250 nM GP33-41 (KAVYNFATM, Synthesized by GL Biochem, Shanghai) in RPMI-1640 for 4 h at 37℃ followed by fixation in 4% paraformaldehyde and twice wash in PBS. The fixed APCs were cocultured with CD8^+^P14 cells at a 2:1 ratio (2×10^6^ APCs + 1×10^6^ P14 cells per well) in a flat-bottom 24-well plate. For chronic stimulation of T cells, at d2, d4, and d6 of coculture, CD8^+^ P14 cells were harvested from a 24-well plate and isolated via a CD8^+^ positive selection kit (Invitrogen, Cat: 11462D), replated into a 24-well plate (1×10^6^ P14 cells per well) and chronically stimulated by coculture with 50 nM GP_33-41_ pulsed APCs. Both the acute and chronic *in vitro* conditions received recombinant mouse IL-2 (10 ng/mL, Cat: CM003-20MP, CHAMOT). On day 2, cells were cultured with either tumor cells supernatants or their delipidated counterparts from KL or KLG4^m/m^ tumor cells (100 μL/well of KL and KLG4^m/m^, respectively). Finally, the cells were harvested and analyzed through flow cytometry on BD LSRFortessaX20.

### Targeted oxi-lipidomic profiling and data analysis

The experimental procedure followed the methodology outlined in previous studies^38,74^. Tumor cells (CD45^-^CD31^-^EpCAM^+^) from lung tumors of KL and KLG4^m/m^ with or without indicated treatments were sorted as previoudly detailed. Alternatively, the tumor cells culture supernatants were lyophilized prior to analysis. For each sample, 5×10^6^ cells were collected and washed once with ice-cold PBS, after centrifuging for 5 min at 1, 000 rpm, cell pellets were resuspended in 140 µL ice-cold nuclease-free water and vortexed for 10 s, added 100ng phosphatidylethanolamine (PE) (D16:1) and 100ng phosphatidylcholine (PC) (D14:1) as internal standard. Lipids were extracted using the Folch method. Briefly, 450 µL ice-cold chloroform/methanol (v/v = 2:1) containing 0.005% BHT was added to each sample, vortexed for 1 min and the sample was then incubated on ice for 15 min to enhance extraction efficiency. Finally, samples were centrifuged for 10 min at 3, 000 g, 4 °C. The organic layers (lower) were collected into new tube and dried under nitrogen using a 12-port drying manifold. Dried samples were resuspended in 60 µL of 100% LC solvent B (methanol /isopropanol solution, v/v, 3:4, containing 5 mM amide acetate). A 50 μL aliquot of the sample was transferred to a new autosampler vial for analysis.

Chromatography was performed using a HILIC HPLC column (Luna 5μm, 100 Å, 50 × 2 mm, Phenomenex) at a flow rate of 0.350mL/ min. Mass spectrometric analysis was performed in the negative ion mode using multiple-reaction monitoring (MRM) of specific precursor–product ion m/z transitions upon collision-induced dissociation. The precursor negative ions monitored were the molecular ions [M−H]^−^ for PE, and the acetate adducts [M+CH3COO]^−^ for PC. Identity was further verified by monitoring at the same time, using polarity switching, the positive molecular ions [M+H]^+^ for both PC and PE molecular species.

The product ions analyzed after collision-induced decomposition, and used for data comparison, were the carboxylate anions corresponding to the non-oxidized or oxidized arachidonoyl chains. The specific precursor–product pairs monitored in negative-ion mode and used for quantification were follows: PE(16:0e_22:5(O)), 766/345; PE(18:1a_22:4(O)), 808/347; PC (18:0a_20:4(2O)), 900/335; PC(18:0a_22:4(2O)),928/363; PE(18:0a_9’-oxo-nonanoyl), 634/171 ; PE(18:0a_5’-oxo-valeroyl), 578/115 ; PE(16:0a_22:4(O)), 782/347; PC(18:0a_HETE), 810/319^12,37,38^. Calculated oxygenated lipids were summarized in Table S1.

### Lipidomic profiling and data analyses

**Lipidomics procedures.** CD45^-^CD31^-^EpCAM^+^ tumor cells and their culture supernatants of KL and KLG4^m/m^ with or without indicated treatments were obtained as described above.

Briefly, the cells were washed with ice-cold PBS twice, for each sample, 5×10^6^ tumor cells were collected and stored at -80°C. Cell pellets were resuspended in 300 μl methanol/water (v/v=3:1), containing of LPC 18:1(d7) as the internal standard. Then, each sample was sonicated for 30 s to ensure homogeneity and added with 750 μl MTBE, vortexed for 60 s, and gently vibrated for another 30 min. After that, the sample was added with 190 μl of nuclease-free water and vortexed for 1 min. After equilibration at room temperature for 10 min, the sample was centrifuged at 14,000g for 15 min. Then 400 μl aliquot of the upper lipid extract was pipetted into the new centrifuge tube and vacuum-dried. The dried sample was dissolved with 150 μl of acetonitrile/isopropanol/water (v/v/v = 65:30:5) for the instrumental analysis in the positive and negative ion mode. The lipid profiling was acquired by using a liquid chromatography-mass spectrometry (LC-MS) system comprising ultra-performance liquid chromatography (UPLC, ExionLC™ AD, https://sciex.com.cn/) coupled to a Tandem Mass Spectrometry (MS/MS, QTRAP^®^6500+, https://sciex.com.cn/). Dissolved samples were centrifuged at 12,000 rpm for 10 min to remove residual cellular debris before injecting 2 μL onto a Thermo Accucore^TM^ C30 column (100×2.1 mm i.d., 2.6 μm). Parameters on the chromatographic separation in the positive ion mode were the same as those in the negative ion mode. The column was eluted at 80% mobile phase A (60:40 v/v acetonitrile/water, containing 0.1% formic acid and 10 mM ammonium formate) and 20% mobile phase B (10:90 v/v acetonitrile/ isopropanol, containing 0.1% formic acid and 10 mM ammonium formate) between 0 and 2 min, raised to 30% B at 2 min, increased linearly from 30% to 95% B between 2 and 17.3 min, returned to 20% B at 17.5 min, and held at 20% B until 20 min.

The flow rate was 0.35 mL/min from 0 to 20 min at 45℃. MS data were acquired using electrospray ionization (ESI) at 500℃, The spray voltage was 5.5 kV and 4.5 kV for positive and negative ionization modes, respectively. Sheath, auxiliary, and sweep gases were 45, 55 and 35 psi, respectively. Data was collected in full MS/dd-MS2 (top 5). Full MS was acquired from 100-1500 m/z with a resolution of 70,000, AGC target of 1×10^6^ and a maximum injection time of 100 ms. MS2 was acquired with resolution of 17,500, a fixed first mass of 50 m/z, AGC target of 1×10^5^ and a maximum injection time of 200 ms. Stepped normalized collision energies were 20, 30 and 40%.

### Lipidomics data processing

Raw data were processed using Analyst 1.6.3 (SCIEX) software for the detection and integration of LC-MS peaks. Lipid identities were determined based on a homemade database-MWDB (metware database) according to the retention time (RT) and pairs of precursor- and product-ion of lipids to be tested and were denoted by total number of carbons in the lipid acyl chain(s) and total number of double bonds in the lipid acyl chain(s). Lipids quantitative analysis was performed on multiple reaction monitoring (MRM) to obtain the area of LC-MS peaks in a low noise background. Peak areas were used in data reporting, data was normalized using internal standards. Negative ion mode analyses of free fatty acids and bile acids (C18-neg) were conducted similarly to that previously described.^75^ The abundance (nmol/g) of each lipid species was calculated by the following formula: X = 0.001×R×c×F×V/m, where X (nmol/g) is the concentration of the lipids, R is the ratio of peak areas of lipids to be tested related to internal standards, c is the concentration (μmol/L) of internal standards, F is the correlation factor of different internal standards, and V and m are the total volumes (μL) of the lipid extracts and weight of tumor tissues of each sample, respectively. Differential-abundance analysis was performed using ‘DEP’ R package (v 1.26.0). Calculated lipidomics datasets were summarized in Table S1.

### Bulk RNA Sequencing

RNA-sequencing and high throughput sequencing were conducted by Seqhealth Technology Co., LTD (Wuhan, China). Intratumoral CD8^+^ T cells were isolated by MACS and tumor-infiltrated tumor cells were sorted by FACS as described above, respectively. And immediately homogenized in 2 mL of TRIzol (Takara), and total RNA was extracted using RNeasy Kit (QIAGEN). RNA concentration and purity were measured by using NanoDrop 2000 (Thermo Fisher Scientific, Wilmington, DE) and RNA integrity was assessed using the RNA Nano 6000 Assay Kit of the Agilent Bioanalyzer 2100 system (Agilent Technologies, CA, USA). Poly(A) mRNA was subsequently purified from 10μg total RNA using NEBNext Oligo d(T)_25_ Magnetic Beads Isolation Module. First-strand complementary DNA was synthesized with NEBNext RNA First-Strand Synthesis Module. NEBNext Ultra II Non-Directional RNA Second Strand Synthesis Module was used for the synthesis of the complementary strand of first-strand cDNA. The resulting double-stranded DNA was purified and Vazyme TruePrep DNA Library Prep kit V2 was used to prepare libraries followed by sequencing on an Illumina Hiseq X platform with 150-bp paired-end reads strategy. Quality control of mRNA-seq data was performed by using Fatsqc (v0.11.9) and low-quality bases were trimmed by Trim_galore (0.6.4_dev). All RNA-seq data were mapped to the mouse genome (Mus_musculus_Ensemble_94) by Hisat2 (v.2.0.5) and allowed a maximum of two mismatches per read. Gene expression level was calculated by FeatureCounts (v.2.0.0) with default parameters and normalized by FPKM (Fragments Per Kilobase of exon model per Million mapped fragments). Differential expression analysis of two groups was performed using the DESeq2. Based on these differentially expressed genes, Gene Ontology (GO) and Kyoto Encyclopedia of Genes and Genomes (KEGG) enrichment analyses were implemented by using the clusterProfiler^76^ (v4.8.2) R package.

To analyze a positive or negative enrichment of the indicated pathways, we performed gene-set enrichment analyses (GSEA) with the mRNA-seq data as previously described.^30,31,70^ In such analyses, the gene sets annotated in MSigDB (Molecular Signatures Database, https://www.gsea-msigdb.org/gsea/msigdb) were listed in Table S4, and their corresponding expression values (Table S2) were analyzed through the GSEA software (http://www.gsea-msigdb.org/gsea/index.jsp).^77^

### CUT & Tag sequencing and data analysis

CUT&Tag experiment and high throughput sequencing were conducted by Seqhealth Technology Co., LTD (Wuhan, China). Briefly, CD45^-^CD31^-^EpCAM^+^ tumor cells were isolated from lung tumors of KL and KLG4^m/m^ mice as previoudly detailed. 1×10^6^ cells from each mouse (three mice in total) were harvested and following the manufacturer’s protocol of the CUT&Tag kit (Cell Signaling Technology, Cat#86652). the cells were permeabilized with digitonin and incubated with anti-H4K4me3 (Abcam, Cat# 8580) or anti-H3K27ac(Abcam, Cat# ab4729) (1:50 dilution for all antibodies) for 2 hours at 4℃. Permeabilized cells were washed and incubated with pAG-MNase enzyme, comprising the IgG-binding domain of protein A/G fused to micrococcal nuclease, for 1 hour at 4℃. MNase activity was dependent on Ca^2+^, and remianed inactive before the addition of CaCl_2_. One hour later, the permeabilized cells were washed and incubated with cold CaCl_2_ for 30 minuates at 4℃ to activate MNase, and to cleave and liberate the chromatin fragments bound by H3K4me3 and H3K27ac. The chromatin digestion reaction was then stopped, and chromatin fragments were purified with DNA purification spin columns. We performed PCR to amplify the chromatin fragments and generated libraries with the SimpleChIP ChIP-seq DNA Library Prep Kit (Cell Signaling Technology), which was compatible with the CUT&Tag kit from the same company. The library products were enriched, quantified and finally sequenced on a DNBSEQ-17 sequencer (MGI Tech Co.,Ltd.China) with PE150 model.

Quality control of sequencing data was performed by using Fatsqc (v0.12.1) and low-quality bases were trimmed by Trim_galore (v0.6.10). The trimmed reads were mapped to the mouse genome (Mus_musculus_Ensemble_94) by BWA (v0.7.15) with default parameters.

Reads that mapped to mitochondrial DNA or that had low mapping quality (<30) were excluded from downstream analysis. Next, Duplicate reads due to PCR amplification of single DNA fragments during library preparation were identified with Picard (v 2.17.3; available at http://broadinstitute.github.io/picard) and removed from downstream analysis. MACS2 (v 2.2.7.1) was used for transcription factor binding signal peak calling and the HOMER software suite (v 4.11.1) was used to assign identified peaks to genes. Multiple peaks could be assigned to a single gene as long as the peaks were located in the gene body or its promoter region (within 2.5 kb upstream). Finally, Integrative Genomics Viewer (IGV, v2.5.0) was used to display the signals of CUT&Tag sequencing.

### Public datasets mining

Single-cell RNA sequencing (scRNA-seq) data were obtained from the GEO dataset GSE207422^45^ (human NSCLC tissues) and GSE165641^31^ (KL NSCLC mouse model). Cells annotated as “tumor cells” (specifically expressed *EPCAM*, *KRT8*, and *KRT19*) in the original metadata were extracted for downstream analysis. Based on the distribution of *GPX4* expression levels, the top 25% of tumor cells were defined as the *GPX4*^High^ group, and the bottom 25% as the *GPX4*^Low^ group. The mRNA levels of *DGAT2*, *APOE*, and *GPD1L* were compared between the two groups. Statistical significance was determined using a two-tailed Student’s *t*-test. Data visualization was performed using the ggplot2 package in R (version 4.3.2).

For correlation analysis between the mean *GPX4* expression in tumor cells and the exhaustion score of T cells across patients in the datasets of GSE148071^57^, HRA002509^45^, HRA001033^58^, and GSE136246^59^, respectively. For each patient from the above dataset, tumor cells were identified based on the cell type annotations provided in the original dataset. The mean *GPX4* expression level within tumor cells of each patient was calculated. To quantify the degree of CD8⁺ T cell exhaustion from the same patients, we used the AddModuleScore function in the Seurat package (version 5.1.0) to compute exhaustion scores for T cells. The T cell exhaustion gene signature included the following genes: *PDCD1*, *HAVCR2*, *CTLA4*, *LAG3*, *TIGIT*, *ENTPD1*, *NT5E*, *BATF*, *NFATC1*, *CD38*, *EOMES*, *SLAMF6*, and *TOX*. Data visualization was performed using the ggplot2 package in R (version 4.3.2). And the relationship between the mean *GPX4* expression in tumor cells and the corresponding T cell exhaustion score across patients was evaluated using Pearson correlation analysis.

### Chromatin Immunoprecipitation (ChIP) assay

The tumor cells (CD45^-^CD31^-^EpCAM^+^) from the autochthonous and syngeneic KL/KLG4^m/m^ tumors were sorted as described above. The tumor cells were fixed with DSG (2 mM) for 30 min and 1% formaldehyde for 15 min, and then quenched by glycine. After washing three times with PBS, they were harvested in ChIP lysis buffer (50 mM Tris·HCl, pH 8.0, 0.5% SDS, 5 mM EDTA) followed by sonication to generate DNA fragments of 250-1000 bp. The lysate was centrifuged at 12,000 rpm for 10 min at 4 °C and was diluted with ChIP dilution buffer (20 mM Tris·HCl, pH 8.0, 150 mM NaCl, 2 mM EDTA, 1% Triton X-100) (4:1 volume). The resulting lysate was then incubated with Protein G agarose and anti-H3K4me3 (Cell Signaling Technology, Cat# 9751), anti-H3K27ac (Cell Signaling Technology, Cat# 8173) or control IgG at 4 °C overnight. DNA was eluted using ChIP elution buffer (0.1 M NaHCO3, 1% SDS, 30 μg/mL proteinase K) by incubation at 65°C overnight, and the DNA was purified with a DNA purification kit (TIANGEN, DP209-03). The purified DNA was assayed by quantitative PCR using the SFX connect system by a fast two-step amplification program with 2×SYBR Green Fast qPCR Master Mix (Aidlab, Cat: 342123AX). The qPCR primer for the *Dgat2* and *Gpd1l* promoter were designed according to CUT&Tag results and sequences were listed in Table S5.

### Statistical analysis

Statistical analysis was performed with GraphPad Prism 8.3.0 software by two-tailed Student’s *t*-test and one-way ANOVA unless indicated otherwise, two-way ANOVA was performed for multiple comparisons, the log-rank (Mantel-Cox) test was performed for comparing mouse survival curves. n represents the number of mice or samples used in the experiment, with the number of individual experiments listed in the legend and no statistical methods were used to predetermine sample size. Experiments in the study were repeated at least two times. Graphs show individual samples and center values indicate the mean. *P* values < 0.05 were considered significant (*:*P* < 0.05; **:*P* < 0.01; \*\*\**P* < 0.001, \*\*\*\**P* < 0.0001; ns stands not significant, *P* values > 0.05). Graphs show mean ± SEM of different mice or samples unless indicated otherwise. A hypergeometric test was performed for Gene pathway enrichment analysis. GSEA was used to rank the probes and analyze the enrichment based on t statistics (http://www.broadinstitute.org/gsea/). The heatmap of the signature genes was generated in GraphPad Prism 8.3.0 software based on their relative expression levels, which were normalized to their mean and then divided by their variance.

### Data and code availability

The RNA-seq data and CUT&Tag data, which support the findings of this study, have been deposited in the Gene Expression Omnibus (GEO) under accession codes GSE281154 and GSE287544, respectively, which can be found in Table S2 and S3.

## Acknowledgments

We thank Drs. Hongbin Ji (Chinese Academy of Sciences, Shanghai) for the Ad-SPC-Cre virus and the 3T3LSL-RFP cells, Liang Cheng (Wuhan University) for P14 cells, Yan Wang (Wuhan University) for the delipidated fetal bovine serum and the critical reading of the manuscript, and members of the Zhong laboratory and the core facilities of Medical Research Institute and College of Life Sciences for technical help. This study was supported by grants from the National Key Research and Development Program of China (Grant Nos. 2023YFC2306100, 2024YFA1803103, and 2024ZD0524901), the Natural Science Foundation of China (Grant Nos. 82425027, 32470960, 32270951, and 82302072), the Fundamental Research Funds for the Central Universities (Grant Nos. 2042022dx0003 and 2042025kf0044), the Natural Science Foundation of Hubei (2025AFA026), and the Natural Science Foundation of Wuhan (2024040701010031). P.W. was supported by the Postdoctoral Fellowship from the China Postdoctoral Science Foundation (2023M732708) and Hubei Province.

## Author contributions

B.Z., D.L., and J.Y. designed and supervised the study; H.Y.D. and F.Z. helped with the analysis of NSCLC scRNA-seq data; P.W. and S.Z. performed the experiments; X.C. performed lipidomics and Oxi-lipidomic profiling; X.-D.Y. and H.S. helped with mouse breeding, genotyping, and cell sorting; H.Y. provided reagents for the lipidomics; and B.Z., D.L., and P.W. wrote the paper. All the authors analyzed the data.

---

**Figure S1.**
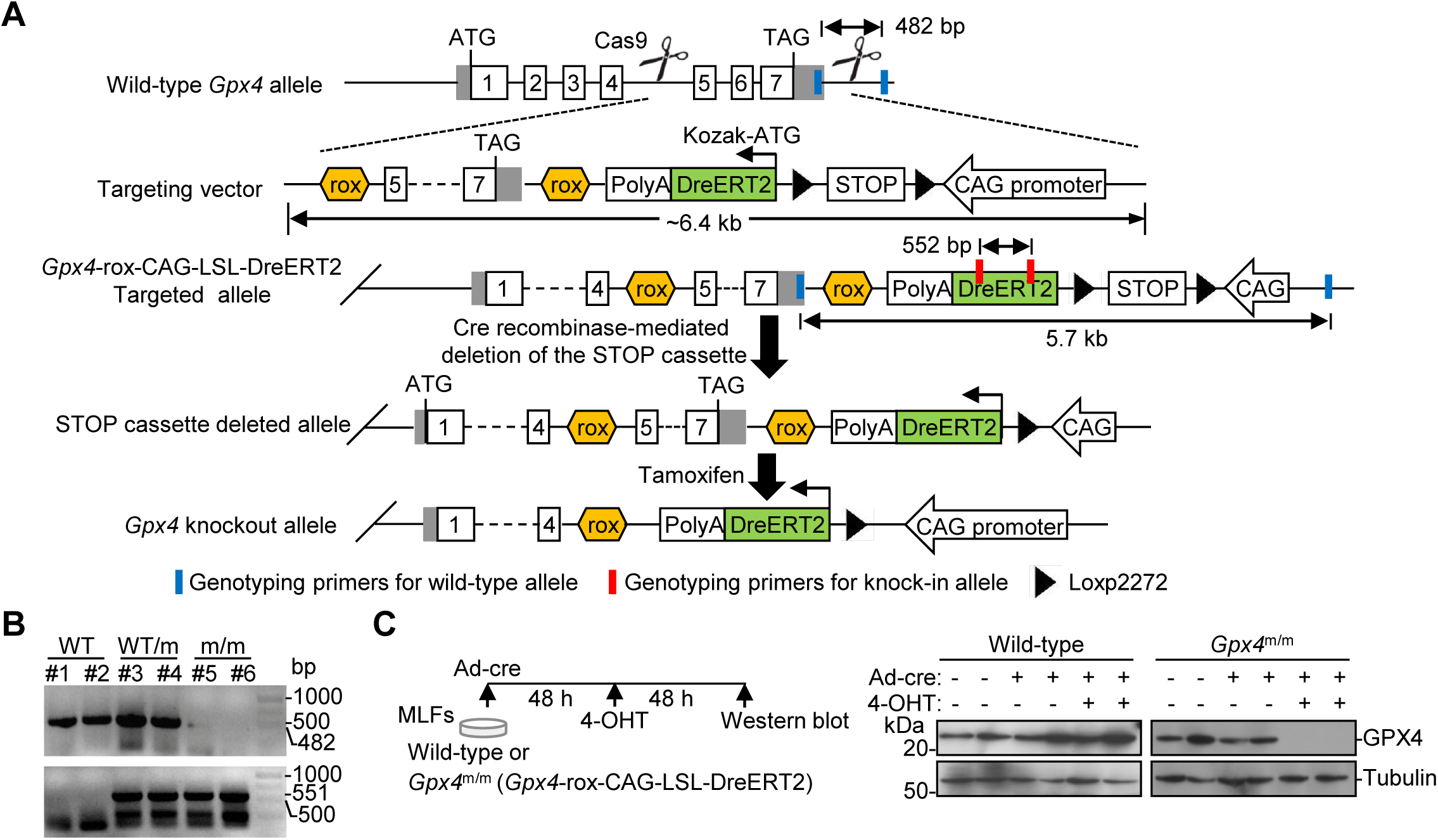
The generation of *Gpx4*^m/m^ mice. (A) A scheme of the generation of the *Gpx4*-rox-CAG-LSL-DreERT2 (*Gpx4*^m/m^) mice. (B) PCR analysis of the tail genomic DNAs from wild-type, *Gpx4*^m/+^, *Gpx4*^m/m^ mice. (C) Experimental design (left scheme) and immunoblot (right panels) analysis of GPX4 in wild-type and *Gpx4*^m/m^ MLFs infected with Ad-Cre (Ad-Cre) for 48 hours followed by treatment with 4-hydroxytamoxifen (4OHT, 1μM) for 48 hours. Data are representative results of two independent experiments (B and C).

**Figure S2.**
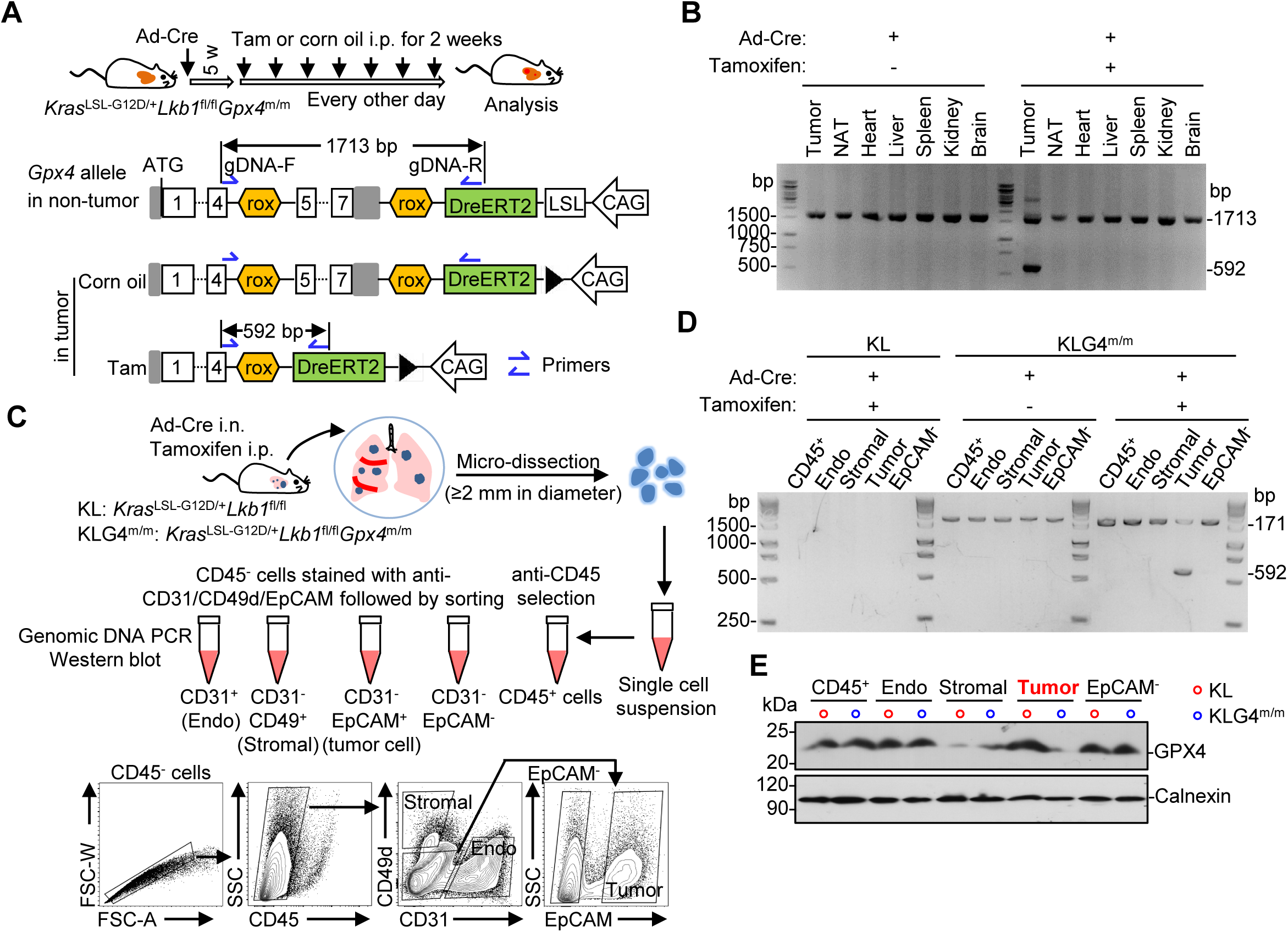
Tamoxifen-mediated knockout of GPX4 in tumor cells of *Kras*^LSL-G12D^*Lkb1*^fl/fl^*Gpx4*^m/m^ (KLG4^m/m^) mice. (A) *In vivo* validation of the recombination of *Gpx4* gene locus in lung tumors of tumor-bearing *Kras*^LSL-G12D^*Lkb1*^fl/fl^*Gpx4*^m/m^ (KLG4^m/m^) mice. The KLG4^m/m^ mice were intranasally injected with Ad-Cre (3×10^6^ PFU per mice) for 5 weeks followed by intraperitoneal injection of either tamoxifen (Tam, 80 mg/kg, resolved in corn oil) or corn oil every other day for 2 weeks for various analyzes (upper). A scheme of the *Gpx4* genomic locus in non-tumor or in Tam- or corn oil-treated tumor tissues of KLG4^m/m^ mice (bottom). (B) PCR analysis of genomic DNAs from different tissues from the Ad-Cre-infected Tam- or corn oil-treated KLG4^m/m^ mice as in (A). (C) Validation of tumor cell-specific recombinant of the *Gpx4* gene locus from lung tumors of KL and KLG4^m/m^ mice treated as in (A). A scheme of magnetic-activated cell sorting (MACS) to obtain CD45^+^ cells and fluorescence-activated cell sorting (FACS) to obtain endothelial cells, stromal cells, and tumor cells form the CD45^-^ population (upper) and the representative gating images for cell sorting (bottom) were shown here. (D-E) PCR (D) and immunoblot (E) analysis of the indicated cells from lung tumors of KL and KLG4^m/m^ mice obtained in (C) to determine the knockout efficiency and specificity of GPX4. Data are representative results of two independent experiments (B-E).

**Figure S3.**
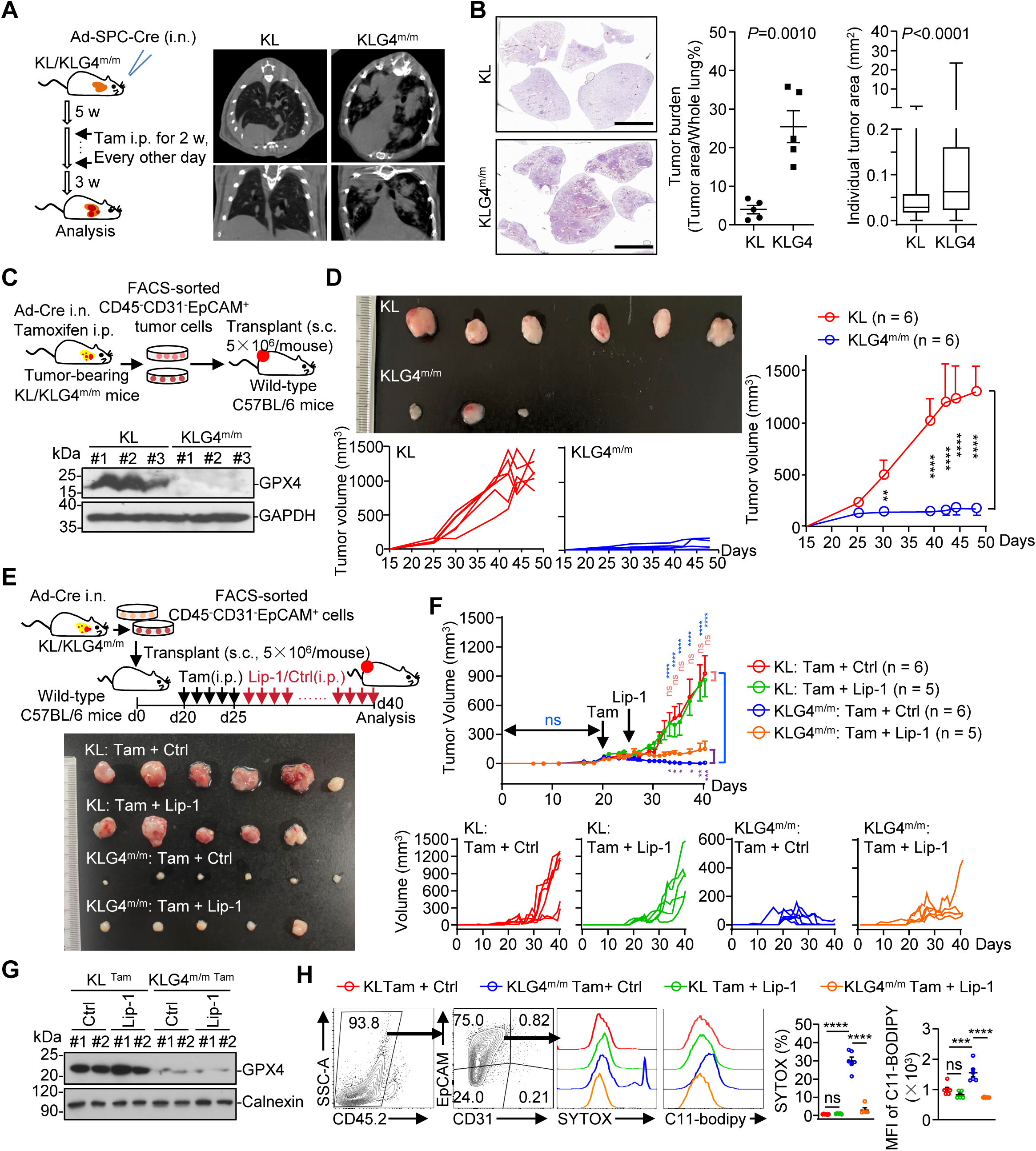
**Tumor cell-specific deficiency of GPX4 inhibits tumor growth in syngeneic graft mouse models.** (A) KL (n = 5) and KLG4^m/m^ (n = 5) mice were intranasally injected with Ad-SPC-Cre (5×10^6^ PFU per mouse) for 5 weeks followed by intraperitoneal injection of tamoxifen every other day for 2 weeks (left scheme). The mice were rested for another 3 weeks followed by micro-CT imaging analysis (right images). (B) Representative images of HE staining (left) and statistics of tumor burdens (middle) and individual tumor sizes (right) of tumor-burdened lungs from the KL (n=5) and KLG4^m/m^ (n=5) mice treated as described in (A). (C) The CD45^-^CD31^-^EpCAM^+^ tumor cells were FACS-sorted from lung tumors of KL and KLG4^m/m^ mice (with Ad-Cre i.n. infection followed by tamoxifen i.p. treatment) and subcutaneously transplanted into the flanks of wild-type C57BL/6 mice for further analysis (upper scheme). Immunoblot analysis of GPX4 and GAPDH in the subcutaneous KL or KLG4^m/m^ tumors (lower panels). (D) Images of the transplanted tumors as described in (C) (left top). Individual tumor growth curves (left bottom) and the overall tumor growth curves (right) of the subcutaneous KL and KLG4^m/m^ (n = 6 in each group). (E) The CD45^-^CD31^-^EpCAM^+^ tumor cells were sorted from lung tumors of KL and KLG4^m/m^ mice (with Ad-Cre i.n. infection only) for10 weeks and transplanted subcutaneously into the flanks of wild-type C57BL/6 mice. When the tumors became palpable, the mice were intraperitoneal injected with tamoxifen (80 mg/kg in corn oil) for 5 successive days followed by daily intraperitoneal injection of Liprostatin-1 (Lip-1, 10 mg/kg) or the control dissolvent (Ctrl) for 15 consecutive days followed by various analyses (upper scheme). The individual tumors were imaged at the end of the study (lower image). (F) The overall tumor growth curves (top) and the individual tumor growth curves (bottom) for KL (n = 6 for Ctrl and n = 5 for Lip-1) or KLG4^m/m^ (n = 6 for Ctrl and n = 5 for Lip-1) subcutaneous tumors as described in (E). (G) Immunoblot analysis of GPX4 in the for KL (n = 2 for Ctrl and Lip-1) or KLG4^m/m^ (n = 2 for Ctrl and Lip-1) subcutaneous tumors as described in (E). (H) Flow cytometry analysis (left flow charts) and the percentages of cell death and lipid peroxidation (right graphs) of CD45^-^CD31^-^EpCAM^+^ tumor cells from KL (n = 6 for Ctrl and n = 5 for Lip-1) or KLG4^m/m^ (n = 6 for Ctrl and n = 5 for Lip-1) subcutaneous tumors as described in (E). Graphs show mean ± SEM (B, D, F, and H). \**P*<0.05, \*\**P*<0.01, *** *P* < 0.001, **** *P* <0.0001, ns: not significant (two-tailed student’s *t*-test for B or two-way ANOVA for D, F, and H). Scale bars represent 5 mm (A and B).Data are representative results of two independent experiments (A-H).

**Figure S4.**
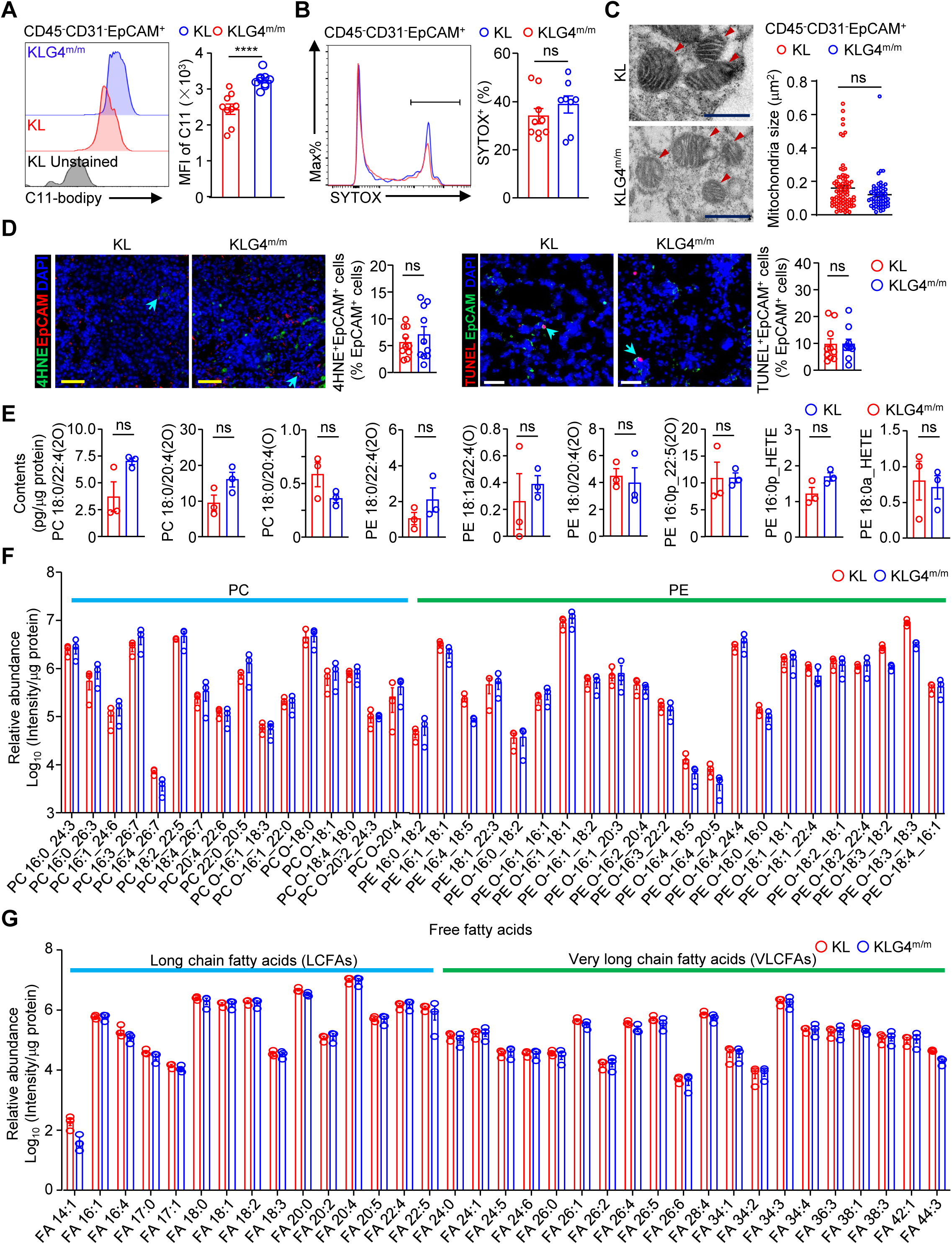
**Tumor cell-specific knockout of GPX4 in the KL autochthonous mouse model does not result in ferroptosis of tumor cells.** (A) Flow cytometry analysis of C11-bodipy staining of CD45^-^CD31^-^EpCAM^+^ tumor cells from the lung tumors of KL (n = 9) and KLG4^m/m^ (n = 9) mice that were intranasally injected with Ad-Cre (2×10^6^ PFU per mice) for 5 weeks followed by intraperitoneal injection of either tamoxifen (Tam, 80 mg/kg, resolved in corn oil) or corn oil every other day for 2 weeks and rest for 3 weeks. (B) Flow cytometry analysis of SYTOX staining of CD45^-^CD31^-^EpCAM^+^ tumor cells from the lung tumors of KL (n = 9) and KLG4^m/m^ (n = 8) mice treated as in (A). (C) Representative images of transmission electron microscopy analysis (left) and quantification of mitochondrial sizes (right) of CD45^-^CD31^-^EpCAM^+^ tumor cells from the lung tumors of KL (n = 85 mitochondria from 15 cells) and KLG4^m/m^ (n = 56 mitochondria from 10 cells) mice treated as in (A). Arrowheads indicated mitochondria. (D) Representative images and quantitative results of immunofluorescence staining of EpCAM (red) and 4HNE (green) (left panel) or EpCAM (green) and TUNEL (red) (right panel) in lung tumors from KL (n = 10) and KLG4^m/m^ (n = 10) mice treated as in (A). Light blue arrows indicated cells co-stained with DAPI, EpCAM and 4HNE (left) or DAPI, EpCAM and TUNEL (right). (E) Quantification of the indicated peroxided phospholipid (mainly including PE and PC) in the CD45^-^CD31^-^EpCAM^+^ tumor cells of KL (n = 3) and KLG4^m/m^ (n = 3) mice treated as in (A). (F, G) Relative quantification of PC and PE (F) or free fatty acid (FFA) (G) in CD45^-^CD31^-^EpCAM^+^ tumor cells of KL (n = 3) and KLG4^m/m^ (n = 3) mice treated as in (A). Graphs show mean ± SEM (A-H). Scale bars represent 500 nm (C), 50 μm (D, yellow) or 25 μm (D, white). *** *P* < 0.001, ns: not significant (two-tailed student’s *t*-test for A-E and multiple *t*-test for F and G). Data are representative results of two independent experiments (A-D).

**Figure S5.**
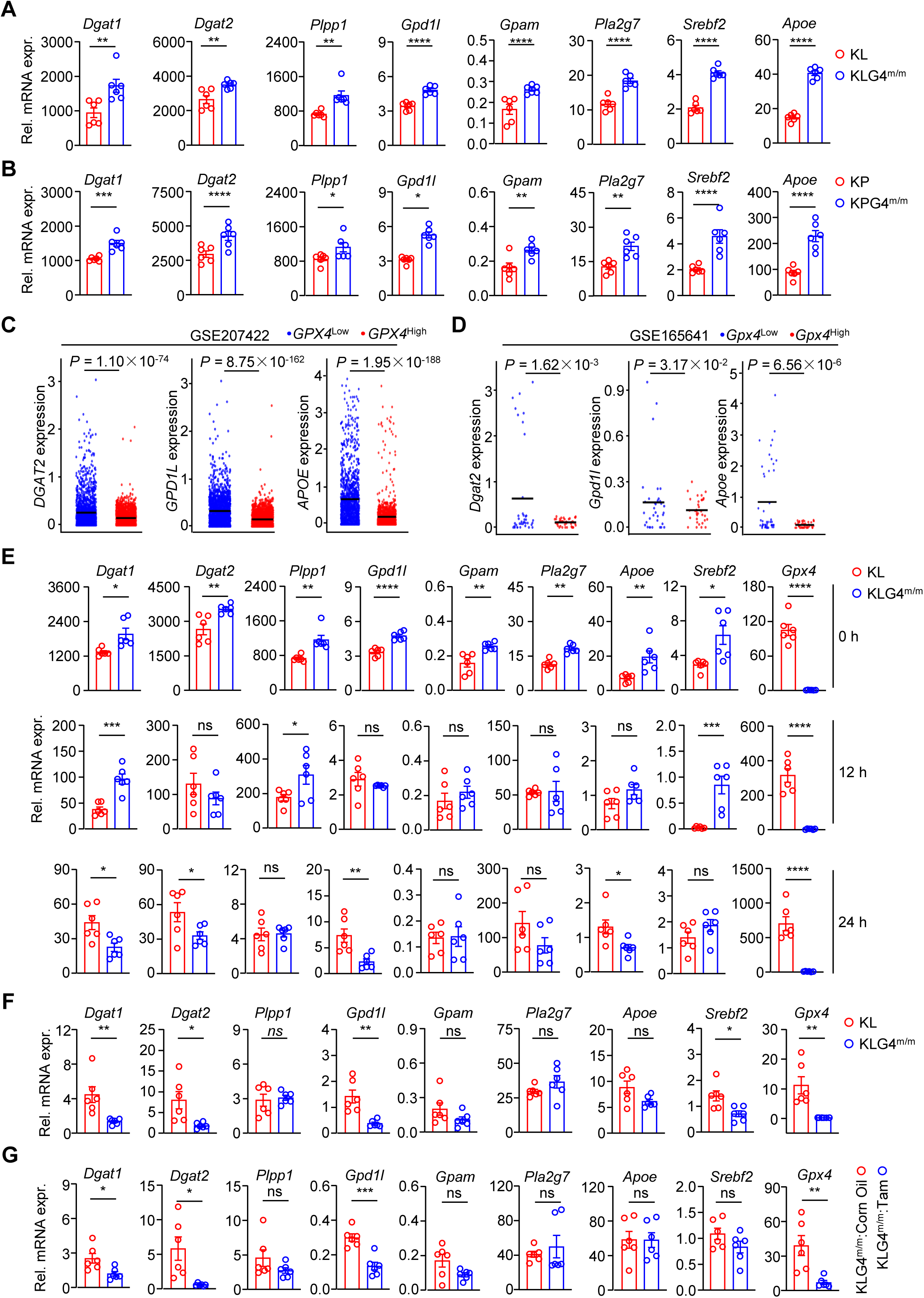
**Expressional patterns of TAG metabolism related genes across different contexts.** (A) Relative mRNA levels of the indicated genes involved in TAG metabolism in CD45^-^CD31^-^EpCAM^+^ tumor cells from lung tumors of KL (n = 6) and KLG4^m/m^ mice (n = 6) that were intranasally injected with Ad-Cre (2×10^6^ PFU per mouse) for 5 weeks followed by intraperitoneal injection of tamoxifen every other day for 2 weeks. The mice were rested for another 3 weeks followed by RT-qPCR analysis. (B) Relative mRNA levels of the indicated genes involved in TAG metabolism in CD45^-^CD31^-^EpCAM^+^ tumor cells from lung tumors of KP (n = 6) and KPG4^m/m^ mice (n = 6) that were intranasally injected with Ad-Cre (2×10^6^ PFU per mouse) for 5 weeks followed by intraperitoneal injection of tamoxifen every other day for 2 weeks. The mice were rested for another 3 weeks followed by RT-qPCR analysis. (C) Scatter plots show the mRNA levels of *DGAT2*, *GPD1L*, and *APOE* between *GPX4*^Low^ (bottom 25%,blue) and *GPX4*^High^ (top 25%, red) tumor cells in the human NSCLC tissues (GEO:207422). (D) Scatter plots show the mRNA levels of *Dgat2*, *Gpd1l*, and *Apoe* between *Gpx4*^Low^ (bottom 25%,blue) and *Gpx4*^High^ (top 25%, red) tumor cells in the KL NSCLC mouse models (GEO:165641). (E) Relative mRNA levels of the indicated genes involved in TAG metabolism in CD45^-^CD31^-^EpCAM^+^ tumor cells from lung tumors of KL (n = 6) and KLG4^m/m^ (n = 6) mice treated as in (A) that were cultured them *in vitro* for 0, 12, 24 hours followed by RT-qPCR analysis. (F) Relative mRNA levels of the indicated genes involved in TAG metabolism in subcutaneous syngeneic KL and KLG4^m/m^ tumors. The CD45^-^CD31^-^EpCAM^+^ tumor cellss (5×10^6^/mouse) sorted from lung tumors of Ad-Cre-infected tamoxifen-treated KL or KLG4^m/m^ mice treated as in (A) were subcutaneously inoculated into the flanks of wild-type C57BL/6 mice (n = 6 for each group) for 50 days followed by RT-qPCR analysis. (G) Relative mRNA levels of the indicated genes involved in TAG metabolism in Tam-treated subcutaneous syngeneic KL and KLG4^m/m^ tumors. The CD45^-^CD31^-^EpCAM^+^ tumor cells (5×10^6^/mouse) sorted from Ad-Cre-infected KLG4^m/m^ mice were subcutaneously inoculated into the flanks of wild-type C57BL/6 mice for 3 weeks (when the tumors were palpable) followed by one-weak tamoxifen treatment (80 mg/kg body weight daily) and RT-qPCR analysis (n = 6 each group). Graphs show mean ± SEM (A-G). * *P* < 0.05, ** *P* < 0.01, *** *P* < 0.001, **** *P* <0.0001, ns: not significant (two-tailed student’s *t*-test). Data are representative results of two independent experiments (A-B and E-G).

**Figure S6.**
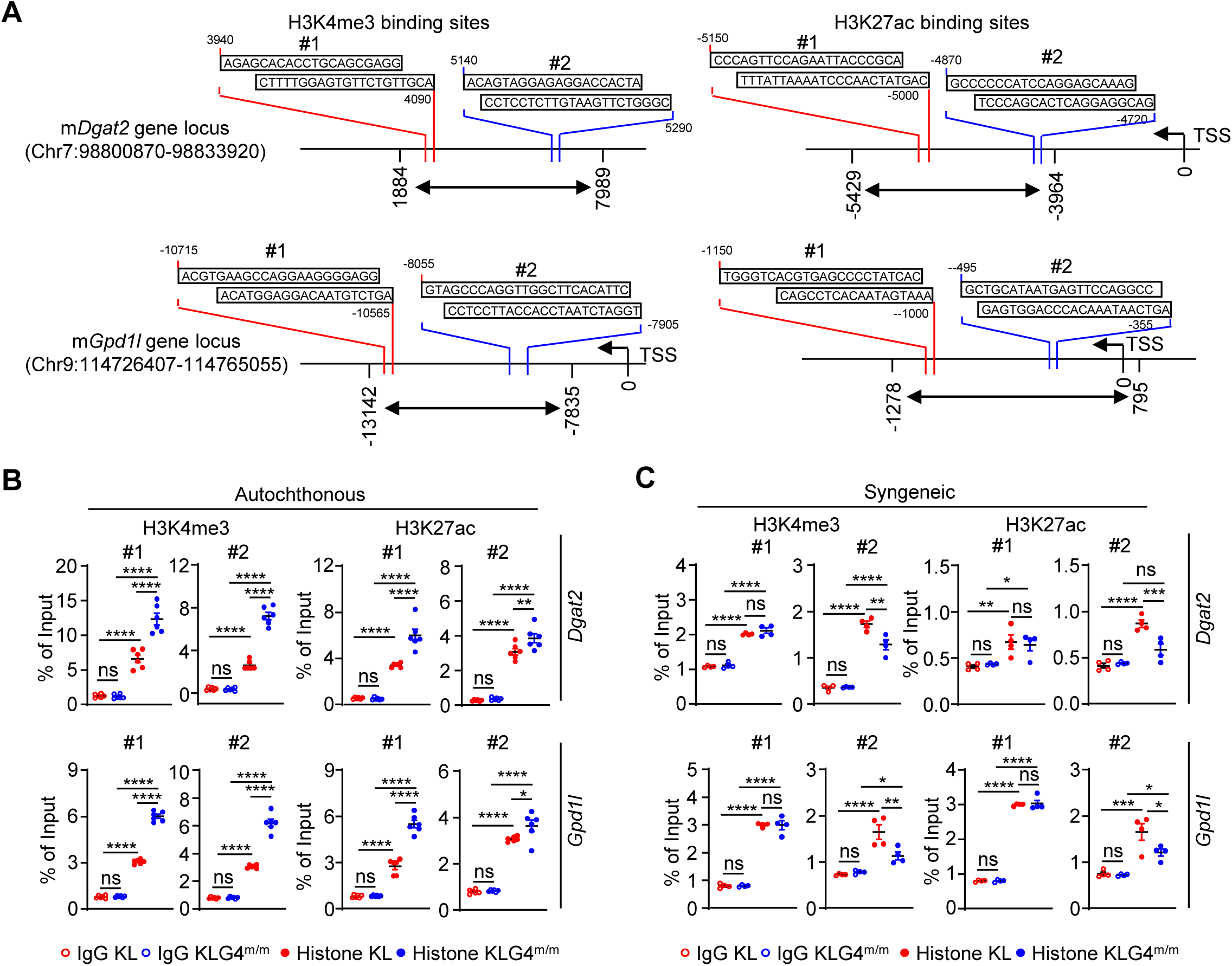
H3K4me3 and H3K27ac modifications on mouse *Dgat2* loci. (A) A scheme of mouse *Dgat2* loci (top) and *Gpd1l* loci (bottom) that contained H3K4me3 (left) and H3K27ac (right) modifications. (B) Chromatin immunoprecipitation qPCR (ChIP-qPCR) assay of H3K4me3 and H3K27ac modifications on the *Dgat2* loci (top) and *Gpd1l* loci (bottom) in CD45^-^CD31^-^EpCAM^+^ tumor cells from lung tumors of Ad-Cre-infected Tam-treated KL (n = 6) and KLG4^m/m^ (n = 6) mice. (C) ChIP-qPCR assay of H3K4me3 and H3K27ac modifications on the *Dgat2* loci (top) and *Gpd1l* loci (bottom) in Tam-treated subcutaneous syngeneic KL (n = 4) and KLG4^m/m^ (n = 4) tumors. The CD45^-^CD31^-^EpCAM^+^ tumor cells from lung tumors of Ad-Cre-infected KL and KLG4^m/m^ mice were subcutaneously inoculated into the flanks of wild-type C57BL/6 mice for 3 weeks (when the tumors were palpable) followed by one-weak tamoxifen treatment (80 mg/kg body weight daily) and ChIP-qPCR analysis. Graphs show mean ± SEM. * *P* < 0.05, ** *P* < 0.01, **** *P* <0.0001, ns: not significant (one-way ANOVA). Data are representative results of two independent experiments.

**Figure S7.**
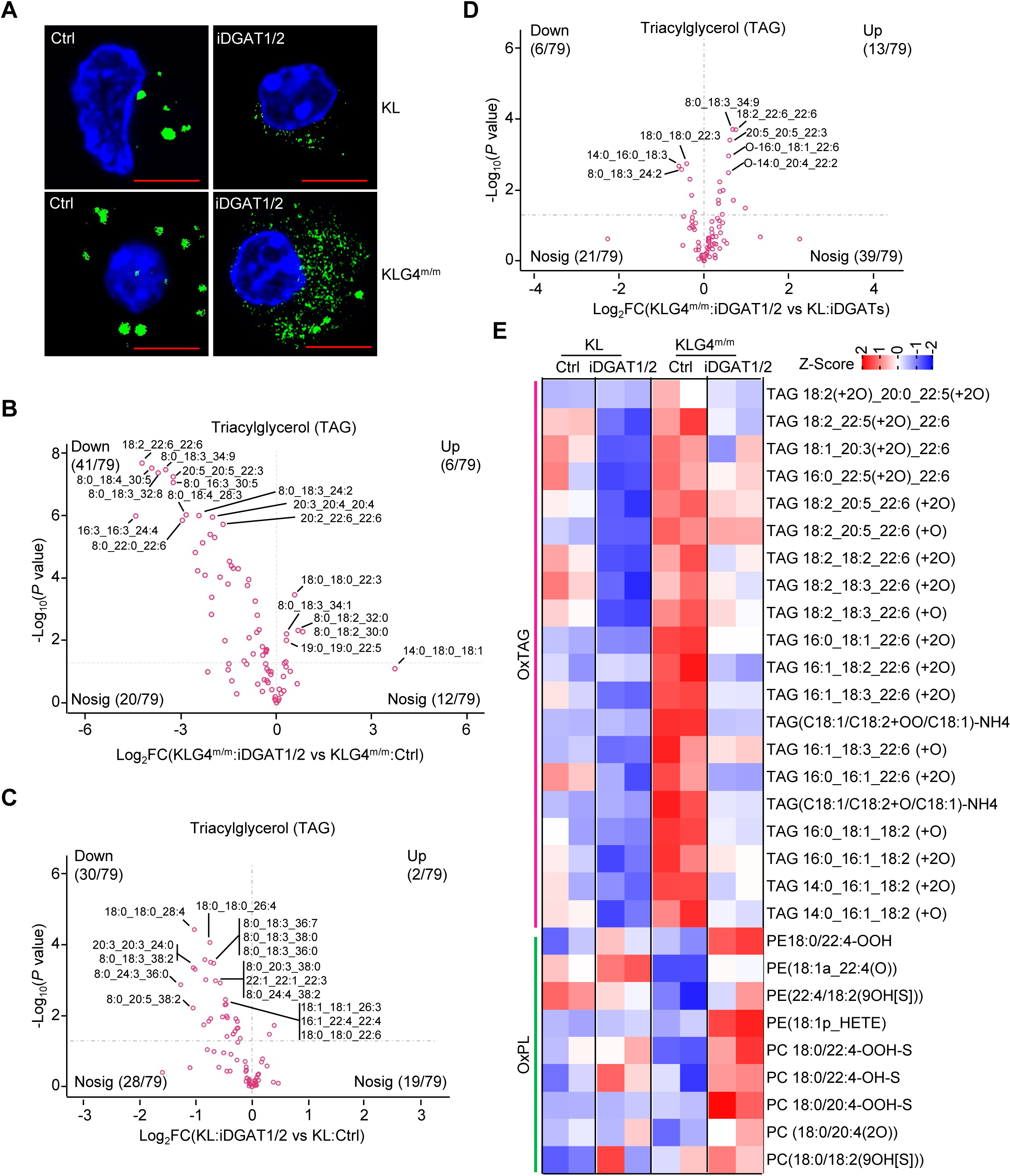
**Inhibition of DGAT1/2 reprograms PUFA-PE/PC and oxPE/PC accumulation in KL and KLG4^m/m^ tumor cells.** (A) Representative images of CD45^-^CD31^-^EpCAM^+^ tumor cells from lung tumors of KL and KLG4^m/m^ mice with or without iDGAT1/2 treatment. The KL and KLG4^m/m^ mice that were intranasally injected with Ad-Cre (2×10^6^ PFU per mouse) for 5 weeks followed by intraperitoneal injection of tamoxifen every other day for 2 weeks. One week after Tam treatment, the mice were injected with iDGAT1/2 (composing of T863 and PF06424439, 20 mg and 40 mg per kg body weight, respectively) every other day by gavage for 5 weeks. The mice were rest for one week followed by subsequent analyses. (B) Volcano plot showing the levels of triacylglycerols (TAGs) in CD45^-^CD31^-^EpCAM^+^ tumor cells derived from iDGAT1/2 treated KLG4^m/m^ mice (n = 2) versus Control treated KLG4^m/m^ mice (n = 2) treated as in (A) under indicated treatment (C) Volcano plot showing the levels of triacylglycerols (TAGs) in CD45^-^CD31^-^EpCAM^+^ tumor cells derived from iDGAT1/2 treated KL mice (n = 2) versus Control treated KL mice (n = 2) treated as in (A) under indicated treatment. (D) Volcano plot showing the levels of triacylglycerols (TAGs) in CD45^-^CD31^-^EpCAM^+^ tumor cells derived from iDGAT1/2 treated KLG4^m/m^ mice (n = 2) versus iDGAT1/2 treated KL mice (n = 2) treated as in (A) under indicated treatment. (E) Heat map showing the changes of classical PUFA-oxTAG and ferroptosis-related oxidized PC and PE in tumor cells from KL and KLG4^m/m^ mice with or without iDGAT1/2 treatment. Rows represent Z-score normalized intensities, columns represent samples, color-coded from red (high intensity) to blue (low intensity).

**Figure S8.**
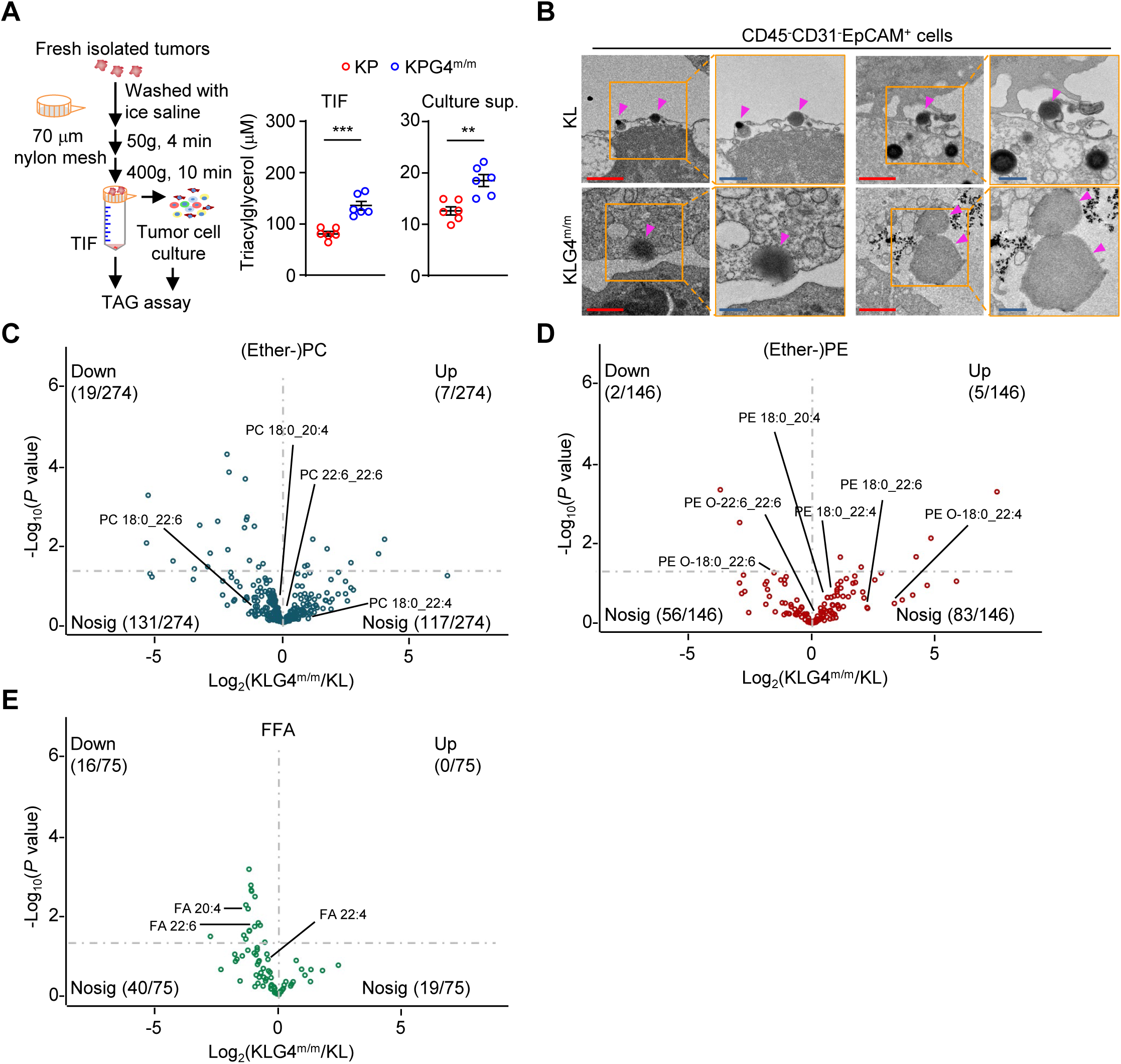
**Quantification of the lipids released from tumor cells.** (A) A scheme of tumor interstitial fluid (TIF) and tumor cells (CD45^-^CD31^-^EpCAM^+^) cultured supernatants collection (left) and quantification of the TAG (right) in TIF and tumor cells cultured supernatants from lung tumors of KP (n = 6) and KPG4^m/m^ (n = 6) mice that were intranasally injected with Ad-Cre (2×10^6^ PFU per mouse) for 5 weeks followed by intraperitoneal injection of tamoxifen every other day for 2 weeks. The mice were allowed to rest for another 3 weeks followed by TAG assessment. (B) Representative images of transmission electron microscopy of CD45^-^CD31^-^EpCAM^+^ tumor cells from the lung tumors of KL (n = 4) and KLG4^m/m^ (n = 4) mice that were intranasally injected with Ad-Cre (2×10^6^ PFU per mice) for 5 weeks followed by intraperitoneal injection of either tamoxifen (Tam, 80 mg/kg, resolved in corn oil) or corn oil every other day for 2 weeks and rest for 3 weeks. The orange boxed areas were shown at a higher magnification on the right. (C-E) Lipidomic profile of phosphatidylcholine (PC) and ether-linked PC (ether-PC) (B), phosphatidylethanolamine (PE) and ether-linked PE (ether-PE) (C), and free fatty acid (FFA) (D) in the cultured supernatants of CD45^-^CD31^-^EpCAM^+^ tumor cells from lung tumors of KL (n = 5) and KLG4^m/m^ (n = 5) mice treated as in (B). Scale bars represent 1 μm (red, B) and 500 nm (blue, B). Data are representative results of two independent experiments (A-B).

**Figure S9.**
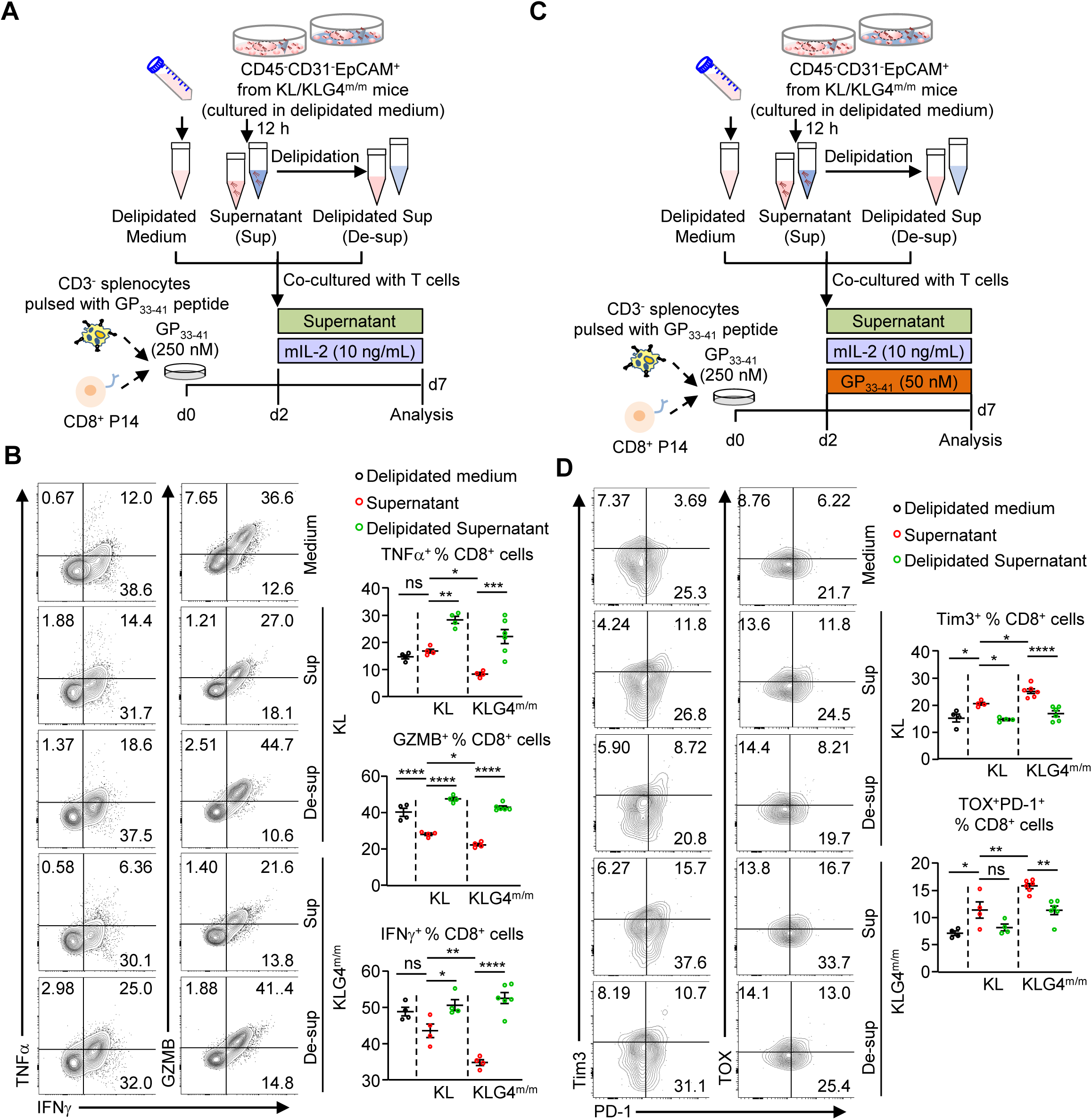
**KPG4^m/m^ tumor cell-secreted lipid mediators promote dysfunction and exhaustion of CD8^+^ T cells.** (A-B) A scheme of acute *in vitro* activation (A) and representative flow charts (B, left) and quantitative analysis (B, right) of P14 cells (n = 4 or 6 for each group) in the presence of the supernatants or the delipidated supernatants from KL or KLG4^m/m^ CD45^-^CD31^-^EpCAM^+^ tumor cells. (C-D) A scheme of chronic *in vitro* activation (C) and representative flow charts (D, left) and quantitative analysis (D, right) of P14 cells (n = 4 or 6 for each group)in the presence of the supernatants or the delipidated supernatants from KL or KLG4^m/m^ CD45^-^CD31^-^EpCAM^+^ tumor cells. Graphs show mean ± SEM (B and D). * *P* < 0.05, ** *P* < 0.01, *** *P* < 0.001, **** *P* <0.0001. ns: not significant (two-way ANOVA for B and D). Data are representative results of two independent experiments (B and D).

**Figure S10.**
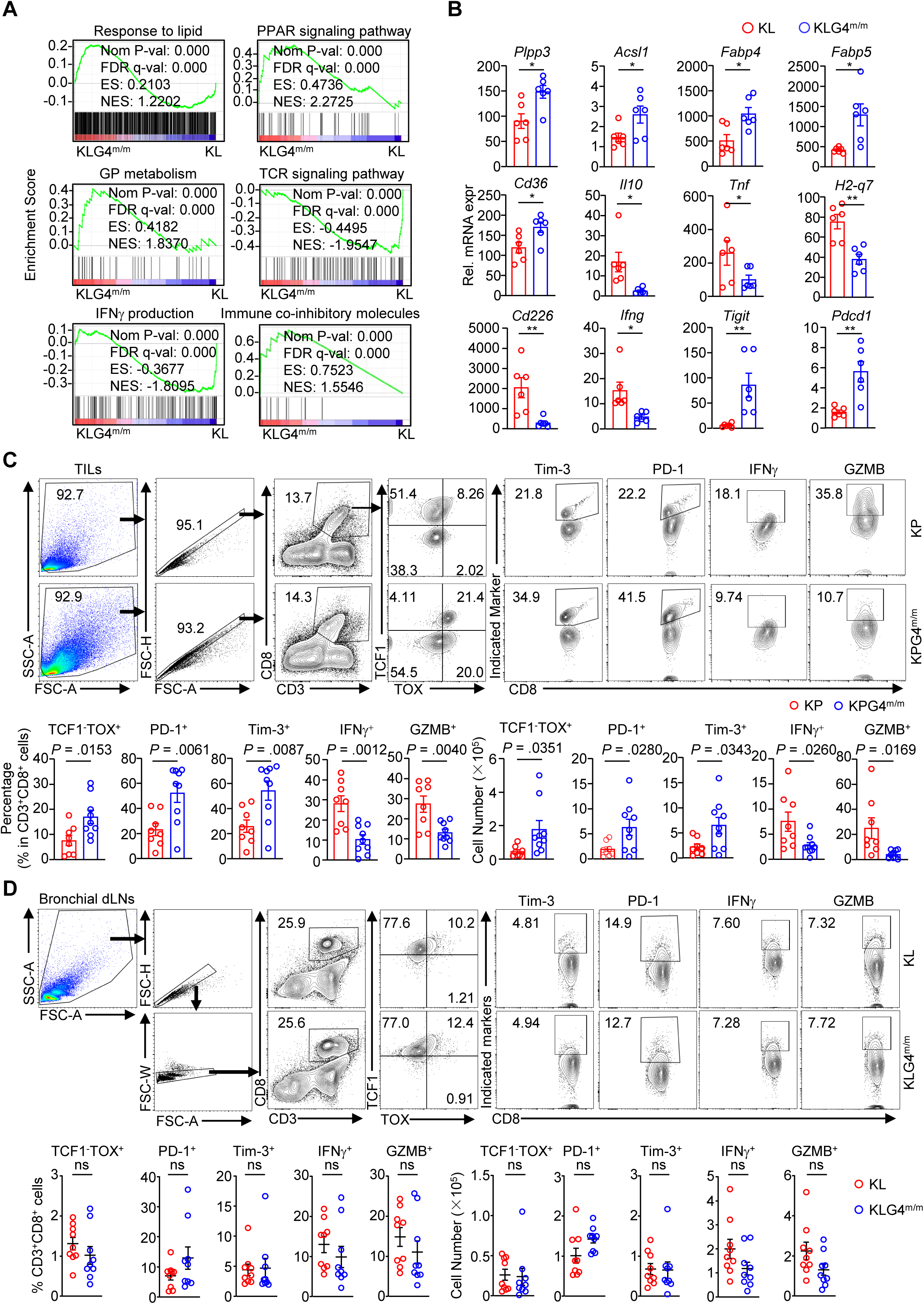
**KPG4^m/m^ tumor cell-secreted lipid mediators promote dysfunction and exhaustion of CD8^+^ T cells.** (A) GSEA analysis of the response to lipid, PPAR signaling pathway, Glyerophospholipid metabolism, TCR signaling pathway, IFNγ production and immune co-inhibitory molecules based on the transcriptome data of CD8^+^ TILs from lung tumors of KLG4^m/m^ (n = 2) and KL (n = 2) mice that were intranasally injected with Ad-Cre (2×10^6^ PFU per mice) for 5 weeks followed by intraperitoneal injection of either tamoxifen (Tam, 80 mg/kg, resolved in corn oil) or corn oil every other day for 2 weeks and rest for 3 weeks. (B) RT-qPCR analysis of the signature genes involved in lipid metabolism and T cells response in CD45^-^CD31^-^EpCAM^+^ tumor cells from lung tumors of KL (n = 6) and KLG4^m/m^ (n = 6) mice treated as in (A). (C) Representative flow cytometry images (upper charts) and quantification analysis (lower graphs) of tumor-infiltrated lymphocytes (TILs) from lung tumors of KP (n = 8) and KPG4^m/m^ (n = 9) mice that were intranasally injected with Ad-Cre (2 × 10^6^ PFU per mouse) for 5 weeks followed by intraperitoneal injection of tamoxifen (Tam, 80 mg/kg, resolved in corn oil) every other day for 2 weeks and rest for 3 weeks. (D) Representative flow cytometry images (upper charts) and quantification analysis (lower graphs) of bronchial draining lymph nodes (dLNs) from lung tumors of KL (n = 9) and KLG4^m/m^ (n = 9) mice that were intranasally injected with Ad-Cre (2 × 10^6^ PFU per mouse) for 5 weeks followed by intraperitoneal injection of tamoxifen (Tam, 80 mg/kg, resolved in corn oil) every other day for 2 weeks and rest for 3 weeks. Graphs show mean ± SEM (B, lower graphs of C and D). * *P* < 0.05, ** *P* < 0.01, *** *P* < 0.001, **** *P* <0.0001. ns: not significant (two-tailed student’s *t*-test for B and lower graphs of C and D). Data are representative results of two independent experiments (B and lower graphs of C and D).

**Figure S11.**
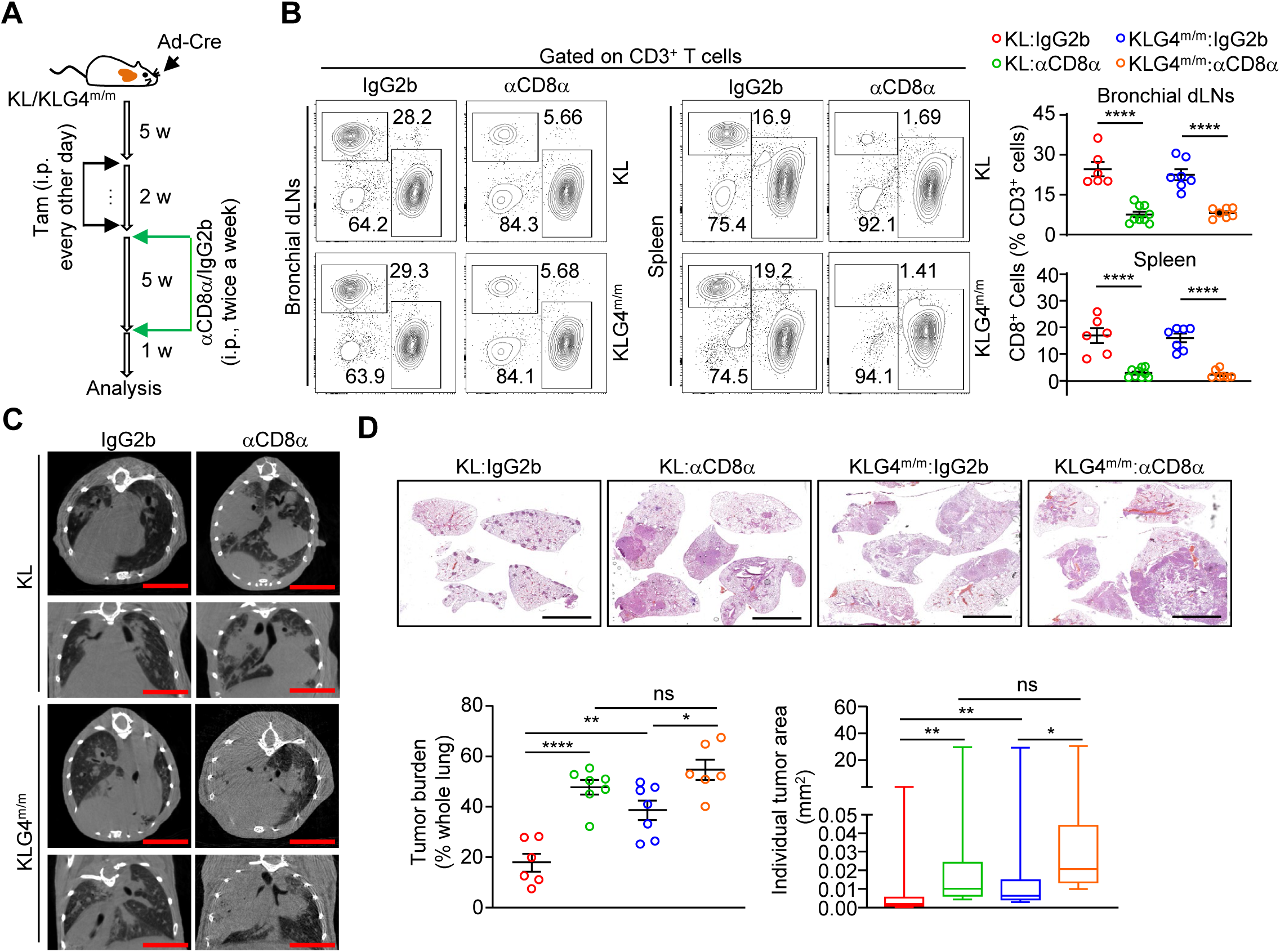
**Depletion of CD8^+^ T cells accelerates tumor progression of tumor-bearing KL/KLG4^m/m^ mice.** (A) Schematic illustration depicts tumor induction and CD8^+^ T cells depletion (anti-mouse CD8α, αCD8α, Clone:2.43) in KL and KLG4^m/m^ mice that were intranasally injected with Ad-Cre (2×10^6^ PFU per mouse) for 5 weeks followed by intraperitoneal injection of tamoxifen every other day for 2 weeks. After completion of tamoxifen treatment, the mice were intraperitoneally injected with αCD8α or IgG2b (200 μg per injection, respectively) twice a week for 5 weeks. The mice were rest for one week followed by various analyses. (B) Representative flow cytometry images (left charts) and quantification analysis (right graphs) of the percentage of CD8^+^ T cells in bronchial draining lymph nodes (top-right of B) or spleen (bottom-right of B) from the tumor-bearing KL (n=9 for αCD8α and n=6 for IgG2b, respectively) and KLG4^m/m^ (n=7 for αCD8α and n=7 for IgG2b, respectively) mice treated as described in (A). (C) Representative images of micro-CT of tumor-burdened lungs from the KL (n=7 for αCD8α and n=6 for IgG2b, respectively) and KLG4^m/m^ (n=6 for αCD8α and n=7 for IgG2b, respectively) mice treated as described in (A). (D) Representative images of HE staining (top) and statistics of tumor burdens and individual tumor sizes (bottom) of tumor-burdened lungs from the KL (n=7 for αCD8α and n=6 for IgG2b, respectively) and KLG4^m/m^ (n=6 for αCD8α and n=7 for IgG2b, respectively) mice treated as described in (A). Graphs show mean ± SEM (B, D). * *P* < 0.05, ** *P* < 0.01, *** *P* < 0.001, **** *P* <0.0001. ns: not significant (one-way ANOVA for B and D). Scale bars represent 5 mm (B and D). Data are representative results of two independent experiments (B-D).

**Figure S12.**
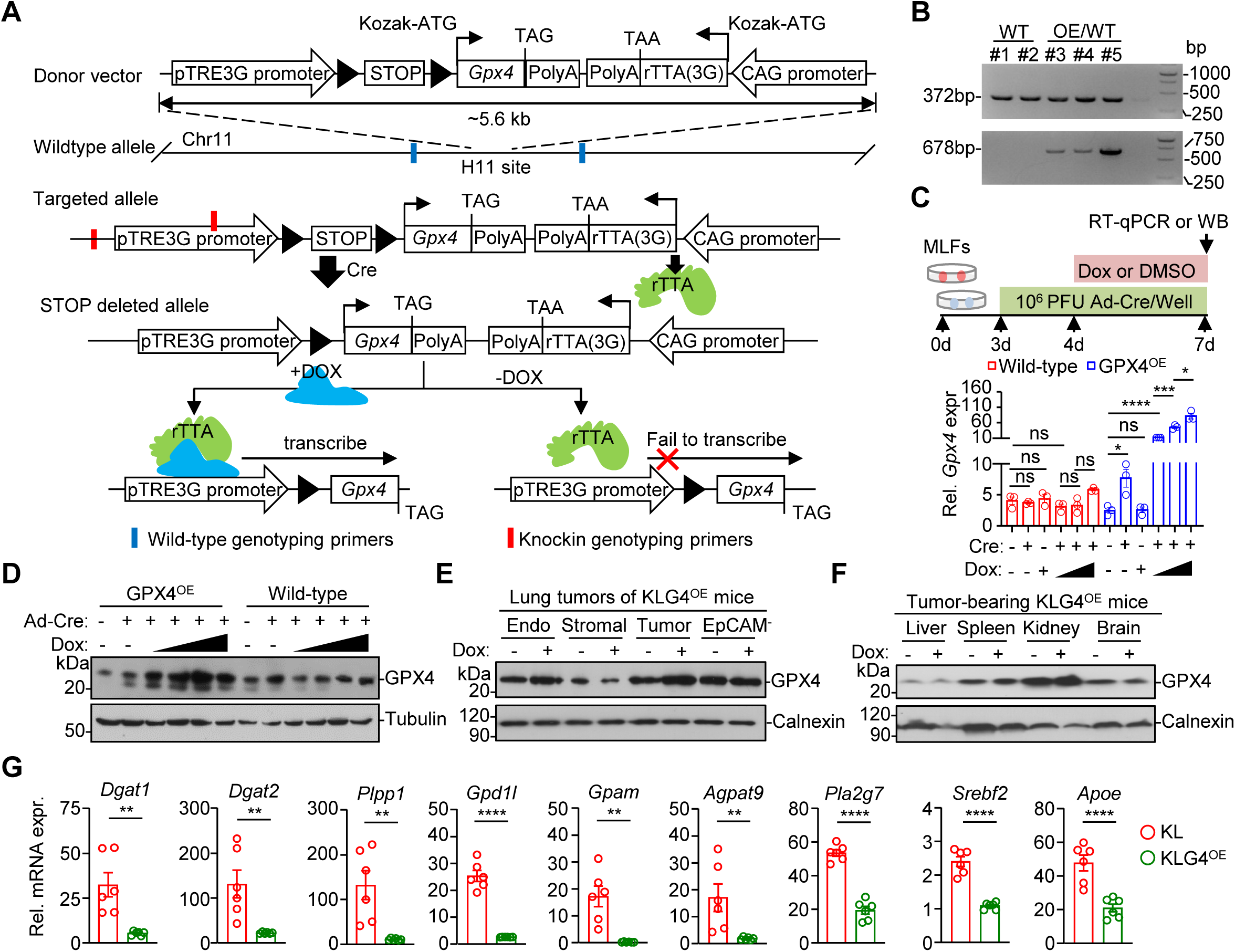
Generation and analysis of tumor cell-specific *Gpx4* overexpression mouse strain. (A) A scheme of the strategy for the generation of H11-pTRE3G-LSL-m*Gpx4*- CAG-rTTA (*Gpx4*^OE^) mice. (B) PCR analysis of the tail genomic DNAs from wild-type (Lane #1, #2) or *Gpx4*^OE^ (Lane #3-5) mice. (C, D) Experimental design (C, upper scheme), RT-qPCR (C, lower graph) (n = 3 biological replicates) or immunoblot analysis (D) of GPX4 in MLFs lysis from wild-type and *Gpx4*^m/m^ MLFs infected with Ad-Cre for 24 hours followed by treatment with Doxycycline (Dox, 20, 50, and 100 μg/mL) for 72 hours. (E, F) Immunoblot analysis of GPX4 in FACS-sorted endothelial cells, stromal cells, tumor cells or EpCAM^-^ cells in the tumors (E) or in liver, spleen, kidney or brain from KLG4^OE^ mice that were infected with Ad-Cre (2×10^6^ PFU per mouse) for 5 weeks followed by feeding of normal (-Dox) or Dox-supplemented (+Dox) chow food for 8 weeks. (G) RT-qPCR analysis of the signature genes involved in TAG synthesis in CD45^-^CD31^-^EpCAM^+^ tumor cells from lung tumors of KL (n = 6) and KLG4^OE^ (n = 6) mice treated as in (E). Graphs show mean ± SEM (C and G). * *P* < 0.05, ** *P* < 0.01, *** *P* < 0.001, **** *P* <0.0001. ns: not significant (one-way ANOVA for C and two-tailed student’s *t*-test for G). Data are representative results of two independent experiments (B-G).

**Figure S13.**
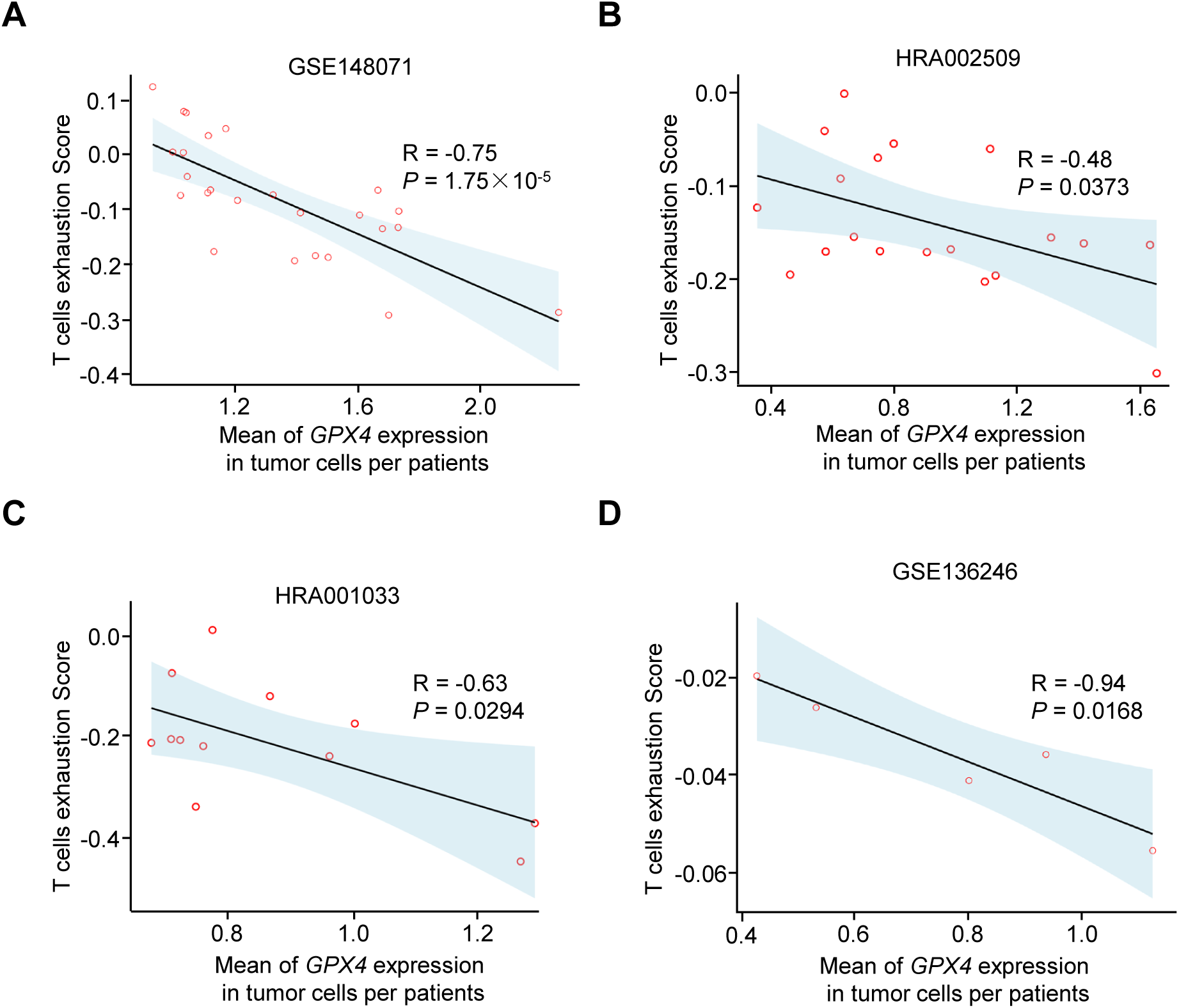
*GPX4* expression of tumor cells negatively correlates with T cells exhaustion score of T cells in NSCLC patients. (A-D) Correlation analysis between the mean *GPX4* expression in tumor cells and the exhaustion score of T cells across patients in the GSE148071 (A), HRA002509 (B), HRA001033 (C), and GSE136246 (D) datasets. Each dot represents one patient. The x-axis indicates the mean *GPX4* expression level in tumor cells from single-cell RNA sequencing data, while the y-axis represents the T cell exhaustion score of T cells from the same patient. R: Pearson correlation coefficient. *P* value were calculated using a two-tailed test based on the *t*-distribution.

**Figure S14.**
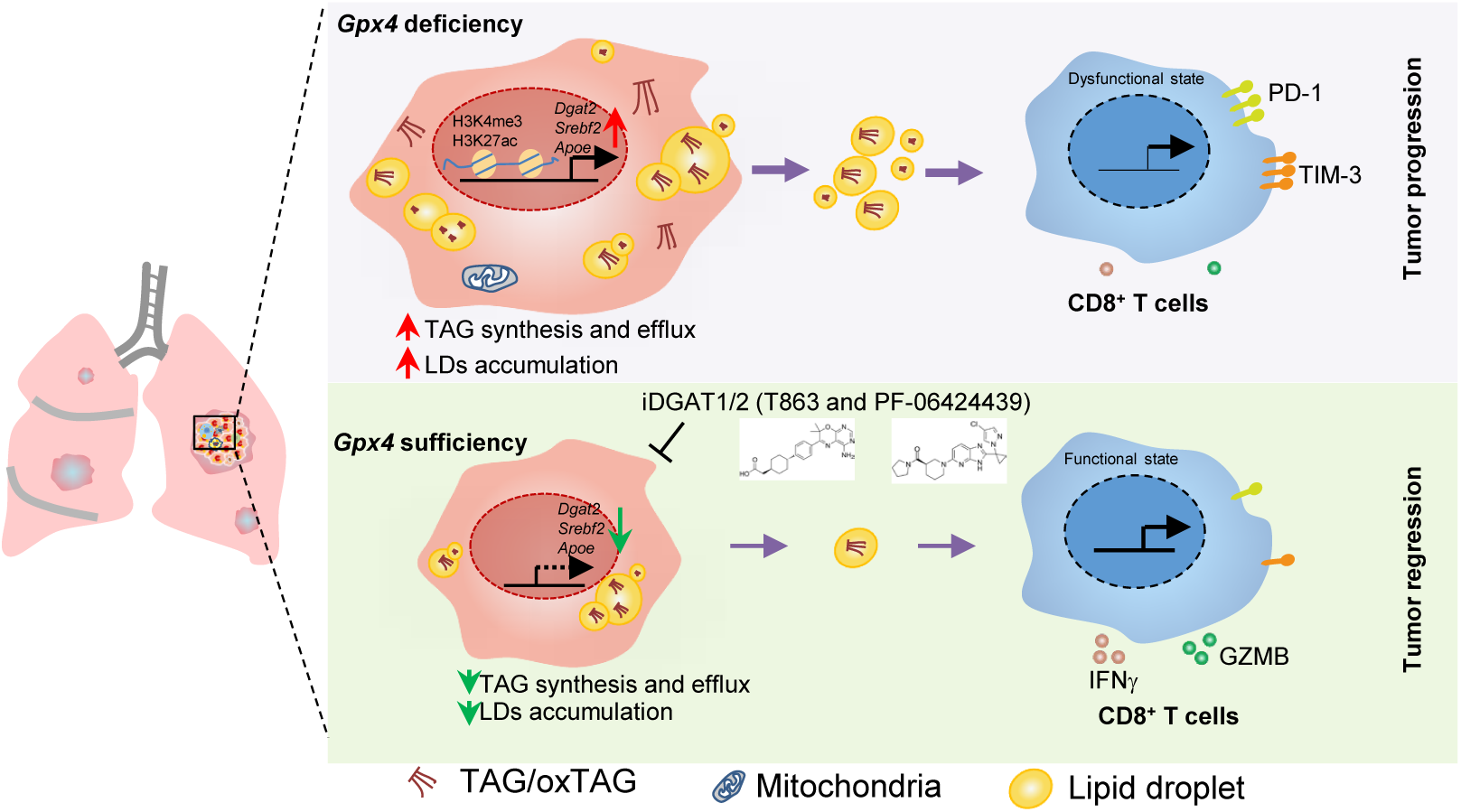
A model of metabolic crosstalk between *Gpx4*-deficient tumor cells and immune cells in the TME of autochthonous NSCLC. Inducible tumor cell-specific knockout of *Gpx4* reprograms the TAG metabolism and efflux to evade ferroptosis in tumor cells and induce dysfunction and exhaustion of CD8^+^ T in the TME was compromised and the progression of NSCLC is aggravated. Inhibition of TAG synthesis re-sensitizes tumor cells to ferroptosis and inhibits the progression of NSCLC. See the Discussion for a detailed description. TAG: Triacylglycerol; MHCI: major histocompatibility complex I; PLs: phospholipids; LDs: lipid droplets; *Dgat2*: Diacylglycerol acyltransferase 2; *Srebf2*: Sterol regulatory element binding transcription factor 2; *Apoe*: Apolipoprotein E.

## References

1. Ursini, F., Maiorino, M., Valente, M., Ferri, L., and Gregolin, C. (1982). Purification from pig liver of a protein which protects liposomes and biomembranes from peroxidative degradation and exhibits glutathione peroxidase activity on phosphatidylcholine hydroperoxides. Biochim Biophys Acta 710, 197–211. 10.1016/0005-2760(82)90150-3.

2. Cozza, G., Rossetto, M., Bosello-Travain, V., Maiorino, M., Roveri, A., Toppo, S., Zaccarin, M., Zennaro, L., and Ursini, F. (2017). Glutathione peroxidase 4-catalyzed reduction of lipid hydroperoxides in membranes: The polar head of membrane phospholipids binds the enzyme and addresses the fatty acid hydroperoxide group toward the redox center. Free Radic Biol Med 112, 1–11. 10.1016/j.freeradbiomed.2017.07.010.

3. Seiler, A., Schneider, M., Forster, H., Roth, S., Wirth, E.K., Culmsee, C., Plesnila, N., Kremmer, E., Radmark, O., Wurst, W., et al. (2008). Glutathione peroxidase 4 senses and translates oxidative stress into 12/15-lipoxygenase dependent- and AIF-mediated cell death. Cell Metab 8, 237–248. 10.1016/j.cmet.2008.07.005.

4. Matsushita, M., Freigang, S., Schneider, C., Conrad, M., Bornkamm, G.W., and Kopf, M. (2015). T cell lipid peroxidation induces ferroptosis and prevents immunity to infection. J Exp Med 212, 555–568. 10.1084/jem.20140857.

5. Canli, O., Alankus, Y.B., Grootjans, S., Vegi, N., Hultner, L., Hoppe, P.S., Schroeder, T., Vandenabeele, P., Bornkamm, G.W., and Greten, F.R. (2016). Glutathione peroxidase 4 prevents necroptosis in mouse erythroid precursors. Blood 127, 139–148. 10.1182/blood-2015-06-654194.

6. Hambright, W.S., Fonseca, R.S., Chen, L., Na, R., and Ran, Q. (2017). Ablation of ferroptosis regulator glutathione peroxidase 4 in forebrain neurons promotes cognitive impairment and neurodegeneration. Redox Biol 12, 8–17. 10.1016/j.redox.2017.01.021.

7. Muri, J., Thut, H., Bornkamm, G.W., and Kopf, M. (2019). B1 and Marginal Zone B Cells but Not Follicular B2 Cells Require Gpx4 to Prevent Lipid Peroxidation and Ferroptosis. Cell Rep 29, 2731–2744 e2734. 10.1016/j.celrep.2019.10.070.

8. Xu, C., Sun, S., Johnson, T., Qi, R., Zhang, S., Zhang, J., and Yang, K. (2021). The glutathione peroxidase Gpx4 prevents lipid peroxidation and ferroptosis to sustain Treg cell activation and suppression of antitumor immunity. Cell Rep 35, 109235. 10.1016/j.celrep.2021.109235.

9. Yao, Y., Chen, Z., Zhang, H., Chen, C., Zeng, M., Yunis, J., Wei, Y., Wan, Y., Wang, N., Zhou, M., et al. (2021). Selenium-GPX4 axis protects follicular helper T cells from ferroptosis. Nat Immunol 22, 1127–1139. 10.1038/s41590-021-00996-0.

10. Yant, L.J., Ran, Q., Rao, L., Van Remmen, H., Shibatani, T., Belter, J.G., Motta, L., Richardson, A., and Prolla, T.A. (2003). The selenoprotein GPX4 is essential for mouse development and protects from radiation and oxidative damage insults. Free Radic Biol Med 34, 496–502. 10.1016/s0891-5849(02)01360-6.

11. Yoo, S.E., Chen, L., Na, R., Liu, Y., Rios, C., Van Remmen, H., Richardson, A., and Ran, Q. (2012). Gpx4 ablation in adult mice results in a lethal phenotype accompanied by neuronal loss in brain. Free Radic Biol Med 52, 1820–1827. 10.1016/j.freeradbiomed.2012.02.043.

12. Friedmann Angeli, J.P., Schneider, M., Proneth, B., Tyurina, Y.Y., Tyurin, V.A., Hammond, V.J., Herbach, N., Aichler, M., Walch, A., Eggenhofer, E., et al. (2014). Inactivation of the ferroptosis regulator Gpx4 triggers acute renal failure in mice. Nat Cell Biol 16, 1180–1191. 10.1038/ncb3064.

13. Ran, Q., Liang, H., Gu, M., Qi, W., Walter, C.A., Roberts, L.J., 2nd, Herman, B., Richardson, A., and Van Remmen, H. (2004). Transgenic mice overexpressing glutathione peroxidase 4 are protected against oxidative stress-induced apoptosis. J Biol Chem 279, 55137-55146. 10.1074/jbc.M410387200.

14. Dixon, S.J., Lemberg, K.M., Lamprecht, M.R., Skouta, R., Zaitsev, E.M., Gleason, C.E., Patel, D.N., Bauer, A.J., Cantley, A.M., Yang, W.S., et al. (2012). Ferroptosis: an iron-dependent form of nonapoptotic cell death. Cell 149, 1060–1072. 10.1016/j.cell.2012.03.042.

15. Yang, W.S., SriRamaratnam, R., Welsch, M.E., Shimada, K., Skouta, R., Viswanathan, V.S., Cheah, J.H., Clemons, P.A., Shamji, A.F., Clish, C.B., et al. (2014). Regulation of ferroptotic cancer cell death by GPX4. Cell 156, 317–331. 10.1016/j.cell.2013.12.010.

16. Dai, E., Han, L., Liu, J., Xie, Y., Zeh, H.J., Kang, R., Bai, L., and Tang, D. (2020). Ferroptotic damage promotes pancreatic tumorigenesis through a TMEM173/STING-dependent DNA sensor pathway. Nat Commun 11, 6339. 10.1038/s41467-020-20154-8.

17. Herbst, R.S., Morgensztern, D., and Boshoff, C. (2018). The biology and management of non-small cell lung cancer. Nature 553, 446–454. 10.1038/nature25183.

18. Tang, R., Wang, H., and Tang, M. (2023). Roles of tissue-resident immune cells in immunotherapy of non-small cell lung cancer. Front Immunol 14, 1332814. 10.3389/fimmu.2023.1332814.

19. You, M., Xie, Z., Zhang, N., Zhang, Y., Xiao, D., Liu, S., Zhuang, W., Li, L., and Tao, Y. (2023). Signaling pathways in cancer metabolism: mechanisms and therapeutic targets. Signal Transduct Target Ther 8, 196. 10.1038/s41392-023-01442-3.

20. Wang, Y., Yan, Q., Fan, C., Mo, Y., Wang, Y., Li, X., Liao, Q., Guo, C., Li, G., Zeng, Z., et al. (2023). Overview and countermeasures of cancer burden in China. Sci China Life Sci 66, 2515–2526. 10.1007/s11427-022-2240-6.

21. Siegel, R.L., Miller, K.D., Wagle, N.S., and Jemal, A. (2023). Cancer statistics, 2023. CA Cancer J Clin 73, 17-48. 10.3322/caac.21763.

22. Gillette, M.A., Satpathy, S., Cao, S., Dhanasekaran, S.M., Vasaikar, S.V., Krug, K., Petralia, F., Li, Y., Liang, W.W., Reva, B., et al. (2020). Proteogenomic Characterization Reveals Therapeutic Vulnerabilities in Lung Adenocarcinoma. Cell 182, 200–225 e235. 10.1016/j.cell.2020.06.013.

23. Xu, J.Y., Zhang, C., Wang, X., Zhai, L., Ma, Y., Mao, Y., Qian, K., Sun, C., Liu, Z., Jiang, S., et al. (2020). Integrative Proteomic Characterization of Human Lung Adenocarcinoma. Cell 182, 245–261 e217. 10.1016/j.cell.2020.05.043.

24. Chen, Y.J., Roumeliotis, T.I., Chang, Y.H., Chen, C.T., Han, C.L., Lin, M.H., Chen, H.W., Chang, G.C., Chang, Y.L., Wu, C.T., et al. (2020). Proteogenomics of Non-smoking Lung Cancer in East Asia Delineates Molecular Signatures of Pathogenesis and Progression. Cell 182, 226–244 e217. 10.1016/j.cell.2020.06.012.

25. Skoulidis, F., and Heymach, J.V. (2019). Co-occurring genomic alterations in non-small-cell lung cancer biology and therapy. Nat Rev Cancer 19, 495–509. 10.1038/s41568-019-0179-8.

26. Johnson, L., Mercer, K., Greenbaum, D., Bronson, R.T., Crowley, D., Tuveson, D.A., and Jacks, T. (2001). Somatic activation of the K-ras oncogene causes early onset lung cancer in mice. Nature 410, 1111–1116. 10.1038/35074129.

27. Ji, H., Ramsey, M.R., Hayes, D.N., Fan, C., McNamara, K., Kozlowski, P., Torrice, C., Wu, M.C., Shimamura, T., Perera, S.A., et al. (2007). LKB1 modulates lung cancer differentiation and metastasis. Nature 448, 807–810. 10.1038/nature06030.

28. Chen, Z., Cheng, K., Walton, Z., Wang, Y., Ebi, H., Shimamura, T., Liu, Y., Tupper, T., Ouyang, J., Li, J., et al. (2012). A murine lung cancer co-clinical trial identifies genetic modifiers of therapeutic response. Nature 483, 613–617. 10.1038/nature10937.

29. Zhang, M., Yang, W., Wang, P., Deng, Y., Dong, Y.T., Liu, F.F., Huang, R., Zhang, P., Duan, Y.Q., Liu, X.D., et al. (2020). CCL7 recruits cDC1 to promote antitumor immunity and facilitate checkpoint immunotherapy to non-small cell lung cancer. Nat Commun 11, 6119. 10.1038/s41467-020-19973-6.

30. Dong, H.P., Li, Y., Tang, Z., Wang, P., Zhong, B., Chu, Q., and Lin, D. (2023). Combined targeting of CCL7 and Flt3L to promote the expansion and infiltration of cDC1s in tumors enhances T- cell activation and anti-PD-1 therapy effectiveness in NSCLC. Cell Mol Immunol 20, 850–853. 10.1038/s41423-023-00991-5.

31. Wang, P., Yang, W., Guo, H., Dong, H.P., Guo, Y.Y., Gan, H., Wang, Z., Cheng, Y., Deng, Y., Xie, S., et al. (2021). IL-36gamma and IL-36Ra Reciprocally Regulate NSCLC Progression by Modulating GSH Homeostasis and Oxidative Stress-Induced Cell Death. Adv Sci (Weinh) 8, e2101501. 10.1002/advs.202101501.

32. Sayin, V.I., Ibrahim, M.X., Larsson, E., Nilsson, J.A., Lindahl, P., and Bergo, M.O. (2014). Antioxidants accelerate lung cancer progression in mice. Sci Transl Med 6, 221ra215. 10.1126/scitranslmed.3007653.

33. Tang, Z., Hu, J., Li, X.C., Wang, W., Zhang, H.Y., Guo, Y.Y., Shuai, X., Chu, Q., Xie, C., Lin, D., and Zhong, B. (2024). A subset ofneutrophils activates anti-tumor immunity and inhibits non- small-cell lung cancer progression. Dev Cell. 10.1016/j.devcel.2024.10.010.

34. Li, F., Han, X., Li, F., Wang, R., Wang, H., Gao, Y., Wang, X., Fang, Z., Zhang, W., Yao, S., et al. (2015). LKB1 Inactivation Elicits a Redox Imbalance to Modulate Non-small Cell Lung Cancer Plasticity and Therapeutic Response. Cancer Cell 27, 698–711. 10.1016/j.ccell.2015.04.001.

35. Sutherland, K.D., Song, J.Y., Kwon, M.C., Proost, N., Zevenhoven, J., and Berns, A. (2014). Multiple cells-of-origin of mutant K-Ras-induced mouse lung adenocarcinoma. Proc Natl Acad Sci U S A 111, 4952–4957. 10.1073/pnas.1319963111.

36. Perl, A.K., Wert, S.E., Nagy, A., Lobe, C.G., and Whitsett, J.A. (2002). Early restriction of peripheral and proximal cell lineages during formation of the lung. Proc Natl Acad Sci U S A 99, 10482–10487. 10.1073/pnas.152238499.

37. Doll, S., Freitas, F.P., Shah, R., Aldrovandi, M., da Silva, M.C., Ingold, I., Goya Grocin, A., Xavier da Silva, T.N., Panzilius, E., Scheel, C.H., et al. (2019). FSP1 is a glutathione-independent ferroptosis suppressor. Nature 575, 693–698. 10.1038/s41586-019-1707-0.

38. Li, Y., Ran, Q., Duan, Q., Jin, J., Wang, Y., Yu, L., Wang, C., Zhu, Z., Chen, X., Weng, L., et al. (2024). 7-Dehydrocholesterol dictates ferroptosis sensitivity. Nature 626, 411–418. 10.1038/s41586-023-06983-9.

39. Kim, R., Hashimoto, A., Markosyan, N., Tyurin, V.A., Tyurina, Y.Y., Kar, G., Fu, S., Sehgal, M., Garcia-Gerique, L., Kossenkov, A., et al. (2022). Ferroptosis of tumour neutrophils causes immune suppression in cancer. Nature 612, 338–346. 10.1038/s41586-022-05443-0.

40. Olzmann, J.A., and Carvalho, P. (2019). Dynamics and functions of lipid droplets. Nat Rev Mol Cell Biol 20, 137–155. 10.1038/s41580-018-0085-z.

41. Mathiowetz, A.J., and Olzmann, J.A. (2024). Lipid droplets and cellular lipid flux. Nat Cell Biol 26, 331–345. 10.1038/s41556-024-01364-4.

42. Bailey, A.P., Koster, G., Guillermier, C., Hirst, E.M., MacRae, J.I., Lechene, C.P., Postle, A.D., and Gould, A.P. (2015). Antioxidant Role for Lipid Droplets in a Stem Cell Niche of Drosophila. Cell 163, 340–353. 10.1016/j.cell.2015.09.020.

43. Lee, H., Horbath, A., Kondiparthi, L., Meena, J.K., Lei, G., Dasgupta, S., Liu, X., Zhuang, L., Koppula, P., Li, M., et al. (2024). Cell cycle arrest induces lipid droplet formation and confers ferroptosis resistance. Nat Commun 15, 79. 10.1038/s41467-023-44412-7.

44. Krahmer, N., Guo, Y., Farese, R.V., Jr., and Walther, T.C. (2009). SnapShot: Lipid Droplets. Cell 139, 1024–1024 e1021. 10.1016/j.cell.2009.11.023.

45. Hu, J., Zhang, L., Xia, H., Yan, Y., Zhu, X., Sun, F., Sun, L., Li, S., Li, D., Wang, J., et al. (2023). Tumor microenvironment remodeling after neoadjuvant immunotherapy in non-small cell lung cancer revealed by single-cell RNA sequencing. Genome Med 15, 14. 10.1186/s13073-023-01164-9.

46. Farese, R.V., Jr., and Walther, T.C. (2023). Glycerolipid Synthesis and Lipid Droplet Formation in the Endoplasmic Reticulum. Cold Spring Harb Perspect Biol 15. 10.1101/cshperspect.a041246.

47. Wang, Y., Chen, W., Qiao, S., Zou, H., Yu, X.J., Yang, Y., Li, Z., Wang, J., Chen, M.S., Xu, J., and Zheng, L. (2024). Lipid droplet accumulation mediates macrophage survival and Treg recruitment via the CCL20/CCR6 axis in human hepatocellular carcinoma. Cell Mol Immunol 21, 1120–1130. 10.1038/s41423-024-01199-x.

48. Mensenkamp, A.R., Van Luyn, M.J., Havinga, R., Teusink, B., Waterman, I.J., Mann, C.J., Elzinga, B.M., Verkade, H.J., Zammit, V.A., Havekes, L.M., et al. (2004). The transport of triglycerides through the secretory pathway of hepatocytes is impaired in apolipoprotein E deficient mice. J Hepatol 40, 599–606. 10.1016/j.jhep.2003.12.011.

49. Getz, G.S., and Reardon, C.A. (2009). Apoprotein E as a lipid transport and signaling protein in the blood, liver, and artery wall. J Lipid Res 50 *Suppl*, S156-161. 10.1194/jlr.R800058-JLR200.

50. Doll, S., Proneth, B., Tyurina, Y.Y., Panzilius, E., Kobayashi, S., Ingold, I., Irmler, M., Beckers, J., Aichler, M., Walch, A., et al. (2017). ACSL4 dictates ferroptosis sensitivity by shaping cellular lipid composition. Nat Chem Biol 13, 91–98. 10.1038/nchembio.2239.

51. Qiu, B., Zandkarimi, F., Bezjian, C.T., Reznik, E., Soni, R.K., Gu, W., Jiang, X., and Stockwell, B.R. (2024). Phospholipids with two polyunsaturated fatty acyl tails promote ferroptosis. Cell 187, 1177–1190 e1118. 10.1016/j.cell.2024.01.030.

52. Liao, P., Wang, W., Wang, W., Kryczek, I., Li, X., Bian, Y., Sell, A., Wei, S., Grove, S., Johnson, J.K., et al. (2022). CD8(+) T cells and fatty acids orchestrate tumor ferroptosis and immunity via ACSL4. Cancer Cell 40, 365–378 e366. 10.1016/j.ccell.2022.02.003.

53. Xu, S., Chaudhary, O., Rodriguez-Morales, P., Sun, X., Chen, D., Zappasodi, R., Xu, Z., Pinto, A.F.M., Williams, A., Schulze, I., et al. (2021). Uptake of oxidized lipids by the scavenger receptor CD36 promotes lipid peroxidation and dysfunction in CD8(+) T cells in tumors. Immunity 54, 1561–1577 e1567. 10.1016/j.immuni.2021.05.003.

54. Ma, X., Xiao, L., Liu, L., Ye, L., Su, P., Bi, E., Wang, Q., Yang, M., Qian, J., and Yi, Q. (2021). CD36-mediated ferroptosis dampens intratumoral CD8(+) T cell effector function and impairs their antitumor ability. Cell Metab 33, 1001–1012 e1005. 10.1016/j.cmet.2021.02.015.

55. Wu, J.E., Manne, S., Ngiow, S.F., Baxter, A.E., Huang, H., Freilich, E., Clark, M.L., Lee, J.H., Chen, Z., Khan, O., et al. (2023). In vitro modeling of CD8(+) T cell exhaustion enables CRISPR screening to reveal a role for BHLHE40. Sci Immunol 8, eade3369. 10.1126/sciimmunol.ade3369.

56. Huang, Q., Wu, X., Wang, Z., Chen, X., Wang, L., Lu, Y., Xiong, D., Liu, Q., Tian, Y., Lin, H., et al. (2022). The primordial differentiation of tumor-specific memory CD8(+) T cells as bona fide responders to PD-1/PD-L1 blockade in draining lymph nodes. Cell 185, 4049–4066 e4025. 10.1016/j.cell.2022.09.020.

57. Wu, F., Fan, J., He, Y., Xiong, A., Yu, J., Li, Y., Zhang, Y., Zhao, W., Zhou, F., Li, W., et al. (2021). Single-cell profiling of tumor heterogeneity and the microenvironment in advanced non-small cell lung cancer. Nat Commun 12, 2540. 10.1038/s41467-021-22801-0.

58. Yan, Y., Sun, D., Hu, J., Chen, Y., Sun, L., Yu, H., Xiong, Y., Huang, Z., Xia, H., Zhu, X., et al. (2025). Multi-omic profiling highlights factors associated with resistance to immuno- chemotherapy in non-small-cell lung cancer. Nat Genet 57, 126–139. 10.1038/s41588-024-01998-y.

59. Maroni, G., Bassal, M.A., Krishnan, I., Fhu, C.W., Savova, V., Zilionis, R., Maymi, V.A., Pandell, N., Csizmadia, E., Zhang, J., et al. (2021). Identification of a targetable KRAS-mutant epithelial population in non-small cell lung cancer. Commun Biol 4, 370. 10.1038/s42003-021-01897-6.

60. Ping, Y., Shan, J., Qin, H., Li, F., Qu, J., Guo, R., Han, D., Jing, W., Liu, Y., Liu, J., et al. (2024). PD- 1 signaling limits expression of phospholipid phosphatase 1 and promotes intratumoral CD8(+) T cell ferroptosis. Immunity 57, 2122–2139 e2129. 10.1016/j.immuni.2024.08.003.

61. Wu, K., Vaughan, A.J., Bossowski, J.P., Hao, Y., Ziogou, A., Kim, S.M., Kim, T.H., Nakamura, M.N., Pillai, R., Mancini, M., et al. (2025). Targeting FSP1 triggers ferroptosis in lung cancer. Nature. 10.1038/s41586-025-09710-8.

62. Jackson, E.L., Willis, N., Mercer, K., Bronson, R.T., Crowley, D., Montoya, R., Jacks, T., and Tuveson, D.A. (2001). Analysis of lung tumor initiation and progression using conditional expression of oncogenic K-ras. Genes Dev 15, 3243–3248. 10.1101/gad.943001.

63. Zilionis, R., Engblom, C., Pfirschke, C., Savova, V., Zemmour, D., Saatcioglu, H.D., Krishnan, I., Maroni, G., Meyerovitz, C.V., Kerwin, C.M., et al. (2019). Single-Cell Transcriptomics of Human and Mouse Lung Cancers Reveals Conserved Myeloid Populations across Individuals and Species. Immunity 50, 1317–1334 e1310. 10.1016/j.immuni.2019.03.009.

64. Camolotto, S.A., Pattabiraman, S., Mosbruger, T.L., Jones, A., Belova, V.K., Orstad, G., Streiff, M., Salmond, L., Stubben, C., Kaestner, K.H., and Snyder, E.L. (2018). FoxA1 and FoxA2 drive gastric differentiation and suppress squamous identity in NKX2-1-negative lung cancer. Elife 7. 10.7554/eLife.38579.

65. Snyder, E.L., Watanabe, H., Magendantz, M., Hoersch, S., Chen, T.A., Wang, D.G., Crowley, D., Whittaker, C.A., Meyerson, M., Kimura, S., and Jacks, T. (2013). Nkx2-1 represses a latent gastric differentiation program in lung adenocarcinoma. Mol Cell 50, 185–199. 10.1016/j.molcel.2013.02.018.

66. Minoo, P., Hu, L., Xing, Y., Zhu, N.L., Chen, H., Li, M., Borok, Z., and Li, C. (2007). Physical and functional interactions between homeodomain NKX2.1 and winged helix/forkhead FOXA1 in lung epithelial cells. Mol Cell Biol 27, 2155–2165. 10.1128/MCB.01133-06.

67. Moya, M., Benet, M., Guzman, C., Tolosa, L., Garcia-Monzon, C., Pareja, E., Castell, J.V., and Jover, R. (2012). Foxa1 reduces lipid accumulation in human hepatocytes and is down-regulated in nonalcoholic fatty liver. PLoS One 7, e30014. 10.1371/journal.pone.0030014.

68. Wang, H., Franco, F., Tsui, Y.C., Xie, X., Trefny, M.P., Zappasodi, R., Mohmood, S.R., Fernandez-Garcia, J., Tsai, C.H., Schulze, I., et al. (2020). CD36-mediated metabolic adaptation supports regulatory T cell survival and function in tumors. Nat Immunol 21, 298–308. 10.1038/s41590-019-0589-5.

69. Calle, R.A., Amin, N.B., Carvajal-Gonzalez, S., Ross, T.T., Bergman, A., Aggarwal, S., Crowley, C., Rinaldi, A., Mancuso, J., Aggarwal, N., et al. (2021). ACC inhibitor alone or co-administered with a DGAT2 inhibitor in patients with non-alcoholic fatty liver disease: two parallel, placebo-controlled, randomized phase 2a trials. Nat Med 27, 1836–1848. 10.1038/s41591-021-01489-1.

70. Yang, W., Dong, H.P., Wang, P., Xu, Z.G., Xian, J., Chen, J., Wu, H., Lou, Y., Lin, D., and Zhong, B. (2022). IL-36gamma and IL-36Ra Reciprocally Regulate Colon Inflammation and Tumorigenesis by Modulating the Cell-Matrix Adhesion Network and Wnt Signaling. Adv Sci (Weinh) 9, e2103035. 10.1002/advs.202103035.

71. Chen, S., Cui, W., Chi, Z., Xiao, Q., Hu, T., Ye, Q., Zhu, K., Yu, W., Wang, Z., Yu, C., et al. (2022). Tumor-associated macrophages are shaped by intratumoral high potassium via Kir2.1. Cell Metab 34, 1843–1859 e1811. 10.1016/j.cmet.2022.08.016.

72. Eil, R., Vodnala, S.K., Clever, D., Klebanoff, C.A., Sukumar, M., Pan, J.H., Palmer, D.C., Gros, A., Yamamoto, T.N., Patel, S.J., et al. (2016). Ionic immune suppression within the tumour microenvironment limits T cell effector function. Nature 537, 539–543. 10.1038/nature19364.

73. Chapman, M.J., Goldstein, S., Lagrange, D., and Laplaud, P.M. (1981). A density gradient ultracentrifugal procedure for the isolation of the major lipoprotein classes from human serum. J Lipid Res 22, 339–358.

74. Yin, H., Cox, B.E., Liu, W., Porter, N.A., Morrow, J.D., and Milne, G.L. (2009). Identification of intact oxidation products of glycerophospholipids in vitro and in vivo using negative ion electrospray iontrap mass spectrometry. J Mass Spectrom 44, 672–680. 10.1002/jms.1542.

75. Paynter, N.P., Balasubramanian, R., Giulianini, F., Wang, D.D., Tinker, L.F., Gopal, S., Deik, A.A., Bullock, K., Pierce, K.A., Scott, J., et al. (2018). Metabolic Predictors of Incident Coronary Heart Disease in Women. Circulation 137, 841–853. 10.1161/CIRCULATIONAHA.117.029468.

76. Wu, T., Hu, E., Xu, S., Chen, M., Guo, P., Dai, Z., Feng, T., Zhou, L., Tang, W., Zhan, L., et al. (2021). clusterProfiler 4.0: A universal enrichment tool for interpreting omics data. Innovation (Camb) 2, 100141. 10.1016/j.xinn.2021.100141.

77. Subramanian, A., Tamayo, P., Mootha, V.K., Mukherjee, S., Ebert, B.L., Gillette, M.A., Paulovich, A., Pomeroy, S.L., Golub, T.R., Lander, E.S., and Mesirov, J.P. (2005). Gene set enrichment analysis: a knowledge-based approach for interpreting genome-wide expression profiles. Proc Natl Acad Sci U S A 102, 15545–15550. 10.1073/pnas.0506580102.

